# Coordination of distinct sources of excitatory inputs enhances motion selectivity in the mouse visual thalamus

**DOI:** 10.1101/2025.01.08.631826

**Authors:** Yue Fei, Michelle Luh, Ashley Ontiri, Dawood Ghauri, Wenxin Hu, Liang Liang

## Abstract

Multiple sources innervate the visual thalamus to influence image-forming vision prior to the cortex, yet it remains unclear how non-retinal and retinal input coordinate to shape thalamic visual selectivity. Using dual-color two-photon calcium imaging in the thalamus of awake mice, we observed similar coarse-scale retinotopic organization between axons of superior colliculus neurons and retinal ganglion cells, both providing strong converging excitatory input to thalamic neurons. At a fine scale of ∼10 µm, collicular boutons often shared visual feature preferences with nearby retinal boutons. Inhibiting collicular input significantly suppressed visual responses in thalamic neurons and specifically reduced motion selectivity in neurons preferring nasal-to-temporal motion. The reduction in motion selectivity could be the result of silencing sharply tuned direction-selective colliculogeniculate input. These findings suggest that the thalamus is not merely a relay but selectively integrates inputs from multiple regions to build stimulus selectivity and shape the information transmitted to the cortex.

**HIGHLIGHTS:**

Chronic dual-color calcium imaging reveals diverse visual tuning of collicular axonal boutons.
Nearby collicular and retinal boutons often share feature preferences at ∼10 µm scale
Silencing of collicular input suppresses visual responses in the majority of thalamic neurons.
Silencing of collicular input reduces motion selectivity in thalamic neurons.

## INTRODUCTION

In typical neural circuits, presynaptic axons from distinct regions converge to drive information processing in postsynaptic neurons. In the visual system, such multi-region convergence happens as early as the primary visual thalamus: the dorsal lateral geniculate nucleus (dLGN). The dLGN is traditionally viewed as a ‘relay’ nucleus, routing signals from the eye to the primary visual cortex^1,2^. However, beyond retinal input, the dLGN also receives projections from several other brain regions, including the visual cortex, the brain stem, the superior colliculus (SC), and other nuclei from the thalamus, pretectum, and midbrain^2–13^. Indeed, non-retinal inputs account for ∼90% of the total input to the dLGN^14^. Many questions remain about these non-retinal inputs: What visual information do they convey to the dLGN? How do they collaborate with retinal input in their organization and function? Does the dLGN act principally as a relay for retinal input, or does it integrate non-retinal inputs to build stimulus selectivity? Given the dLGN’s crucial role in conveying visual information to the cortex, the selective regulation of information through integrating retinal and non-retinal inputs within the dLGN could significantly shape the information flow and influence cortical computation.

Among non-retinal inputs, projections from the colliculus to the geniculate (colliculogeniculate) exhibit several distinct characteristics. These projections have been observed in all mammals studied to date, including rodents, felines, and non-human primates, and have been shown to innervate homologous regions in the dLGN across species^6,15–17^. Notably, colliculogeniculate axons resemble retinogeniculate axons in several synaptic properties: Collicular axonal boutons synapse on proximal dendrites of dLGN neurons and can converge with retinal boutons on the same dendrites^18^. Moreover, most colliculogeniculate axons are glutamatergic and can provide ‘driver-like’ excitation to their target dLGN neurons when tested in brain slices^18^. However, the *in vivo* implications of ‘driver-like’ excitation from the SC for the dLGN remain elusive.

Although both the dLGN and SC are major recipients of retinal input, they have long been thought to contribute to distinct visual functions. While the dLGN is the gateway to the cortex responsible for image-forming vision, the SC appears to mediate rapid, reflexive responses for non-conscious visual processing^19–34^. In addition to transmitting visual signals to the deeper SC, the superficial SC (sSC) also sends multiple projections to the thalamus. The sSC projection to the higher-order visual thalamus is implicated in blindsight^35–39^, and the GABAergic SC projection to the LGN (especially the ventral LGN [vLGN]) regulates defensive behaviors to visual threat^31^. However, the presence of the highly conserved, strong excitatory sSC-dLGN projection suggests that the SC may also actively participate in the retino-geniculo-cortical pathway by directly influencing visual computation in the dLGN. It also indicates that early visual centers may work more closely together for processing conscious vision than previously thought.

In addition to neurons with classical center-surround receptive fields, neurons in the mouse dLGN shell region also exhibit higher-order visual feature selectivity, such as direction and orientation selectivity^40–44^. Feature selectivity in dLGN neurons has been thought to depend mainly on retinal input, with cortical feedback modulating response magnitudes rather than affecting feature preferences or selectivity^41,43–47^. Interestingly, excitatory neurons in the superficial lamina of the stratum griseum superficiale (SGS) in the SC are highly direction-selective^48–53^. Moreover, colliculogeniculate projections originate primarily from the upper SGS and target the dLGN shell^6,54,55^. This raises the possibility that dLGN-projecting SC neurons exhibit higher-order feature selectivity, and these ‘driver-like’ excitatory inputs from the SC may shape receptive field properties in dLGN neurons.

The functional organization of axons is foundational to their influence on target neurons. While the anatomical mapping of excitatory inputs from distinct brain regions has started^56^, their fine-scale functional arrangement remains unresolved. Neurons in the dLGN and SC follow retinotopic organization^27,42^, but how precisely colliculogeniculate projections are organized by retinotopy remains unclear. Axonal boutons from multiple retinal ganglion cells (RGCs) cluster on proximal dendrites of dLGN neurons^57,58^, and within ∼10 μm, nearby retinal boutons from distinct axons often share visual feature preferences^59^. We sought to determine whether similar fine-scale functional logic guides the convergence between retinal and collicular axonal boutons. Such selective convergence could reinforce shared visual information in target neurons, allowing collicular boutons to act in concert with retinal boutons at a subcellular level to impact feature coding in target dLGN neurons.

To investigate visual properties and functional convergence between retinal and collicular inputs, we developed a dual-color chronic two-photon imaging platform for simultaneously recording calcium activity in hundreds of individual retinal and collicular boutons within the same dLGN shell region of awake, head-restrained mice. Additionally, we combined chemogenetic silencing of colliculogeniculate inputs with functional imaging of dLGN neurons to measure the causal effects of collicular input on visual response magnitudes and properties in different dLGN neuron types. By linking the visual responses and fine-scale functional organization of input axons with changes in dLGN cellular responses following activity manipulation, our findings reveal how distinct streams of excitatory input coordinate at multiple levels in the dLGN to enhance specific motion signals and shape motion selectivity as visual information is routed to the cortex.

## RESULTS

### Dual-color Calcium Imaging Revealed Diverse Visual Responses in Both Retinal and Collicular Axons in the Dorsal dLGN

To characterize the visual response properties of colliculogeniculate axons and compare them with retinogeniculate axons targeting the same dLGN region, we conducted chronic dual-color two-photon calcium imaging through a cranial window positioned above the dLGN of awake, head-restrained mice (Figure 1A). To selectively label the SC neurons that project to the dLGN over the lateral posterior nucleus (LP) of the thalamus, we used Rorβ-Cre mice^11,54,55,60^. Most colliculogeniculate neurons (∼95%) are non-GABAergic, and Rorβ-Cre-positive colliculogeniculate axons were reported to provide only excitatory input to dLGN neurons^18,54^. In Rorβ-Cre mice, we injected AAVs carrying green fluorescent calcium indicator GCaMP6f into the contralateral eye to label retinogeniculate axons, and AAVs carrying Cre-dependent red fluorescent calcium indicator jRGECO1a into the ipsilateral superficial SC to label colliculogeniculate axons (Figure 1B; see also Figure 3C)^61,62^. We simultaneously recorded the calcium activity of hundreds of retinogeniculate and colliculogeniculate boutons in the upper 50-120 µm of the dLGN by imaging GCaMP6f and jRGECO1a simultaneously (Figure 1C). To minimize bleed-through signals between the green and red fluorescence channels, we implemented a color de- mixing method (Figure S1A-S1D; see STAR Methods).

**Figure 1.**
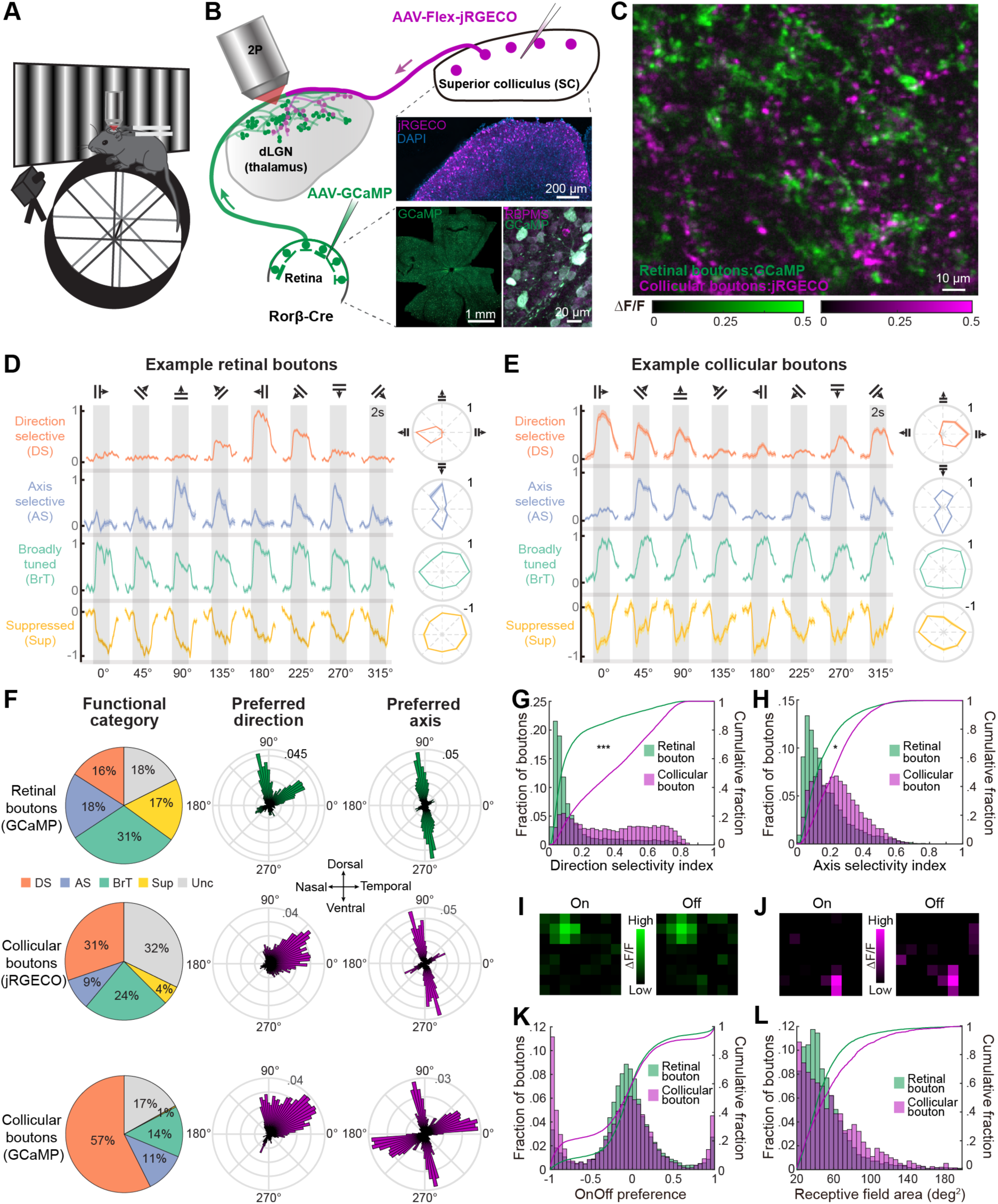
Dual-color Two-photon Calcium Imaging Reveals Visual Response Properties of Retinal and Collicular Boutons in the Same dLGN Region in Awake Mice. (A) Schematic of imaging setup in awake, head-restrained mice. (B) Left and top: Schematic of dual-color two-photon calcium imaging. Middle: jRGECO1a-labeled neurons in the SC of a Rorβ-Cre mouse. Bottom left: GCaMP6f-labeled RGCs in a retina. Bottom right: co-staining of RBPMS (magenta) and GCaMP6f (green) in RGCs. (C) Visually evoked fluorescence changes in retinal and collicular boutons expressing GCaMP6f and jRGECO1a, respectively, in an example field of view (FOV) in the dLGN. ΔF/F, the fractional change in fluorescence. (D) Retinal boutons were classified into functional categories based on their responses to a 2-s presentation of drifting gratings (gray bars). Left: Example normalized response time courses of an example retinal bouton in each category (mean ± SEM). Right: Normalized response tuning curves (mean ± SEM). (E) Similar to D but in example collicular boutons. (F) Left: Distribution of boutons in each functional category (retinal boutons vs. jRGECO collicular boutons or retinal boutons vs. GCaMP collicular boutons: p < 0.001, Chi-squared test; p < 0.0001 for each category, *post hoc* Pearson residuals test). Middle: Polar distribution of the preferred motion direction for direction-selective boutons. Right: Polar distribution of the preferred motion axis for axis-selective boutons. Top, middle, bottom: Distribution of GCaMP6f-labeled retinal boutons, jRGECO1a-labeled collicular boutons, and GCaMP6f-labeled collicular boutons. DS, direction-selective (n = 2563 retinal boutons, 3079 jRGECO1a collicular boutons, 6456 GCaMP6f collicular boutons); AS, axis-selective (n = 3271 retinal boutons, 911 jRGECO1a collicular boutons, 1268 GCaMP6f collicular boutons); BrT, broadly-tuned (n = 5485 retinal boutons, 2465 jRGECO1a collicular boutons, 1568 GCaMP6f collicular boutons); Sup, suppressed (n = 3100 retinal boutons, 553 jRGECO1a collicular boutons, 39 GCaMP6f collicular boutons); Unc, responsive but not classified (n = 3139 retinal boutons, 3375 jRGECO1a collicular boutons, 1896 GCaMP6f collicular boutons). Retinal boutons and jRGECO1a-labeled collicular boutons were from the same 18 FOVs, 8 mice; GCaMP6f-labeled collicular boutons were from separately recorded 10 FOVs, 5 mice. (G) Distributions of the direction selectivity index (DSI) across retinal boutons and across GCaMP6f-labeled collicular boutons were significantly different (p < 10^-3^). The cumulative distributions of the DSI are overlaid. (H) Distributions of motion axis selectivity index (ASI) were significantly different between retinal and collicular boutons (p = 0.02). (I) The On and Off receptive fields for a retinal bouton mapped by sparse flashing squares. (J) Similar to I but for a collicular bouton. (K) Distributions of the On/Off preference for retinal and collicular boutons (p = 0.003). (L) No significant difference in receptive field areas between retinal and collicular boutons (p = 0.74). G, H, K, and L: n = 11319 retinal boutons, 18 FOVs, 8 mice; n = 9292 collicular boutons, 10 FOVs, 5 mice; linear mixed-effects model. See also Figures S1, S2 and S9.

We observed functional diversity in both retinogeniculate and colliculogeniculate axons innervating the same dorsal dLGN region. To compare the receptive field properties of retinogeniculate and colliculogeniculate axons, we delivered a battery of visual stimuli, including full-field drifting gratings of 8 different directions, 3 spatial frequencies and 3 temporal frequencies and sparse 5° flashing white or black squares (see STAR Methods). Based on the responses to drifting gratings, we classified axonal boutons into different functional categories: direction-selective (DS; preferentially responsive to one direction of motion), axis-selective (AS; preferentially responsive to opposite directions of motion along the same axis), broadly-tuned (BrT; evenly responsive across all directions of motion), and suppressed (Sup; suppressed during visual stimulation)^59^. We found boutons of each functional category in both retinogeniculate and colliculogeniculate axons (Figure 1D and 1E).

### Comparison of Visual Response Properties Between Retinal and Collicular Axons

However, the distribution of boutons across categories differed between the two distinct sources of input (p < 0.001, Chi-squared test). Strikingly, the fraction of jRGECO1a-labeled DS colliculogeniculate boutons almost doubled that of retinogeniculate boutons, and a very limited fraction of colliculogeniculate boutons was suppressed by drifting gratings. We confirmed that the enrichment of DS boutons was not due to the use of different calcium indicators (GCaMP6f vs. jRGECO1a), as an even larger fraction of DS boutons were observed among GCaMP6f-labeled colliculogeniculate axons in a separate cohort of mice (Figure 1F bottom row; comparisons of other visual receptive properties between GCaMP6f- and jRGECO1a-labeled collicular boutons are shown in Figure S1E-S1H). Moreover, compared to the retinogeniculate boutons, the DS colliculogeniculate boutons disproportionally preferred the nasal-to-temporal direction of motion (p < 0.001, Chi-squared test with *post hoc* adjusted Pearson residuals test), and the AS colliculogeniculate boutons had a larger fraction of boutons preferring the horizontal motion axis (Figure 1F; p < 0.001, Chi-squared test with *post hoc* adjusted Pearson residuals test). Notably, when we compared the direction selectivity indices (DSI) and axis selectivity indices (ASI) between retinogeniculate and colliculogeniculate bouton populations that both expressed GCaMP6f, we found colliculogeniculate boutons exhibited sharper tuning (Figure 1G-1H).

Retinogeniculate and colliculogeniculate boutons also exhibited diverse preferences for other visual features, such as luminance changes (On/Off preference) and the spatial frequency (SF) and temporal frequency (TF) of drifting gratings. Both retinal and collicular boutons displayed multimodal distributions for On/Off preferences, with peaks centered around Off, On-Off, and On (corresponding to On/Off indices of -1, 0, and 1, respectively) (Figure 1I-1K). Compared to retinal boutons, larger fractions of collicular boutons were tuned to the middle SF and TF (Figure S2E and S2F). Both input sources exhibited similar distributions in receptive field sizes, with a median diameter of ∼8° (Figure 1L).

Furthermore, we calculated the coefficient of variance to quantify the trial-to-trial response variability in retinogeniculate boutons, dLGN neurons, and colliculogeniculate boutons. Retinal boutons were the least variable, dLGN neurons showed the highest variability, and collicular boutons fell in between (Figure S1I), aligning with previous reports of increasing variability along the visual hierarchy from the retina to the V1^63,64^. This result suggests that the SC sends post-processed information to the dLGN, rather than merely duplicating retinal inputs.

We also analyzed the distributions of receptive field properties based on individual axons, not just boutons, to avoid bias from axons with many boutons. Boutons were assigned to the same axon if they exhibited high correlations in spontaneous activity during blank trials with uniform luminance, using a hierarchical clustering method previously established for retinogeniculate axons^59^. As illustrated in the example field of view (FOV), boutons from different axons exhibited low correlation even when in close proximity, whereas boutons from the same axon maintained high correlation despite being more than 100 µm apart (Figure S2A). We confirmed the axon assignment of collicular boutons by the highly similar visual responses among boutons classified into the same axon (Figure S2B) and by the well-separated distributions of the intra-axon and inter-axon correlation of spontaneous activity and visually evoked activity (Figure S2C and S2D). We found that the distributions of functional categories and visual feature preferences were highly consistent whether analyzed by boutons or axons (Figure S2E and S2F).

Together, these data demonstrate that the colliculogeniculate and retinogeniculate axons both convey diverse channels of visual information to the dLGN and exhibit large overlap in their population distributions across multiple visual receptive field properties. However, they differ in several ways: collicular input consists of a much larger fraction of direction-selective axons, exhibits higher motion selectivity, and a larger proportion of direction-/axis-selective collicular axons are tuned to motion along the horizontal axis.

### Highly Consistent Retinotopic Organization between the Retinogeniculate Input and the Colliculogeniculate Input

While our previous work demonstrated that retinal boutons are organized according to retinotopy on a coarse spatial scale with an accuracy of approximately 20 μm^59^, the precision of the retinotopic organization among collicular boutons and the retinotopic alignment between collicular and retinal boutons remained unknown. To measure the retinotopic preferences of individual retinal and collicular boutons innervating the same dLGN region, we presented 5° x 40° horizontal or vertical bars containing spatiotemporal noise patterns at several locations along the azimuth and vertical axes in visual space (see STAR Methods). We observed highly consistent coarse-scale retinotopic maps between the retinal and collicular input (Figure 2A). A magnified image showed that nearby retinal and collicular boutons exhibited peak responses at similar elevations or azimuth locations (Figure 2B).

**Figure 2.**
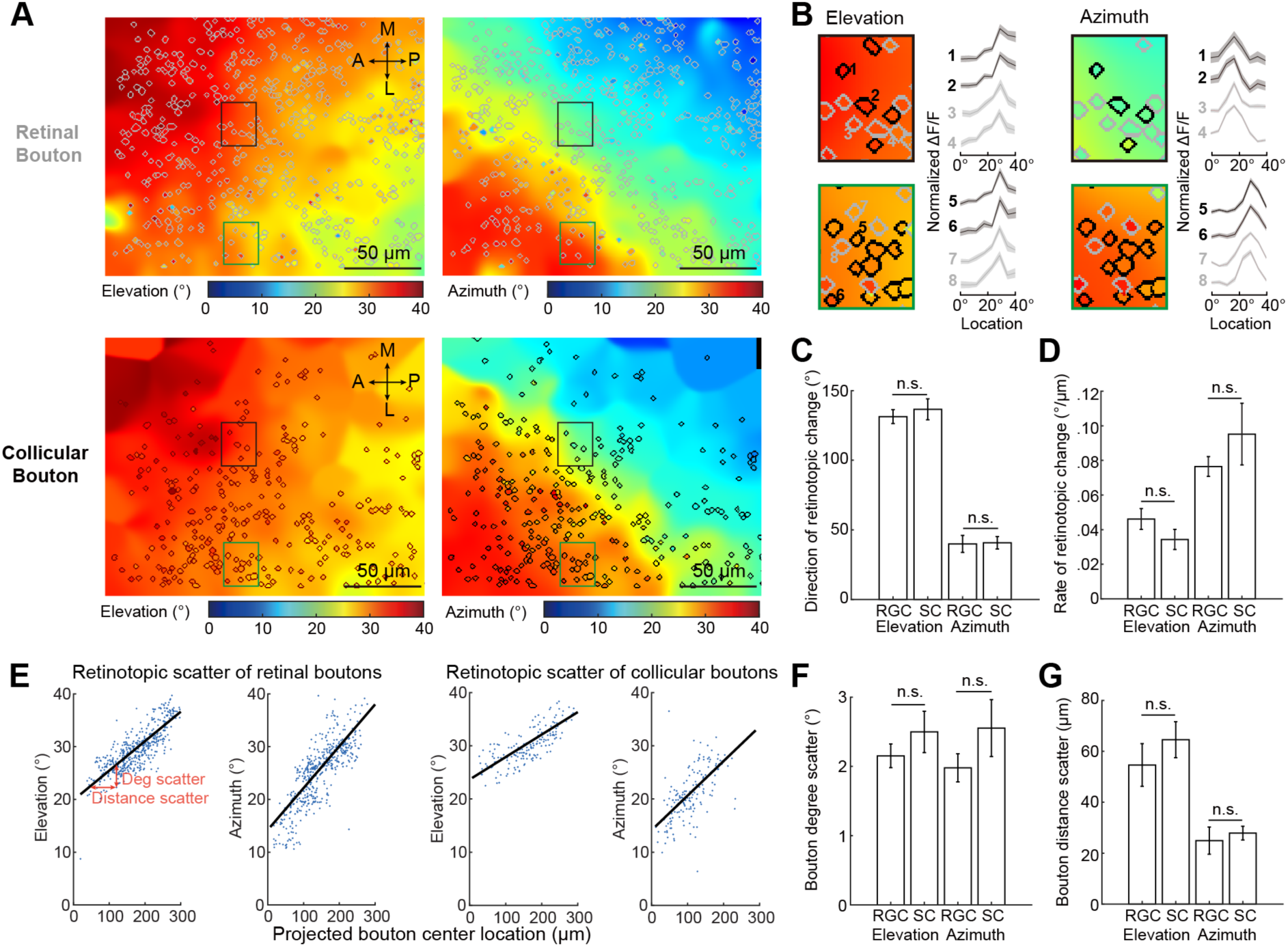
Similar Retinotopic Organization of Retinal and Collicular Axons. (A) Retinotopic maps of retinal (top) and collicular boutons (bottom) in the same FOV of the dLGN. Individual bouton masks are outlined by circles. Colors represent the preferred azimuth and elevation location. The background is a smoothed estimate of neuropil retinotopic preferences. (B) Magnified images illustrating retinotopic preferences in neighboring retinal (outlined in gray) and collicular boutons (outlined in black) from the elevation or azimuth retinotopic maps in A. Right: Normalized tuning curves of elevation or azimuth locations from example boutons (mean ± SEM). (C) No significant differences were found between retinal and collicular boutons in their directions of maximal retinotopic gradient for visual stimulation along either the elevation or azimuth axis (elevation axis: retinal boutons 131° ± 5°, collicular boutons 137° ± 8°, p = 0.53; azimuth axis: retinal boutons 40° ± 6°, collicular boutons 41° ± 4°, p = 0.34). Results in C-G are from 10 FOVs of 7 Rorβ-Cre mice. (D) The rates of change in retinotopic preferences were comparable between retinal and collicular boutons (elevation axis: retinal boutons 0.046 ± 0.006, collicular boutons 0.034 ± 0.006, p = 0.61; azimuth axis: retinal boutons 0.077 ± 0.006, collicular boutons 0.095 ± 0.018, p = 0.73). (E) Left: The scatterplot of retinal boutons’ retinotopic preferences versus their projected locations along the elevation or azimuth axis of their retinotopic progression map from the same FOV in A. Blue dots: individual boutons in the FOV. Black line: the linear fit of the data points. The slope of the black line indicates the average rate of the retinotopic progression (‘rate of change’). Deviation in retinotopic preference (’deg scatter’) and distance from the expectation (’distance scatter’) for a given bouton are shown with orange lines. Right: Scatterplot of retinotopic preferences for collicular boutons from the same FOV in A. (F) No significant differences were found between retinal and collicular boutons in their degree of scatter along either the elevation or azimuth axis (Elevation: retinal boutons 2.15 ± 0.17, collicular boutons 2.50 ± 0.30, p = 0.26; azimuth: retinal boutons 1.98 ± 0.21, collicular boutons 2.55 ± 0.41, p = 0.25). (G) Retinotopy was disordered below the spatial scale of ∼60 μm (elevation) or ∼30 μm (azimuth) for both collicular and retinal boutons (elevation: retinal boutons 54.63 ± 8.38, collicular boutons 64.57 ± 7.09, p = 0.88; azimuth: retinal boutons 24.92 ± 5.31, collicular boutons 27.86 ± 2.69, p = 0.45). All bar plots show mean ± SEM. Statistical tests in C, D, F, and G: linear mixed-effects model.

Our quantification revealed that retinal and collicular boutons progressed their retinotopy along similar axes and at comparable rates (Figure 2C and 2D). To further quantify this organization, we plotted the receptive field center preferences of individual boutons as a function of their projected positions along the elevation or azimuth axis of the retinotopic progression map in the FOV and linearly fitted the relationship (Figure 2E). Based on this linear fitting, we defined the distance scatter and degree scatter for individual boutons as their displacement from the expected projected positions and expected receptive field centers (Figure 2E). We observed average retinotopic displacements of ∼2° in both retinal and collicular boutons along both axes in visual space (Figure 2F) and a similar two-fold difference in the distance scatter between the elevation and azimuth axes for both retinal and collicular boutons (Figure 2G). These quantifications revealed a highly consistent retinotopic organization between retinogeniculate and colliculogeniculate axons, including their axes and rates of retinotopic progression and the level of organizational precision along both the elevation and azimuth axes.

### Similar Fine-scale Functional Clustering of Retinogeniculate and Colliculogeniculate Boutons

Next, we examined how colliculogeniculate boutons are organized below the scale of retinotopic precision. We previously discovered that retinal boutons from distinct axons often share one or several visual feature preferences when they are within a distance of ∼6 μm, the same spatial scale for bouton clusters on the proximal dendrites of dLGN neurons^59^. We observed similar fine-scale bouton clusters among collicular boutons in the dLGN (inset of Figure 3C). We asked whether fine-scale functional logic for arranging retinal boutons also applied to collicular boutons. To assess the functional similarity between boutons, it is essential to avoid analyzing pairs of boutons from the same axon. This is achieved by classifying boutons into distinct axons by their spontaneous correlation^59^ (see also Figure S2). As shown in Figure 3B and 3D, we observed examples of neighboring retinal or collicular boutons from different DS axons that shared similar direction preferences.

**Figure 3.**
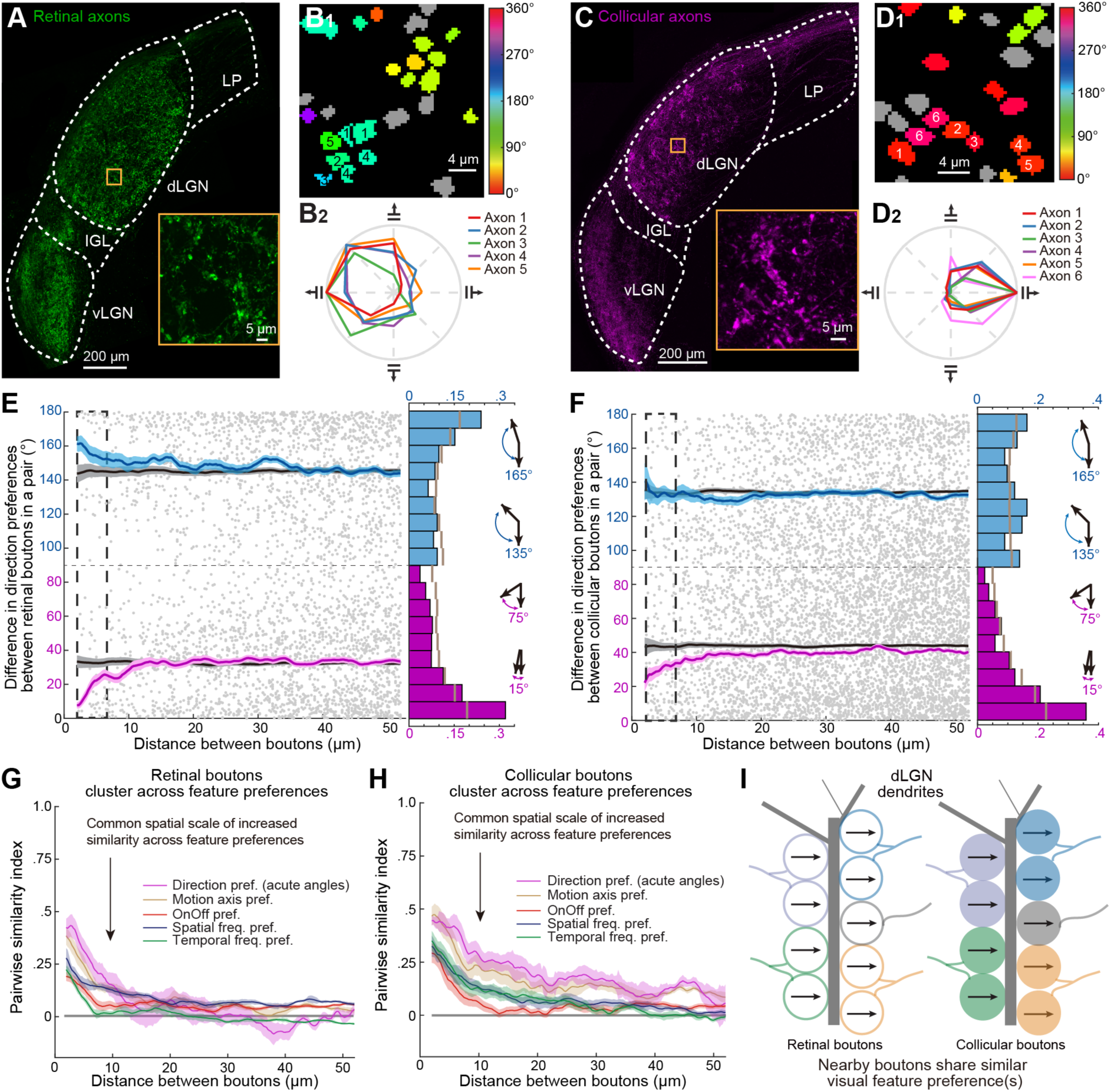
Fine-scale Functional Clustering of Retinal Boutons and Collicular Boutons, Respectively. (A) An example immunostaining image of retinal boutons in the dLGN (maximal z projection across 3 μm). Inset: Clusters of retinal boutons at higher magnification (maximal z projection over 3.5 μm). LP, lateral posterior nucleus; IGL, intergeniculate leaflet; vLGN, ventral LGN. (B) B_1_: Retinal boutons from a subregion of an example FOV, each colored according to its preferred motion direction (gray: boutons not exhibiting direction selectivity). B_2_: Average direction tuning curves for DS axons numbered in B_1_. Boutons from the same axon are labeled by the same axon number. These nearby axons preferred similar directions of motion (Axons 1–5). (C-D) Similar to A and B, but for collicular boutons. Collicular boutons were labeled by injecting Cre-dependent calcium indicators into the SC of Rorβ-Cre mice (4 μm maximal z projection image). C inset: 4.5 μm maximal z projection of a subregion at higher magnification. Note the absence of bouton clusters in the vLGN. (E) Left: The absolute difference in direction preferences as a function of inter-bouton spacing for pairs of retinal boutons that exhibited significant direction selectivity and belonged to different axons from an example FOV. Red and Blue lines, mean differences (4 μm-wide sliding window), generated from 1877 (with acute angles) or 1464 (with obtuse angles) pairs of boutons. Black line, similar analysis following random permutations of direction preference differences across all DS bouton pairs spaced 2–155 μm apart. Gray dots, individual retinal bouton pairs. Middle: Distributions of direction preference differences for pairs of boutons spaced 2-6 μm apart (magenta and blue bars, bouton pairs with differences between 0° and 90° or between 90° and 180°, respectively) or 2-155 μm apart (short brown lines), pooled from 18 FOVs of 8 mice. Right: Schematics illustrating the direction preference differences for distinct pairs of boutons. (F) Similar to E, but for collicular boutons (generated from 4316 (acute) or 2493 (obtuse) collicular bouton pairs). (G) Pairwise similarity indices for several visual feature preferences as a function of spatial distances between pairs of retinal boutons from different axons. Each color represents a visual feature. Gray, results from randomly shuffled data. (H) Similar pairwise analyses as in G but for collicular boutons. (I) Schematic showing that nearby retinal or collicular boutons from different axons often share similar visual feature preferences. All plots show mean ± SEM. Results in E-H are from 18 FOVs of 8 Rorβ-Cre mice.

Quantification of the pairwise similarity in feature preferences among retinal boutons from the dual-color imaging experiment confirmed our previous finding that nearby retinal boutons from distinct axons exhibit similar feature preferences^59^. For example, for all pairs of direction-selective retinal boutons from distinct axons in a FOV, we quantified the absolute differences in their preferred directions as a function of their spatial distances. We then calculated the average differences for all pairs within a sliding window of distances. We also assessed the sliding average of the chance level by shuffling the direction preferences across all direction-selective boutons within 155 µm in the FOV. When considering pairs of DS retinal boutons with preferences differing by less than 90° (i.e., acute angles), we observed functional clustering of similar direction preferences for pairs spaced 2–6 µm apart. For pairs of DS retinal boutons with preferences differing by more than 90° (i.e., obtuse angles), we observed significant fine-scale functional clustering in pairs with nearly opposite direction preferences (Figure 3E). To test whether similar fine-scale functional clustering of retinal boutons also existed for other visual features, we defined a pairwise similarity index for all the measured visual features (0: chance-level similarity; 1: identical feature preferences). We observed that the similarity indices were significantly higher than chance levels when retinal boutons were within a distance of ∼ 10 µm for each of the visual features examined, including motion axis, On/Off preference, spatial frequency, and temporal frequency.

Consistent with the results for retinal boutons, quantification of the pairwise similarity in feature preferences among collicular boutons revealed fine-scale functional clustering of collicular boutons according to similar visual feature preferences. When considering pairs of DS collicular boutons with preferences differing by less than 90°, we observed significantly smaller differences in direction preferences for pairs spaced 2–6 µm apart. For pairs of DS collicular boutons with preferences differing by more than 90°, we did not observe significant fine-scale functional clustering for nearby boutons tuned to opposite directions (Figure 3F). When considering other visual features, including motion axis, On/Off preference, spatial frequency, and temporal frequency, we observed that nearby collicular boutons from different axons often exhibited similar feature preferences, as evidenced by the above-chance-level pairwise similarity (Figure 3H). This fine-scale functional clustering of collicular boutons occurred at a spatial scale of approximately 10 µm, similar to that of retinal boutons (Figure 3I).

### Fine-Scale Functional Clustering between Nearby Retinal and Collicular Boutons

The similarity in the fine-scale functional organization of retinal and collicular boutons raises the question of whether there is a functional convergence between them. EM and anatomical studies suggest that collicular boutons could be adjacent to retinal boutons on the same dendrites of dLGN neurons^18^. Immunostaining showed that neighboring collicular and retinal boutons formed cluster-like structures (Figure 4A). Moreover, we observed shared direction preferences among neighboring retinal and collicular axons (Figure 4B). We calculated the feature preference difference between retinal and collicular boutons as a function of their spatial distance in each FOV. For each visual feature, we observed significantly positive pairwise similarity indices for retinal and collicular boutons within a distance of ∼10 µm. This indicates that neighboring retinal and collicular boutons often possess similar feature preferences and exhibit functional clustering (Figure 4C). These findings suggest that the fine-scale functional logic we previously discovered — where axonal boutons within a distance of ∼10 µm often share their visual feature preferences — not only applies to the organization of each input source separately but also to the convergence of inputs from different visual centers, thus providing a general principle for arranging excitatory inputs in the dLGN (Figure 4D). The functional convergence between retinal and collicular boutons suggests a synergy between the visual functions of both sources of inputs and implies that the colliculogeniculate input may reinforce the visual information carried by the retinal axons in their shared target dLGN neurons.

**Figure 4.**
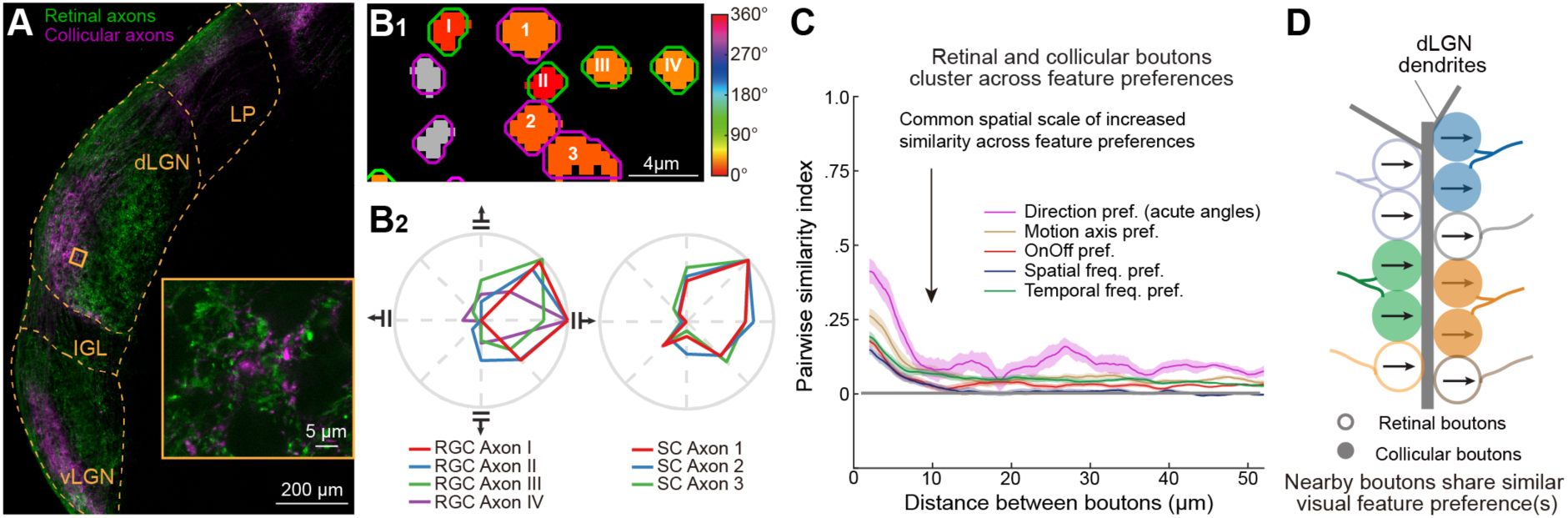
Fine-scale Functional Clustering Between Retinal and Collicular Boutons. (A) An example immunostaining image of retinal and collicular boutons in the dLGN. Retinal boutons were labeled by injecting AAV-GCaMP6f into the retina, and collicular boutons were labeled by injecting AAV-FLEX-jRGECO1a into the SC of Rorβ-Cre mice (6 μm maximal z projection). Inset: Higher magnification of retinal and collicular axonal boutons (a 0.8 μm maximal z projection). (B) B_1_: Retinal boutons (outlined in green) and collicular boutons (outlined in magenta) from a subregion of an example FOV, each colored according to its preferred motion direction (gray: boutons not exhibiting direction selectivity). B_2_: Average direction tuning curves for DS axons numbered in B_1_. These nearby axons preferred similar directions of motion (Axons I-IV are retinal axons, and Axons 1-3 are collicular axons). (C) Pairwise similarity indices for several visual feature preferences as a function of spatial distances for retinal-collicular bouton pairs (mean ± SEM). Different colors represent different visual features. Gray, results from the same analysis for randomly shuffled data. N = 18 FOVs from 8 mice. (D) Schematic shows that nearby retinal and collicular boutons could exhibit similar feature preferences.

The pairwise analysis above considered each visual feature separately. Individual retinal and collicular boutons often multiplex distinct channels of visual information by being tuned to multiple visual features. Using a group-wise analysis to consider the convergence of multiple visual features among nearby groups of boutons, we previously demonstrated two distinct modes of fine-scale functional convergence among retinal boutons^59^. Under “relay mode” convergence, nearby retinal boutons share multiple feature preferences in common, whereas under “combination mode” convergence, some preferences are matched while others are not.

Here, we investigated the occurrence of the two modes of convergence between retinal and collicular boutons (Figure S3A). We observed examples of nearby collicular and retinal boutons possessing similar preferences for multiple visual features, such as the motion axis, spatial frequency, and temporal frequency (Figure S3C). To determine the similarity of feature preferences among nearby retinal and collicular boutons, we selected all pixels that had a group of retinal and collicular boutons within a 6 μm radius (typically 3-7 boutons). For each valid pixel, we calculated a group-wise similarity index for the preferences of each feature among these boutons (1: identical feature preference; 0: chance-level similarity) using a method adapted from our previous study^59^. The group-wise similarity maps for retinal and collicular boutons in Figure S3C are shown in Figure S3D. Consistent with pair-wise analysis, group-wise analysis demonstrated significant fine-scale functional clustering between retinal and collicular boutons for each visual feature (Figure S3G). An overlay of similarity indices under each feature in this example FOV illustrated the existence of both ‘relay’ (significant similarity indices for multiple visual features) and ‘combination’ modes (significant similarity indices for only one visual feature) (Figure S3E), with the values of the similarity indices illustrated in a scatterplot (Figure S3F). Summarizing all FOVs, we observed combination-mode bouton groups with similar preferred motion axes and relay-mode convergence between retinal and collicular boutons when they shared preferred motion axes (Figure S3H).

### Silencing Colliculogeniculate Input Suppressed Visual Responses in dLGN Neurons

Previous pharmacological silencing of the SC in cats and rabbits led to both enhancement and suppression of visual responses in different dLGN neurons^65–67^. However, these studies did not establish a connection between the visual information transmitted by colliculogeniculate axons and their functional impact on the dLGN, nor did they examine whether colliculogeniculate input differentially influences distinct dLGN neuron subtypes and contributes to their tuning properties, potentially shaping the flow of visual information in the retino-geniculo-cortical pathway^60^. To investigate this, we selectively and acutely suppressed the excitatory colliculogeniculate input by injecting AAVs carrying Cre-dependent chemogenetic inhibitor, hM4Di-mCherry, into the SC of Rorβ-Cre mice and expressed GCaMP6f in dLGN neurons to assess visual responses (Figure 5A)^68^. Labeling of the colliculogeniculate projection was verified through *in vivo* imaging and *post hoc* histology. We also examined the retinae to confirm that RGCs were not retrogradely labeled from the AAV injection into the SC (data not shown). We recorded calcium responses to the same visual stimuli in the same dLGN neurons, first following saline injection and then after administration of Clozapine N-oxide (CNO), the ligand that activates hM4Di (Figure 5B and 5C). Control experiments were performed using AAVs carrying Cre-dependent mCherry injected into the SC.

**Figure 5.**
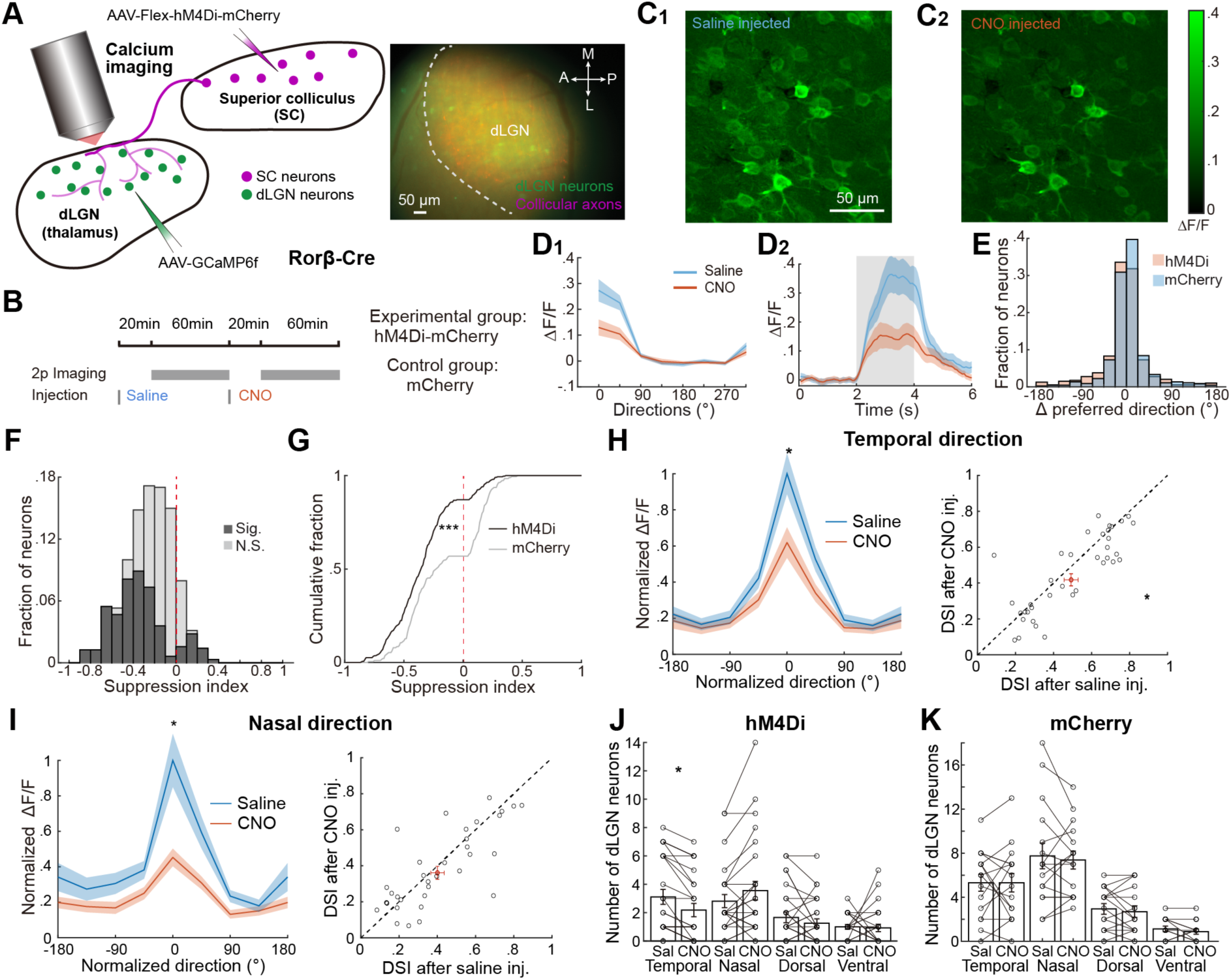
Suppression of Colliculogeniculate Neurons Reduced Visual Responses in dLGN Neurons and Decreased the Number of dLGN Neurons Preferring Temporal Direction. (A) Left: schematic of silencing colliculogeniculate axons via the inhibitory DREADD hM4Di-mCherry and measuring visual responses in dLGN neurons via GCaMP6f. Right: epifluorescence image through the cranial window showing GCaMP6f-expressing thalamic neurons (green) and hM4Di-mCherry-expressing collicular axons (magenta). The dLGN was identified by retinotopy, with its anterior border outlined. (B) Experimental timeline: 60-min imaging of visual responses in the same dLGN neurons was performed 20 minutes after saline injection and again 20 minutes after CNO injection. (C) Maximal projections of ΔF/F image from an example FOV after saline (C_1_) and after CNO (C_2_) injections, respectively. (D) D_1_: Direction tuning curves of an example dLGN neuron after saline injection (blue) and CNO injection (orange). D_2_: Average activity trace in response to 0° drifting gratings; gray indicates the visual stimulus presentation period. Mean ± SEM. (E) Distributions of changes in the preferred direction (Preferred direction_CNO_ – Preferred direction_Saline_) were similar between hM4Di-injected mice and mCherry-injected mice (p = 0.52). E, G-K: linear mixed-effects model; hM4Di: N = 27 FOVs from 11 mice; mCherry: N = 16 FOVs from 6 mice. (F) The majority of dLGN neurons showed negative suppression indices (-1, decrease in responses; 1, increase in responses) after silencing colliculogeniculate input. Gray: not significantly affected; black: significantly affected (303 out of 641 neurons showed significant changes in their visual responses). (G) The cumulative distribution of suppression indices from significantly impacted dLGN neurons in hM4Di-vs. mCherry-injected mice (p < 0.0001). (H) Left: Normalized tuning curves of all temporal-direction-preferring dLGN neurons after CNO injection (orange) compared to after saline injection (blue) in hM4Di-labeled mice (n = 41 neurons from 27 FOVs across 11 mice; p = 0.65 at -180°, p = 0.010 at 0°). The preferred direction of each neuron was set to 0°. Right: DSI values across these dLGN neurons were significantly lower after CNO injection compared to after saline injection (DSI after saline inj., 0.493 ± 0.033, DSI after CNO inj., 0.418 ± 0.036; p = 0.02). The red cross represents mean ± SEM. Results in H-I are from hM4Di-injected mice. (I) Similar to H but for nasal-direction-preferring neurons (n = 39 neurons; p = 0.29 at -180°, p = 0.011 at 0°). Right: DSIs after saline and after CNO injections were not significantly different (DSI after saline inj., 0.401 ± 0.036, DSI after CNO inj., 0.365 ± 0.036; p = 0.36). (J) Only the number of temporal-direction-preferring DS dLGN neurons was significantly reduced after administering CNO in hM4Di-labeled mice (temporal saline: 3.11 ± 0.52; temporal CNO: 2.18 ± 0.46; nasal saline: 2.82 ± 0.46; nasal CNO: 3.55 ± 0.66; dorsal saline: 1.67 ± 0.38; dorsal CNO: 1.26 ± 0.31; ventral saline: 1.00 ± 0.17; ventral CNO: 0.93 ± 0.26; temporal p = 0.046; nasal p = 0.24; dorsal, p = 0.39; ventral, p = 0.82). Temporal: 0°-60°; dorsal: 90°-150°; nasal: 180°-240°; ventral: 270°-330°. (K) Same as H but in mCherry-labeled mice (temporal saline: 5.31 ± 0.76; temporal CNO: 5.31 ± 0.83; nasal saline: 7.75 ± 1.17; nasal CNO: 7.38 ± 0.82; dorsal saline: 2.94 ± 0.49; dorsal CNO: 2.69 ± 0.53; ventral saline: 1.13 ± 0.26; ventral CNO: 0.88 ± 0.24; temporal p = 0.98; nasal p = 0.71; dorsal, p = 0.65; ventral, p = 0.41). All tuning curves and bar plots are shown as mean ± SEM. *, p < 0.05; **, p < 0.01; ***, p < 0.001. G-K: linear mixed-effects model. See also Figures S4, S5, and S6.

We observed significant reductions in the maximum responses of most dLGN neurons after CNO injection compared to saline injection (Figure 5C). Although the preferred direction remained unchanged in example DS dLGN neurons, responses for the preferred direction markedly decreased following CNO injection (Figure 5D). At the population level, motion direction and axis preferences were not significantly altered after silencing collicular input (Figure 5E and S4F). To quantify changes in neuronal activity for the preferred direction, we defined a suppression index (SuI), with a negative SuI indicating activity decrease after silencing the SC. Silencing collicular input suppressed visual responses in the majority of dLGN neurons, with the SuI distribution significantly left-shifted compared to the mCherry control group (Figure 5F and 5G). Additionally, 85% of dLGN neurons in hM4Di- injected mice showed reduced baseline activity, compared to 64% in mCherry-injected mice (Figure S4A). However, the SuIs of baseline activity were an order of magnitude smaller than the SuIs of visual responses (Figure 5G and S4A), indicating that silencing colliculogeniculate input imposed a stronger suppressive effect on visual responses of dLGN neurons.

Similar to retinal and collicular inputs, we classified dLGN neurons into DS (direction-selective), AS (axis-selective), BrT (broadly-tuned), and Sup (suppressed) functional categories based on their responses to full-screen drifting gratings. Following CNO injection, DS, AS, and BrT dLGN neurons in hM4Di-injected mice showed stronger suppressive effects compared to those in mCherry-injected mice (Figure S4B). In contrast, the impacts on Sup dLGN neurons were not significantly different between the two groups, consistent with the small fraction of Sup boutons among the colliculogeniculate input (Figure 1F).

To limit the confounding effects of arousal modulation and potential drift of arousal states, we assessed the silencing effects using trials with closely matched pupil sizes (see STAR Methods). For all analyses, we excluded trials with pupil areas in the lower or upper 20% of the pupil area distribution for each imaging session. Our results showed that dLGN neurons in hM4Di-injected mice were significantly more suppressed than those in mCherry-injected mice, even when we varied the pupil area thresholds (Figure S4C-S4E). Moreover, we calculated the suppression index from subsets of trials with matched pupil positions and limited pupil motion to minimize the influence of pupil position and eye movement. We found no significant change in SuIs after matching trials (Figure S4F-S4H).

Moreover, we measured the effects of silencing the Rorβ-Cre-positive colliculogeniculate input using a two-day imaging paradigm (Figure S5). Here, we separately administered saline and CNO in the middle of imaging sessions conducted on two consecutive days. We compared the visual responses to the same stimuli before and after injecting saline on day 1 and CNO on day 2 and calculated the SuIs for each day (Figure S5A). We then compared the SuIs from the two sessions to assess the impact of saline vs. CNO. Consistent with the same-day imaging results, we found that SC inactivation suppressed responses in the majority of dLGN neurons, and the suppression was significantly stronger than that in the control group (Figure S5D).

Since it is unclear whether Rorβ-Cre labels all the excitatory colliculogeniculate neurons, we also silenced excitatory SC neurons in Vglut2-Cre mice. Additionally, Vglut2-Cre allowed us to record visual responses specifically from excitatory dLGN thalamocortical cells (TCs) (Figure S6A). Consistent with the result from silencing Rorβ-Cre-positive SC neurons, the majority of TCs exhibited suppressed responses following CNO administration in hM4Di-injected mice, with the suppression significantly stronger than in mCherry-injected control mice (Figure S6B-S6C). Moreover, the suppression effect on baseline activity of TCs was much smaller than on their visual responses after silencing Vglut2-positive collicular neurons (Figure S6D). These findings confirm that silencing excitatory colliculogeniculate input significantly reduces visually-evoked activity in dLGN TCs. In other words, collicular input significantly and positively contributes to response magnitudes in the majority of dLGN TCs.

### Silencing Colliculogeniculate Input Reduced Motion Selectivity in Specific Subsets of dLGN Neurons

Given that collicular input carries significant motion-selective signals to the dLGN, we tested whether silencing colliculogeniculate neurons would affect motion selectivity in direction-selective (DS) and axis-selective (AS) dLGN neurons. We further divided DS and AS dLGN neurons according to their preferred motion directions or axes. As expected, we found that the peak responses in these subgroups were significantly reduced after CNO injection in hM4Di-injected mice, but not in mCherry-injected mice (Figure 5H, 5I, and S4J-S4P). However, our findings also revealed that collicular input differentially impacted motion selectivity among distinct dLGN neuron subgroups. Specifically, direction selectivity significantly decreased in DS dLGN neurons preferring the temporal direction but not in those preferring the nasal direction (Figure 5H and 5I). Additionally, axis selectivity significantly decreased in AS dLGN neurons tuned to the horizontal axis but not in those tuned to the vertical axis (Figure S4J and S4K). Following CNO administration in hM4Di-injected mice, the number of DS neurons only decreased significantly for the temporal-direction-preferring subgroup but not for those tuned to other motion directions (Figure 4J). The number of horizontal-axis-preferring AS neurons also decreased significantly, whereas the number of vertical-axis-preferring neurons did not (Figure S4L). Motion selectivity and the number of motion-selective neurons remained unchanged in mCherry-injected control mice (Figure 5K, S4M-S4Q). These results were replicated using a two-day paradigm with Rorβ-Cre mice (Figure S5E-S5H) and the same paradigm with Vglut2-Cre mice (Figure S6E-S6P). These findings align with the observation that a larger fraction of collicular boutons prefer the temporal direction or horizontal motion compared to retinal boutons (Figure 1F) and demonstrate that the colliculogeniculate projection significantly and selectively contributes to motion selectivity in dLGN neurons.

### Silencing Colliculogeniculate Input Resulted in Distinct Types of Suppression

Silencing colliculogeniculate input could potentially suppress responses in direction-selective (DS) and axis-selective (AS) dLGN neurons in several ways, including altering the preferred motion direction or axis (displacement suppression)^69^, reducing the response magnitude equally across directions (subtractive suppression)^70^, scaling down response magnitude linearly across directions (divisive suppression)^70^, or imposing additional suppression at the preferred motion direction or axis (selective suppression) (Figure 6A). We ruled out displacement suppression since the preferred direction/axis in DS and AS dLGN neurons remained largely unchanged after silencing collicular input (Figure 5E and S4F). To identify dLGN neurons exhibiting the other three forms of suppression, we compared the suppression ratios in the preferred and null directions (opposite direction for DS and both orthogonal directions for AS neurons). Subtractive suppression shows a larger suppression ratio in the null direction, divisive suppression has similar ratios across all directions, and selective suppression exhibits a higher ratio in the preferred direction (Figure 6B).

**Figure 6.**
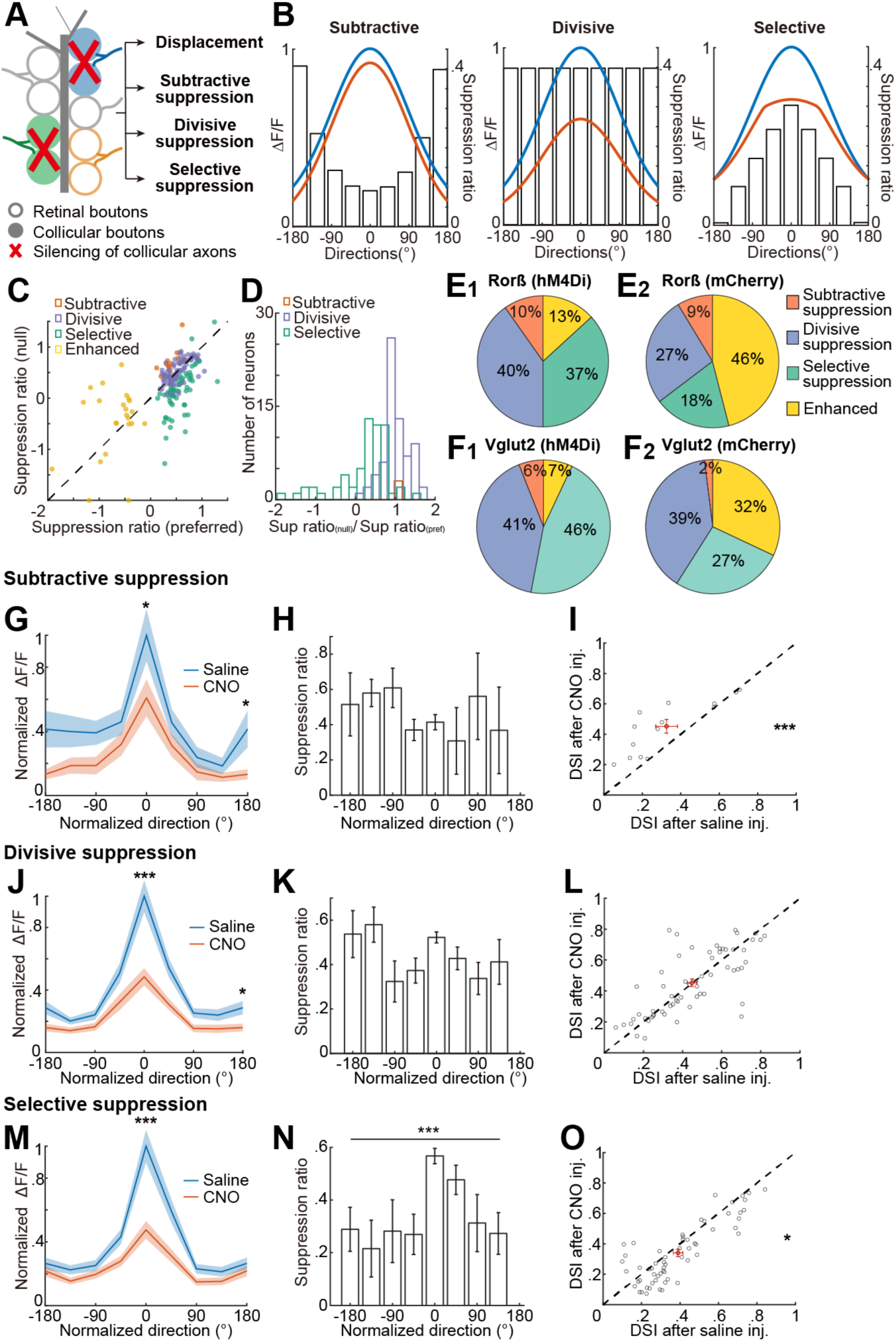
Silencing Colliculogeniculate Input Divisively or Selectively Suppressed Direction-selective dLGN Neurons. (A) Schematic of potential outcomes on direction-selective (DS) dLGN neurons after silencing collicular input. Displacement, shift of the preferred motion direction; subtractive, divisive, and selective suppression are defined in (B). (B) Schematic illustrating direction tuning curves of DS dLGN neurons under three types of suppression after silencing collicular input. Subtractive: Responses are shifted down equally across directions, with the suppression ratio peaking in the null direction. Divisive: Responses are suppressed by the same ratio across all directions. Selective: Responses are suppressed by a higher ratio in the preferred direction. (C) Scatterplot of the suppression ratio in the null direction vs. the preferred direction for DS dLGN neurons under subtractive (orange), divisive (blue), or selective (green) suppression, or enhancement following collicular input silencing. Results in C, D, G-O are from 27 FOVs of 11 hM4Di-injected Rorβ-Cre mice. (D) Distinct distributions of the ratio between the suppression ratios in the null direction and the preferred direction for dLGN neurons under distinct types of suppression in C. (E) Larger fractions of DS dLGN neurons underwent divisive and selective suppression in (E_1_) hM4Di-injected Rorβ-Cre mice (n = 17, 70, 64, and 23 neurons for subtractive, divisive, selective suppression, and enhanced categories, respectively) compared to (E_2_) mCherry-injected Rorβ-Cre mice (n = 13, 40, 28, and 69 for subtractive, divisive, selective suppression, and enhanced categories). p<0.001, Chi-squared test; p = 0.73, 0.010, 0.0003, and 8.9*10^-10^ for subtractive, divisive, selective, enhanced categories, respectively, *post hoc* Pearson residuals test. (F) (F_1_) hM4Di-injected Vglut2 mice had a significantly higher proportion of selectively-suppressed excitatory DS neurons (n = 10, 70, 78 and 12 for subtractive, divisive, selective, and enhanced categories, respectively) compared to (F_2_) mCherry-injected Vglut2 mice (n = 4, 62, 44 and 51 for subtractive, divisive, selective, and enhanced categories, respectively). p<0.001, Chi-squared test; p = 0.12, 0.62, 0.0004, and 1.4*10^-7^ for subtractive, divisive, selective, and enhanced categories, respectively, *post hoc* Pearson residuals test. (G) Averaged tuning curves after saline injection and after CNO injection for the DS dLGN neurons that exhibited subtractive suppression following CNO injection (p = 0.04 at -180°, p = 0.02 at 0°; n = 17 neurons). Peak directions are set to 0°. (H) Suppression ratios across all normalized directions for neurons in G (p = 0.14, Kruskal-Wallis test). (I) Significantly different DSI after CNO injection compared to saline injection in subtractively-suppressed DS neurons (DSI: after saline inj. 0.326 ± 0.044, after CNO inj. 0.452 ± 0.056; p < 10^-3^). (J) Averaged tuning curves after saline injection and after CNO injection for the DS dLGN neurons that exhibited divisive suppression following CNO injection (p = 0.012 at -180°, p < 10^-4^ at 0°; n = 70 neurons). (K) Suppression ratios across all normalized directions were not significantly different for neurons in J (p = 0.056; Kruskal-Wallis test). (L) DSI values after CNO injection and after saline injection were not significantly different in divisively-suppressed DS dLGN neurons (DSI: after saline inj. 0.448 ± 0.024, after CNO inj. 0.453 ± 0.024; p = 0.90). (M) Similar to (G) but for the DS dLGN neurons exhibiting selective suppression for the preferred direction after CNO injection (p = 0.54 at -180°, p < 0.001 at 0°; n = 64 neurons). (N) Suppression ratios for neurons in M were significantly different across normalized directions (p < 0.001, Kruskal-Wallis test; p = 0.0006 between -180° and 0°, p = 0.02 between -135° and 0°, p = 0.02 between -90° and 0°, p = 0.02 between -45° and 0°, p = 0.011 between 0° and 135°, Dunn’s multiple comparisons test). (O) In selectively-suppressed DS dLGN neurons, the DSI after CNO injection was significantly lower than after saline injection (DSI: after saline inj. 0.405 ± 0.024, after CNO inj. 0.352 ± 0.024. p = 0.04). All the data was plotted as mean ± SEM. *, p < 0.05; **, p < 0.01; ***, p < 0.001. G, I, J, L, M and O: linear mixed-effects model. See also Figures S7, S8.

DS dLGN neurons undergoing different types of suppression were separated in the scatter plot and histogram of the relative suppression ratios (Figure 6C and 6D). Following colliculogeniculate silencing, most DS dLGN neurons showed either divisive or selective suppression, while a smaller proportion exhibited subtractive suppression (Figure 6C-6F). Divisively-suppressed and selectively-suppressed neurons contributed comparably large fractions of DS dLGN neurons in hM4Di-injected mice (Figure 6E_1_ and 6F_1_). In contrast, in mCherry-injected mice, a smaller fraction of neurons exhibited suppression after CNO injection, and the selectively-suppressed group was about two-thirds the size of the divisively-suppressed group, significantly differing from the proportions in hM4Di-injected mice (Rorβ-Cre: p<0.0001; Vglut2-Cre: p<0.0001, Chi-squared test) (Figure 6E_2_ and 6F_2_).

We further characterized the groups exhibiting subtractive, divisive, or selective suppression. For neurons in each group, we averaged their normalized tuning curves following saline and CNO injections, aligned by their peak directions. All groups showed a significant decrease in response magnitude in their peak directions (Figure 6G, 6J, and 6M). We confirmed that the suppression ratio stayed similar across directions for neurons exhibiting divisive suppression but was significantly higher in the peak direction for neurons showing selective suppression (Figure 6K and 6N). Direction selectivity after CNO injection significantly decreased in selectively-suppressed neurons, increased in subtractively-suppressed neurons, and did not change in divisively-suppressed dLGN neurons (Figure 6I, 6L, and 6O). The visual response magnitudes before CNO injection were not significantly different between divisively and selectively-suppressed neurons, indicating that the differences in suppression in the null direction and preferred motion direction between the two populations were not caused by the ‘floor effect’ or saturation of the calcium indicator, respectively (Figure S7A)^71^.

Similarly, AS dLGN neurons also underwent subtractive, divisive, or selective suppression or enhanced their responses following CNO injection (Figure S7C and S7D). hM4Di-injected mice had a higher fraction of suppressed AS neurons than mCherry-injected mice, and the proportion of selectively- suppressed neurons relative to divisively-suppressed neurons was higher in hM4Di mice than in mCherry controls (Figure S7E). Moreover, the ASIs of the selectively-suppressed, but not divisively- suppressed, AS neurons were significantly reduced after collicular input silencing (Figure S7F-S7N). Consistent with our findings that temporal-direction-preferring DS neurons and horizontal-axis- preferring AS neurons both exhibited reduced motion selectivity after collicular input silencing (Figure 5H and S4J), we observed higher proportions of selectively-suppressed neurons among those tuned to this direction/axis compared to neurons tuned to the nasal or vertical direction/axis (Figure S7B and S7P).

Similar analyses on the effects of silencing excitatory SC neurons on thalamocortical neurons in Vglut2- cre mice yielded consistent results (Figure S8). Collectively, our findings demonstrate that motion selectivity in distinct DS or AS dLGN neurons can be differentially affected by collicular input silencing. Specifically, some subsets of dLGN neurons experienced divisive suppression across all directions, while others showed additional suppression in the preferred motion direction or axis, leading to reduced direction/axis selectivity.

### Sharply Tuned Collicular Input Enhanced dLGN Motion Selectivity

Collicular boutons exhibit higher motion selectivity compared to retinal boutons (Figure 1G-H) and provide ‘driver-like’ input to dLGN neurons^18^, offering a possible explanation for the observed selective suppression in dLGN neurons when collicular input is silenced (Figure 7A). We tested this hypothesis using *in silico* simulation of the visual responses of dLGN neurons. The tuning curve of a dLGN neuron was generated from a linear combination of randomly weighted retinal and collicular tuning curves, along with random noise. The retinal and collicular tuning curves were randomly sampled from the recorded DS retinal and DS collicular boutons, with peak directions aligned to a common direction (Figure 7B). Notably, the recorded collicular DSI is higher than the retinal DSI (Figure 1G). When collicular input was ‘silenced,’ the tuning curves of the dLGN neurons were only composed of the retinal input and random noise following the same distribution. The proportions of simulated dLGN neurons under different suppression types closely matched our *in vivo* findings (Figure 6E, 6F, and 7C; p = 0.10, Chi-squared test between Figure 6E and 7C). In contrast, when the DSI distribution of collicular input was made identical to that of retinal input, the proportion of selectively-suppressed dLGN neurons significantly decreased (Figure 7D; p = 0.006, Chi-squared test between Figure 6E and 7D). Further analyses of the peak-aligned normalized tuning curves, suppression ratios across directions, and DSIs after ‘silencing’ collicular input in the simulated dLGN neurons under the three types of suppression recaptured the *in vivo* results (Figure 7E-7M). These simulation results underscore the significant contribution of highly tuned collicular input to direction selectivity in dLGN neurons.

**Figure 7.**
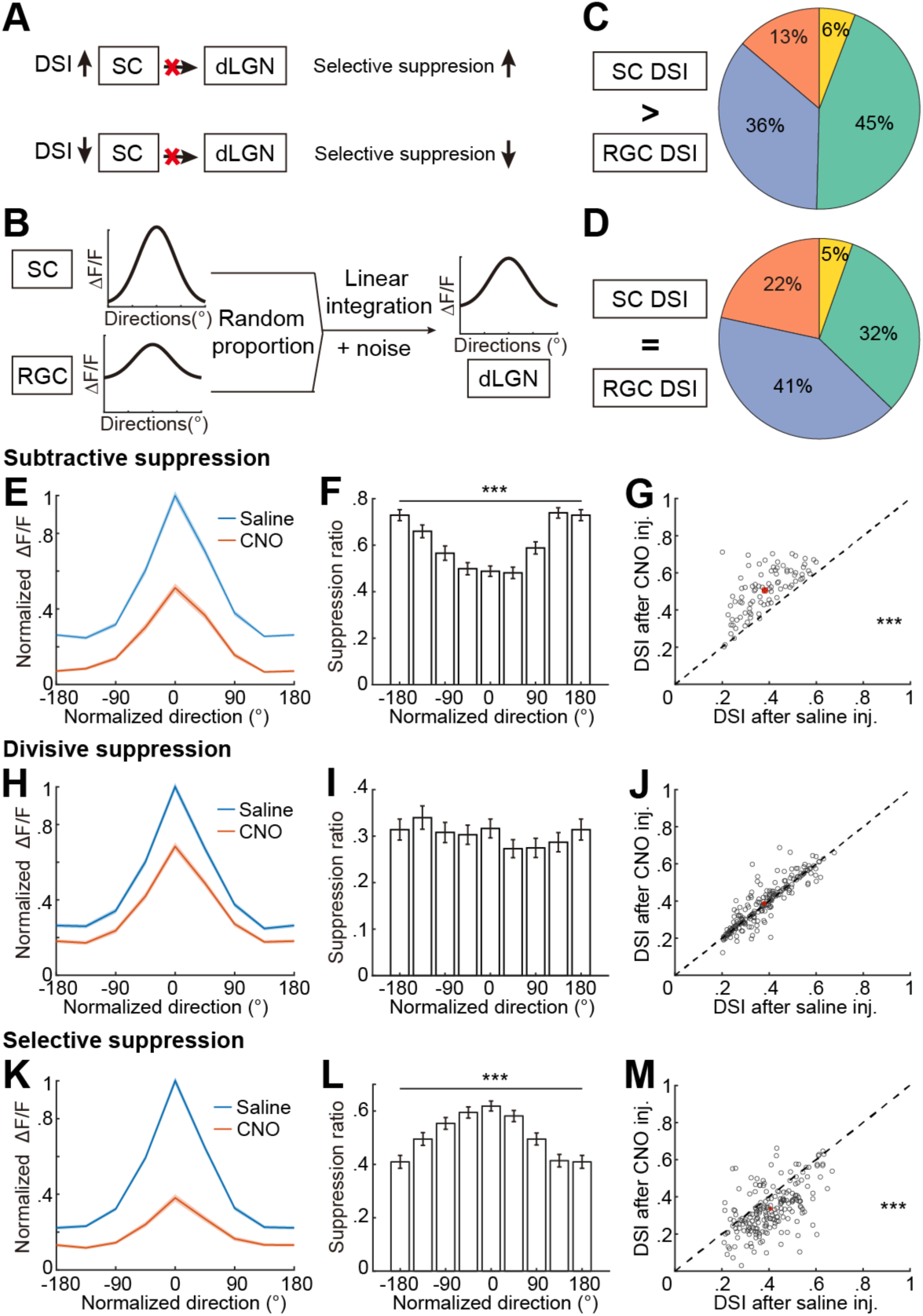
The High Proportion of Selective Suppression is Likely due to More Tuned Collicular Input. (A) Schematic illustrating the hypothesized relationship between the DSI of collicular input and the fraction of dLGN neurons exhibiting selective suppression following SC silencing. (B) Schematic of the simulation: Visual responses of dLGN neurons are established by linearly integrating random proportions of retinal and collicular inputs, along with random noise (see STAR Methods). Tuning curves of retinal and collicular inputs are randomly sampled from the recorded tuning curves of their respective populations. (C-D) When collicular boutons had higher DSIs than retinal boutons at the population level (using the recorded dataset of collicular and retinal boutons), the proportion of DS dLGN neurons showing selective suppression after SC input removal was larger compared to when collicular boutons had the same DSI distribution as retinal boutons. The distribution of suppression types in C is similar to Figure 6E_1_ (p = 0.10; Chi-squared test), whereas the distribution in D is significantly different from Figure 6E_1_ (p = 0.006; Chi-squared test; p = 0.0005, 0.16, 0.0008, 0.15 for the subtractive, divisive, selective suppression, and enhanced categories, respectively, *post hoc* Pearson residuals test). (E) Average tuning curves of simulated dLGN neurons exhibiting subtractive suppression after ‘CNO injection’ (following removal of collicular input) compared to after ‘saline injection’ (when dLGN neurons integrate both retinal and collicular inputs). (F) Suppression ratios of visual responses after ‘CNO injection’ were significantly different across normalized directions for neurons in E (n = 65 simulated neurons; p < 0.0001, Kruskal-Wallis test; p = 0.02 between -180° and -135°; p < 0.0001 between -180° and -90°, -45°, 0°, 45°; p = 0.021 between - 180° and 90°; Dunn’s multiple comparisons test). (G) DSI significantly increased after ‘CNO injection’ compared to after ‘saline injection’ in simulated dLGN neurons exhibiting subtractive suppression (DSI: after ‘saline inj.’ 0.375 ± 0.016, after ‘CNO inj.’ 0.499 ± 0.013; p < 0.0001). G, J, and M: Red: mean ± SEM; Wilcoxon signed-rank test. (H) Similar to (E) but for all simulated dLGN neurons showing divisive suppression after ‘CNO injection’. (I) Suppression ratios of visual responses after ‘CNO injection’ were not significantly different across normalized directions for neurons in H (n = 180 simulated neurons; p = 0.58, Kruskal-Wallis test). (J) DSI did not significantly change after ‘CNO injection’ compared to after ‘saline injection’ in simulated dLGN neurons exhibiting divisive suppression (DSI: after ‘saline inj.’ 0.389 ± 0.009, after ‘CNO inj.’ 0.396 ± 0.009; p = 0.47). (K) Similar to (E) but for all simulated dLGN neurons showing selective suppression after ‘CNO injection’. (L) Suppression ratios for neurons in K were significantly different across normalized directions (n = 225 simulated neurons; p < 0.0001, Kruskal-Wallis test; p < 0.0001 between 0° and -180°, -135°, 90°, 135°, Dunn’s multiple comparisons test). (M) DSI significantly decreased after ‘CNO injection’ compared to after ‘saline injection’ in simulated dLGN neurons exhibiting selective suppression (DSI: after ‘saline inj.’ 0.408 ± 0.008, after ‘CNO inj.’ 0.335 ± 0.008; p < 0.001). Data was plotted as mean ± SEM. *, p < 0.05; **, p < 0.01; ***, p < 0.001.

## DISCUSSION

Our results revealed that colliculogeniculate axons exhibited diverse visual response properties and shared similar retinotopic organization with retinogeniculate axons. At a spatial scale of ∼10 µm, nearby collicular boutons from different collicular axons often shared common feature preferences, following fine-scale functional arrangements similar to retinal boutons. Notably, neighboring collicular and retinal boutons also displayed similar visual feature preferences, suggesting they may act in concert to promote their shared information in common target neurons. Chemogenetic inhibition of collicular input decreased visual responses in the majority of dLGN neurons and reduced motion selectivity in dLGN neurons tuned to motion along the temporal direction or horizontal axis of motion. Simulation of dLGN visual responses *in silico* supported the hypothesis that sharp tuning of collicular axons could be responsible for the reduction of motion selectivity in dLGN neurons after silencing the SC. The findings suggest that excitatory collicular input plays a critical role in selectively enhancing motion selectivity in dLGN neurons. Our results elucidate how excitatory inputs from distinct brain regions collaborate to shape receptive field properties and reinforce specific information channels in the dLGN, thereby sculpting the information transmitted to downstream visual centers along the thalamocortical pathway^60^. Additionally, the data indicate that the dLGN and SC have a more complex role in the image-forming pathway than previously understood.

### Diverse and Sharp Visual Responses in Colliculogeniculate Axons

The superficial SC comprises several neuron types with distinct visual response properties and functions, including stellate, narrow-field, wide-field, and horizontal cells^11,29,48,54,72–75^. Previous mouse studies demonstrated that inhibitory SC projection neurons primarily target the parabigeminal nucleus (PBG) and the ventral LGN (vLGN)^31,76^, whereas excitatory colliculogeniculate neurons predominately originate from stellate cells in the upper SGS^11,54^. In vivo electrophysiological recordings demonstrated that stellate cells, including those at the dorsal sSC surface, respond to full-field stimulation and possess small receptive fields; however, they are rarely direction-selective despite showing a broad distribution of response properties^11^. In vivo two-photon imaging of a large population of superficial SGS neurons revealed that the topmost layer of the SGS is enriched with neurons that have compact receptive fields, overlapping On and Off subregions, and high direction selectivity^49,72,77^. However, the extent of overlap between stellate cells and highly direction-selective cells in the upper SGS remains unclear. It also remains unknown whether the dLGN is among the targets of the upper SGS direction-selective neurons.

Our findings demonstrate rich functional diversity among colliculogeniculate axons and reveal a wide range of preferences and selectivity for motion directions/axes, On/Off preference, and spatial and temporal frequencies of visual stimuli (Figure 1). Notably, a large fraction of colliculogeniculate boutons exhibit a high degree of direction selectivity, On-Off receptive fields, and/or small receptive fields, indicating that these SC neuron types indeed constitute a significant fraction of the stellate cells projecting to the dLGN. We also directly imaged visual responses in hundreds of superficial SC neurons labeled by Rorβ-Cre, a mouse line specifically labeling SC stellate cells and local inhibitory horizontal cells^54^ (Figure S9). Consistent with findings in colliculogeniculate axons, we observed a high level of functional diversity and enrichment in high motion selectivity, On-Off preference, and compact receptive fields among Rorβ-Cre-positive neurons in the upper SGS (Figure S9).

The direction selectivity in sSC neurons originates from the input provided by direction-selective ganglion cells^78^. Consistent with the nonlinear effect of spike thresholding, the direction selectivity calculated from neuronal spiking was greater than that from membrane potential changes in most recorded SC neurons^78^. Our observation that Rorβ-positive colliculogeniculate axons and neurons had right-shifted DSI and ASI distributions compared to retinal axons aligns with this previous report^78^. Direction-selective ganglion cells also innervate the shell of the dLGN. A combination of direction-selective retinal input with collicular input that is further sharpened through signal convergence and non-linear transformation in the SC can result in enhanced direction selectivity in dLGN neurons compared to using retinal input alone. Since visual signals in SC neurons are derived from an integration of retinal input, local amplification, and local inhibition, diversified or even novel receptive field properties may be created in the SC and then routed to the dLGN. Future studies could explore these possibilities by incorporating additional types of visual stimuli, such as chirp stimuli and natural images/movies^79–83^.

### Common Logics for Organizing Excitatory Inputs in the dLGN

Visual centers, from the retina to higher-order visual cortices, maintain a retinotopic arrangement to retain the mapping of neighboring visual space in closely-situated neurons. However, beyond the retina, retinotopic organization experiences scattering on fine spatial scales, perhaps due to the extensive spatial reach of axon arbors and stochastic factors during development. Both retinal and collicular axons are organized by retinotopy, and they exhibit highly consistent axes and rates of retinotopic progression in the same dLGN regions (Figure 2). Moreover, these axons display similar degrees of scatter along both the azimuth and elevation axes (Figure 2). These results suggest that non-retinal inputs, such as colliculogeniculate axons, also undergo highly regulated retinotopic refinement during development, and the mechanisms instructing retinogeniculate axon refinement likely extend to colliculogeniculate axons.

Below the spatial scale of retinotopic scatter, we previously discovered a fine-scale functional logic for arranging nearby retinal boutons according to their visual feature preferences^59^. We reported that nearby retinal boutons have a strong tendency to share similar feature preference(s) when they are less than ∼10 um apart^59,84^—the same spatial scale of retinal bouton clusters on proximal dendrites of dLGN neurons^57,58^. Collicular boutons can intermingle with retinal boutons and synapse on the same proximal dendrites of dLGN neurons, suggesting their participation in the same bouton clusters^18^. Consistently, we observed cluster-like structures among collicular boutons and between retinal and collicular boutons in the dLGN (Figures 3 and 4). Using pairwise and group-wise analyses similar to those applied to retinal boutons, we observed a high degree of similarity in feature preference(s) among nearby collicular boutons from different axons and between nearby collicular and retinal boutons. These findings demonstrate that the same fine-scale functional logic that organizes retinal axons also governs the convergence among collicular boutons and between retinal and collicular boutons, likely representing a general principle for arranging excitatory inputs in bouton clusters in the dLGN.

Although we did not directly record the dendrites postsynaptic to the neighboring axonal boutons, previous electron microscopy (EM) analysis demonstrated that anatomical proximity strongly predicts the convergence of retinal boutons onto the same dendritic domains, with a ∼50% likelihood for boutons within 6 μm to converge onto the same target neuron^59^. This high convergence rate is likely due to the nature of individually wrapped bouton clusters and the spatial separation between clusters. Since collicular boutons intermingle with retinal boutons and participate in bouton clusters^18^ (Figure 4), the likelihood of nearby retinal boutons synapsing onto the same dendritic domain likely applies similarly to collicular boutons and to the co-clustering between collicular and retinal boutons. The presence of the other ∼50% non-converging nearby boutons could add random noise and reduce the average similarity indices. Nevertheless, feature preference similarity was significantly above chance among boutons within a distance of 6 μm, strongly supporting the selective convergence between neighboring collicular boutons and between neighboring retinal and collicular boutons.

### Shaping Early-Stage Visual Processing by Non-Retinal Input

Cortical feedback and collicular input are two main sources of excitatory non-retinal input to the dLGN, and they exhibit significant differences in both morphology and function. While axons from layer 6 V1 neurons target distal dendrites of thalamocortical neurons with small boutons, collicular input innervates proximal dendrites with their intermediate-sized boutons interspersed among retinal boutons^14,18,85–87^ (Figure 4A). Consistent with their distinct anatomy, cortical feedback has been reported to modulate the response gain but not directly contribute to dLGN motion selectivity: silencing V1 neurons resulted in mixed effects on the response amplitudes to drifting gratings across mouse dLGN neurons without altering their orientation selectivity^41,81^, and suppressed dLGN responses to naturalistic movie clips through a predominantly divisive effect^81^. In contrast, our findings showed that silencing colliculogeniculate input significantly reduced motion selectivity in temporal-motion-selective or horizontal-axis-selective dLGN neurons and decreased the number of these neurons. The strong impact of collicular axons on dLGN motion selectivity supports their ‘driver-like’ roles, as suggested by brain slice recordings^18^.

Enhanced responses to motion may help animals better detect approaching predators and navigate their environment. Compared to retinogeniculate axons, colliculogeniculate input comprises a larger fraction of DS boutons, exhibits higher motion selectivity, and is enriched with direction-selective and axis-selective axons that encode motion along the temporal direction or the horizontal axis, respectively. Although visual signals conveyed by collicular input may be slightly delayed by a few milliseconds compared to retinal input^88^, the delay is much shorter than the time it takes for an object to move across the receptive field of a dLGN neuron (a few hundreds of milliseconds), allowing collicular input to still strongly affect motion processing in dLGN neurons and their downstream cortical circuits.

### Technical Considerations

Rorβ-Cre labels stellate cells, a subset of narrow-field cells and horizontal cells in the SC, and these cells could provide local and long-range output to other SC neurons and several subcortical nuclei^54,55^. Although chemogenetic silencing of Rorβ-Cre+ cells could affect those non-dLGN neurons, our main findings are unlikely to be impacted. This is because the colliculogeniculate projection is directly silenced by hM4Di, and other subcortical nuclei that are ipsilaterally innervated by the Rorβ-Cre+ SC neurons—such as the vLGN, LP, and PBG^55^—do not significantly innervate the ipsilateral dLGN^89^.

Moreover, Rorβ-Cre labels a small number of neurons in the retina (data not shown) and AAV injected into the SC may be taken up by retinal axons and retrogradely label RGCs^11^. However, we did not observe significant retrograde RGC labeling within six weeks of virus injection into the SC, consistent with the findings of limited retrograde labeling of RGCs when AAVs encoding Cre-dependent proteins were injected into the SC^11^. In a separate control experiment, we injected AAV2/2-FLEX-GCaMP6f into the contralateral eye of Rorβ-Cre mice to assess whether RGC axons were labeled, and no fluorescent signals were detected in the dLGN when examined in vivo (data not shown). Nevertheless, we limited the imaging of collicular axons and silencing of collicular neurons to within six weeks of viral injection.

The reduction in motion selectivity in dLGN neurons following collicular input silencing was primarily caused by extensive suppression of responses in the preferred direction. This ‘selective suppression’ contrasts with the ‘iceberg effect’, which would increase motion selectivity instead^90^. The observed selective suppression cannot be explained by the potential saturation of calcium signals in the preferred direction before silencing the SC, as any signal saturation would reduce the proportion of suppression in the peak direction relative to other directions in dLGN neurons after silencing the SC. Our simulation suggested that selective suppression could be attributed to collicular input with higher direction selectivity than the converging retinal input (Figure 7). However, our model may be limited by its ‘linear’ assumption for synaptic integration. Alternatively, converging retinal and non-retinal signals may be non-linearly amplified in postsynaptic neurons^91^. Future experiments will be necessary to confirm the nature of signal integration in dLGN neurons.

### Perspectives of the Developmental Convergence of Retinal and Collicular Axons

How retinal and collicular axons refine their convergence during development remains of great interest. It remains unknown whether retinal axons establish their synaptic clusters first, with collicular axons following, or if both coordinate the development of their fine-scale organization simultaneously. Additionally, the roles of spontaneous activity and visual experiences in guiding this developmental process are unclear. Retinal axons arrive at the dLGN before birth. Initially, retinogeniculate axons are guided by molecular cues and then instructed by spontaneous retinal waves before eye-opening around postnatal day 14 (P14)^92–94^. Initial projections of colliculogeniculate axons are also guided by molecular cues, such as the transcription factor Rorβ^55^. Collicular axons are detectable in the dLGN by P4/P5, with a sharp increase in axon density by P7^55^. Before eye-opening, retinal waves synchronize local activity between presynaptic retinal axons and postsynaptic neurons, guiding the formation of initial receptive field properties in the dLGN and SC^95,96^. The spontaneous activity of colliculogeniculate and retinal axons originating from the same group of RGCs should thus be highly synchronized. This synchronization likely establishes the similar retinotopic organization of retinal and collicular axons in the dLGN. The fine-scale arrangement of axonal boutons according to their common visual feature preferences likely occurs after retinotopy refinement, as the spatial scale of axonal bouton clusters is below the retinotopic scatter (Figure 2 and 3). Retinal axon arbors become more complex in the SC earlier than in the dLGN^93^, suggesting that colliculogeniculate neurons may establish visual receptive field properties earlier, allowing for subsequent co-refinement of their axon targeting with retinogeniculate axons. Furthermore, the positions of retinal boutons are adjusted along the axon arbor to form bouton clusters between P12 and P20^97^, indicating that the formation and refinement of retinal and collicular bouton clusters may continue after eye-opening. New imaging strategies are needed to assess axon activity and dynamics in the dLGN of young mice to reveal the timing and mechanisms underlying this fine-scale refinement of retinal and collicular boutons.

## ACKNOWLEDGMENTS

We thank Chinfei Chen, Michael Crair, Jonathan Demb, Chen Li, Chang Liu, In-Jung Kim, and Yao Xue for providing critiques of the manuscript, and members of the Liang and Crair labs for discussions. We thank Elise Savier for teaching us the SC cranial window implant. Support was provided by the National Institutes of Health (NIH) (R01 EY034697 to L.L.), grants from the Smith Family Foundation, the Whitehall Foundation, the E. Matilda Ziegler Foundation, the Klingenstein-Simons Fellowship Award in Neuroscience, and the Lawrence Young Investigator program (to L.L.), Yale College Dean’s Research Fellowship and Benjamin Franklin College Richter Fellowship (to M.L.), WTI Undergraduate Fellowship (to D.G.).

## AUTHOR CONTRIBUTIONS

YF and LL designed the experiments. YF and LL developed the dual-color imaging and associated image analysis protocol. YF performed surgeries and imaging and analyzed the data. ML and AO performed surgeries. DG and WH assisted in image analyses. YF and LL wrote the manuscript.

## STAR METHODS

### CONTACT FOR REAGENT AND RESOURCE SHARING

Further information and requests for resources and reagents should be directed to, and will be fulfilled by, the Lead Contact, Liang Liang.

## EXPERIMENTAL MODEL AND SUBJECT DETAILS

### Animals

Rorβ-Cre (stock number 023526) and Vglut2-Cre (stock number 016963) were purchased from the Jackson Laboratory. All animal care and experimental procedures were approved by the Yale Institutional Animal Care and Use Committee (IACUC) and federal guidelines. Animals were housed with standard mouse chow and water provided *ad libitum*. Both male and female mice were used in all studies; no sexual dimorphisms were observed.

## METHOD DETAILS

### Viral injections

To label retinal ganglion cell axons, 1 µL of AAV2/2.CAG.GCaMP6f.WPRE.SV40 (Addgene #100836; packaged by Yale Vision Core) was gently injected intravitreally into the right eye after the mice were anesthetized by a mixture of ketamine (100 mg/kg) and xylazine (10 mg/kg). Special care was taken during the injection procedure to minimize bleeding and prevent cataract formation. Confirmation of RGC infection was obtained through histological assessment.

To label colliculogeniculate neurons and axons, mice were anesthetized with isoflurane in 100% O_2_ (induction, 3%–5%; maintenance, 1%–2%). Subcutaneous infiltration with lidocaine (less than 7 mg/kg) was performed five minutes before making the first skin incision. A total of 600 nL of AAVs was stereotaxically injected into the left SC at four locations in the area 0.7-1.0 mm lateral and -0.15-0.85 mm posterior to Lambda, and 1.1-1.4 mm ventral to the skull. For dual-color calcium imaging of both retinogeniculate and colliculogeniculate axons, we injected AAV2/1.Syn.Flex.NES.jRGECO1a.WPRE.SV40 (Addgene #100853-AAV1) into the SC of Rorβ-Cre mice two weeks after the intravitreal injection of AAV-GCaMP6f. In separate cohorts of Rorβ-Cre mice, we injected AAV2/1.hSyn.Flex.GCaMP6f.WPRE.SV40 (Addgene #100854-AAV1) into the left SC to compare the visual response properties of collicular axons labeled by jRGECO1a and by GCaMP6f, or to measure visual response properties of Rorβ-Cre+ SC neurons. To assess the causal influence of colliculogeniculate input, we injected AAV2/9.hSyn.DIO.hM4D(Gi).mCherry (Addgene #44362- AAV9) or AAV2/1.hSyn.DIO.mCherry (Addgene # 50459-AAV1; control for hM4D(Gi) manipulation) into the left SC of Rorβ-Cre or Vglut2-Cre mice.

To label neurons in the dLGN, a total of 120 nL of AAVs was stereotaxically injected into the left dLGN at two locations. The injection coordinates were 2.25-2.33 mm lateral and 2.3-2.7 mm posterior to Bregma, and 2.6-2.8 mm ventral to the skull. To label dLGN neurons in Rorβ-Cre mice, a 1:1 mixture of AAV2/1.hSyn.Flex.GCaMP6f.WPRE.SV40 (Addgene #100837-AAV1) and AAV2/1.hSyn.Cre (Addgene #105553-AAV1) was injected. To label thalamocortical cells in Vglut2- Cre mice, AAV2/1.hSyn.Flex.GCaMP6f.WPRE.SV40 (Addgene #100837-AAV1) was injected.

### Headpost and cranial window implant for dLGN imaging

A headpost and cranial window were implanted 1-3 weeks after viral injection, using previously reported methods^59^. Mice were administered 0.06 ml of dexamethasone sodium phosphate (4.8 mg/kg) approximately 2 hours prior to surgery to reduce brain edema. The mice were anesthetized with isoflurane in 100% O_2_ (induction, 3%–5%; maintenance, 1%–2%) and placed on a heating pad (CWE) in a stereotaxic apparatus (KOPF). Ophthalmic ointment (Vetropolycin) was applied to their eyes. Buprenorphine extended-release injectable suspension (3.25 mg/kg; Fidelis Animal Health) was administered. Subcutaneous infiltration with lidocaine (less than 7 mg/kg) was performed five minutes prior to making the first skin incision. A two-pronged headpost was attached to the skull, centered approximately 2.7 mm lateral and 2.1 mm posterior to Bregma over the left hemisphere and oriented tangentially to the curved skull surface. The head was then tilted to secure the headpost in clamps (Thorlabs) that aligned the headpost parallel to the stereotaxic apparatus platform. A 3-mm diameter craniotomy was performed at the center of the headpost. Cortical and hippocampal tissue underneath was gently aspirated to reach the thalamus surface while preserving the thalamic surface and optic tract. A 3 mm x 3.4 mm (diameter x height) stainless steel cylindrical cannula (MicroGroup) with a 3-mm diameter coverslip glued to the bottom by UV-cured Norland Optical Adhesive 71 was stereotaxically inserted into the craniotomy and lowered approximately 2.75 mm below the skull’s surface to slightly press against the thalamus surface. The cannula was affixed to the skull with C&B Metabond (Parkell). To facilitate imaging with a water-immersion objective and light shielding, a low-profile adaptor was created by gluing a neodymium ring magnet (Indigo® Instruments, outer diameter, inner diameter, height: 7.5 mm, 5 mm, 1 mm) to the skull around the cannula. During two-photon imaging sessions, this ring magnet held the light shielding in place by contacting an 8 mm x 0.3 mm (diameter x height) spring steel round shim (McMaster) attached to the blackout fabric (Thorlabs). Finally, meloxicam (0.5 mg/kg) was administered, and the mouse was allowed to recover.

### Headpost and cranial window implant for SC imaging

A headpost and cranial window were implanted 1-3 weeks after viral injection, using previously reported methods^64^. The mice were anesthetized with isoflurane in 100% O_2_ (induction, 3%–5%; maintenance, 1%–2%) and placed on a heating pad (CWE) in a stereotaxic apparatus (KOPF). Ophthalmic ointment (Vetropolycin) was applied to their eyes. Buprenorphine extended-release injectable suspension (3.25 mg/kg; Fidelis Animal Health) was administered. Subcutaneous infiltration with lidocaine (less than 7 mg/kg) was performed five minutes prior to making the first skin incision. A ∼4-mm diameter craniotomy was performed around the Lambda. A customized coverslip with four layers of glass glued together by UV-cured Norland Optical Adhesive 71 was inserted into the craniotomy and pushed against the transverse sinus anteriorly, exposing the SC for optical imaging^64^. A two-pronged headpost was attached to the skull, centered around Lambda over the left hemisphere and oriented tangentially to the curved skull surface. Both the coverslip and headpost were affixed to the skull with C&B Metabond (Parkell). Finally, meloxicam (0.5 mg/kg) was administered, and the mouse was allowed to recover.

### Two-photon calcium imaging

Two-photon calcium imaging was conducted on awake, head-restrained mice after they were habituated to head restraint while on a running wheel. Imaging of the dLGN was performed using a resonant-scanning two-photon microscope (Neurolabware), with a 20x, 1.0 NA, 5.6 mm WD objective (Zeiss) at 2 x (∼308 x 419 µm^2^) – 3.4 x (∼181 x 246 µm^2^) digital zoom, or a 10x, 0.6 NA, 8 mm WD objective (Olympus) at 4 x (∼308 x 419 µm^2^) – 5.7 x (∼216 x 294 µm^2^) digital zoom. SC soma imaging was performed using a resonant-scanning two-photon microscope (Bruker), with a 16x, 0.8 NA, 3.0 mm WD objective (Nikon) at 2.5 x (∼308 x 419 µm^2^) digital zoom. Light shielding was positioned around the objective to prevent the monitor light from reaching the sample. For dLGN imaging, fields of view (FOVs) were imaged at depths ranging from 80-150 µm below the surface of the optic tract (roughly corresponding to the upper 20-90 µm of the dLGN shell; high-quality images could be obtained throughout the upper ∼140-150 µm of the dLGN, data not shown). For SC imaging, FOVs were imaged at depths 50-130 µm below the SC surface. Dual-color calcium imaging of retinal boutons and collicular boutons in the same dLGN region was achieved by simultaneously imaging both GCaMP6f and jREGECO1a at 1000 nm using a tunable laser (80 MHz; Insight X3, Spectra-Physics). Emitted signals were detected by two GaAsP PMTs after a 562nm longpass dichroic mirror, a 510/84 bandpass, and a 607/70 bandpass filter, respectively. GCaMP6f signals during one-color calcium imaging were acquired at 960 nm. The laser power measured at the objective’s front aperture through the cannula ranged from 11 to 40 mW. Images were collected at a rate of 15.6 frames/s, 796 × 512 pixels/frame, using the Scanbox software (Neurolabware), or at a rate of 15.1 frames/s, 512 × 512 pixels/frame, using the Prairie View software (Bruker). Each imaging run lasted approximately 30 minutes, and an imaging session consisted of 4-8 imaging runs. The same FOV was imaged throughout a session. When acquiring data from different FOVs from the same mouse, the FOVs were at least 15 µm apart along the dLGN depth for imaging axonal boutons or at least 25 µm apart for imaging somas. Two-photon imaging experiments typically started one week after headpost and cranial window implant, continued daily for one to two weeks, and were completed within 6 weeks of AAV injection into the SC. No significant retrograde labeling of retinal ganglion cells from injection in the SC was observed within this time window.

### Pupil videography

During two-photon imaging, a pre-selected region of interest around the eye facing the LCD screen, including the pupil area, was recorded using a CCD camera (Mako U-130B) at 15.5Hz, synchronized with two-photon image acquisition (Scanbox, Neurolabware). The pupil was illuminated by the two-photon excitation infrared light emanating from within the brain.

### Visual Stimulation

Visual stimuli were generated using Psychtoolbox-3^98^, and displayed on a luminance-calibrated LCD monitor (Dell, 17”,1280 x 1024 pixels, 60 Hz refresh rate) placed 23 cm from the mouse’s right eye and spanning 80° × 70° of visual space (azimuth: 3° – 83°; elevation: -10° – 60°). We estimated the mean luminance of the LCD to be 24.5 lux on the screen and 16.4 lux to the mice eyes. The brightness of all stimuli was gamma-corrected.

To map the retinotopy of the imaging field, we utilized a binarized version of a bandpass-filtered noise stimulus at 80% contrast. This noise stimulus had a spatial frequency corner at 0.05 cycles per degree (cpd), a cutoff of 0.32 cpd, and a temporal frequency cutoff of 4 Hz^99^. The visual stimulus was presented within 5° x 40° bars, arranged vertically at one of 8 azimuth locations, and arranged horizontally at one of 8 elevations. The stimuli were presented for a duration of 2 seconds each, with a 2-second inter-stimulus interval at the mean luminance level. Visual stimulation also included a blank condition (mean luminance). The order of the stimuli was randomized within a single repeat, which consisted of a single presentation of each stimulus condition. A total of 20 repeats were delivered during one imaging session.

To assess visual tuning properties during two-photon imaging, we used full-screen sine-wave drifting gratings at 80% contrast. These gratings were presented at one of eight directions of motion, spaced 45° apart. For imaging collicular and retinal axons, drifting gratings were delivered at temporal frequencies of 0.5, 2, and 8 Hz and spatial frequencies of 0.02, 0.08, and 0.32 cpd. For imaging dLGN neurons after saline injection and after clozapine N-oxide (CNO) injection within the same imaging session, drifting gratings were delivered at a fixed temporal frequency of 2 Hz and spatial frequencies of 0.08 and 0.32 cpd. Each stimulus was displayed for 2 seconds, followed by a 2-second inter-stimulus interval of the gray screen at the mean luminance level. A single repeat of the experiment involved the presentation of drifting gratings of all the conditions in a random order. Typically, a total of 24 repeats were included in a single session for imaging retinal and collicular axons; and a total of 50 repeats were included 20 minutes after saline injection and 20 minutes after CNO injection, respectively, for imaging dLGN cells.

In a two-day paradigm of silencing colliculogeniculate neurons using hM4Di, we imaged the same field of view of the dLGN on two consecutive days. On the first day, a total of 12-16 repeats of the full-screen sine-wave drifting gratings (temporal frequencies of 0.5, 2, and 8 Hz and spatial frequencies of 0.02, 0.08, and 0.32 cpd; 80% contrast) were given before saline injection and 20 minutes after saline injection, respectively. On the second day, the same ROI was imaged with the same set of drifting gratings before CNO injection, and 20 minutes after CNO injection.

To assess the variability of single-trial mean responses, we displayed full-screen sine-wave drifting gratings at 80% contrast in eight directions (spaced 45° apart), with a temporal frequency of 2 Hz and a spatial frequency of 0.08 cpd. Typically, 50 repeats were included in a single session.

To evaluate the responses to luminance increment and decrement (On or Off responses) and to map the receptive fields more precisely, we used flashing square stimulation^100^. During this process, a black or white square at 80% contrast, measuring 5° x 5° in visual angle, was flashed on a gray background. The square was presented for 1.5 seconds per trial in an 8 x 8 grid that covered an area of 40° x 40° in visual angle. Each repeat included 64 white squares, 64 black squares, and 2 blank trials with only the gray background, all interleaved randomly. This stimulus set was repeated ten to twenty times for each FOV.

### CNO preparation and administration

To assess the effect of suppressing colliculogeniculate input, we administered the same volume of saline and CNO via intraperitoneal (ip) injection while the mouse was awake and head-restrained on a running wheel during two-photon imaging sessions. To prepare the stock solution of CNO, 10 mg of CNO dihydrochloride (Tocris Bioscience) was diluted with 0.825 mL ddH2O to achieve a final concentration of 10 mg/mL. 150 uL of 1 μg/μL CNO working solution was freshly prepared by mixing 15 μL of the CNO stock solution aliquot with 135 μL of saline. 125 μL of the CNO working solution was administered to achieve a final concentration of 5 mg/kg in each mouse^101^.

### Histology and immunohistochemistry

Mice were anesthetized with Euthanasia III solution (>150 mg/kg) by intraperitoneal injection and transcardially perfused with phosphate buffered saline (PBS) immediately followed by 10% formalin. Brains were post-fixed overnight at 4°C and subsequently transferred into 20% sucrose solutions. 40 μm cryosections were made with a microtome (Thermo Scientific HM430). Simultaneously, retinae were fixed with 10% formalin overnight and subsequently dissected. Brain sections or retinae were washed in PBS, before incubation in a blocking solution containing 5% normal donkey serum (NDS) (Sigma Aldrich D9663-10ML), 0.4% Triton X-100 (Sigma) in PBS (0.4% PBST) and 0.02% NaN3 for 60 min at room temperature. Brain sections or retinas were incubated with primary antibodies in the blocking buffer for 3-7 days at 4°C. After three washes with 0.4% PBST buffer, brain sections or retinae were incubated with secondary antibodies in the 0.4% PBST buffer containing 0.02% NaN3 for 2-5 days at 4°C. After three washes in PBS, brain sections were mounted in VECTASHIELD Antifade Mounting Medium with DAPI. Retinae were stained in 300 nM DAPI solution (Invitrogen D1306) before being mounted in VECTASHIELD Antifade Mounting Medium without DAPI (VECTOR LABORATORIES). Images from a single Z plane or automatically tiled z stacks were collected using confocal microscopes (Zeiss), including LSM 800, LSM 880 or LSM 900, with 10x, 20x, or 63x objectives. The primary antibodies used were chicken anti-GFP (1:1000, Invitrogen, A10262; used for GCaMP6f), rabbit anti-RFP (1:1000, Rockland, 600-401-379, used for jRGECO1a and mCherry), guinea pig anti-RBPMS (1:500, PhosphoSolutions, 1832-RBPMS), guinea pig anti-VGLUT2 (1:500, Synaptic systems, 135404), and mouse anti-VGAT (1:200, Synaptic systems, 131011). The secondary antibodies used were donkey anti-chicken Alexa Fluor 488 (1:800, Life Technologies, 703-545-155), donkey anti-rabbit Alexa Fluor 594 (1:800, Life Technologies, 711-585-152), donkey anti-mouse Alexa Fluor 647 (1:500, Life Technologies, A-31571), goat anti-guinea pig Alexa Fluor 594 (1:800, Life Technologies, A-11076), and goat anti-guinea pig Alexa Fluor 647 (1:800, Life Technologies, A-21450). Histology was performed typically within 6 weeks after AAV infections. No significant retrograde labeling of retinal ganglion cells from injection in the SC was observed within this time window under all experimental conditions.

## QUANTIFICATION AND STATISTICAL ANALYSIS

### Image processing

#### Image preprocessing

To correct for x-y motion along the imaged plane, a series of image registration and data-cleaning steps were applied. Initially, the movies captured on each imaging day were registered to a common average field-of-view using efficient subpixel registration methods^102^. Following registration, the movies were spatially downsampled by a factor of 2 and temporally downsampled by a factor of 5. Subsequently, denoising was performed using principal component analysis (PCA) on the concatenated movies across the entire imaging session. By performing PCA, the spatial principal components with the highest eigenvalues were identified. These components typically represented pixels with signal variations across time due to neural activity rather than photon shot noise. Each image could then be characterized by a weighted sum of these principal components. We used only the first 400 principal components (with the highest eigenvalues) out of a total of approximately 30,000 total to reconstruct the registered and downsampled movie while removing shot noise^103^. Note that this PCA de-noising procedure was solely used to improve the image warping coregistration steps that followed. A local image normalization method was applied to each frame to normalize the fluorescence intensity across boutons and increase the contrast between boutons and neuropil. After normalization, image warping was implemented using the ‘imregdemons’ function (MATLAB) to align all images with a new common average field of view. The pixel-wise displacement obtained from the imregdemons function was then spatially upsampled by 2 and applied to the original, subpixel-registered movies without PCA de- noising. Subsequently, a second round of image de-noising, local normalization and warping, was applied to the full-size processed movies. The newly-computed pixel-wise displacement was then applied to the aligned movies from the first round of image warping. Following these image registration and warping steps, no obvious x-y motion was observed. As a final step, PCA de-noising was performed for the third time. Notably, while the use of PCA de-noising did increase the signal-to-noise ratio and consequently the yield of usable boutons or cells, the observed results were not dependent on this operation (data not shown).

For preprocessing recordings of dLGN neurons, the aforementioned procedure was only implemented up to the second round of PCA denoising, as no discernible x-y motion was observed at this stage already.

#### Green and Red channel co-registration

Co-registration methods were used to ensure the alignment of the red channel (jRGECO1a) and green channel (GCaMP6f) for the same field of view. We co-registered the two channels by applying the pixel-wise displacement obtained from efficient subpixel registration and warping of the green channel to both the green and red channels, frame by frame.

#### Green and Red channel de-mixing

As the emission spectra of GCaMP6 and jRGECO1a had a minor overlap, the signals detected by both the green- and red-channel PMTs had bleed-through signals from the other channels. We developed an algorithm to linearly de-mix the fluorescence signals in both channels. Specifically, we formulated the recorded fluorescence in the red and green channels to be as follows:

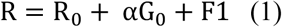

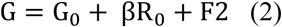

R and G represented the recorded fluorescence in the red and green channels, respectively; R_!_ and G_!_ represented the real fluorescence in the red and green channels; α and β were the green-into-red leakage ratio and red-into-green leakage ratio; F1 and F2 represented the background constant (dark noise) in the red and green channel. R_0_ and G_0_ could be derived by combining equations (1) and (2):

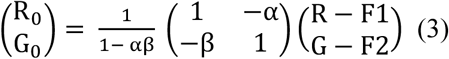

We derived α, β, F1, and F2 under two special cases, when R_0_ = 0 and when G_0_ = 0. When R_!_ = 0, equations (1) and (2) led to,

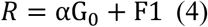

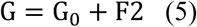

By subtracting equation (4) from equation (5), we derived

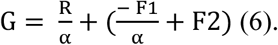

On the other hand, when R_0_= 0, R had contributions from the bleed-through signals from G_0_and constant dark noise F1 and therefore reached the minimum value for any given G. In the scatter plot of G as a function of R, the data points with minimal R values were located at the left bound of the plot. To identify these data points, we sorted the G values in ascending order and divided the lower 95% of the values into equal-sized bins with 30 values per bin. Within each bin, we selected the data point with the minimum R value. We then performed linear fitting of these data points:

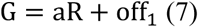

Comparing formulas (6) and (7), we derived the relations,

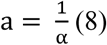

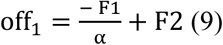

In a similar manner, when G_0_ = 0, we derived

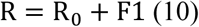

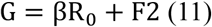

The combination of (10) and (11) led to

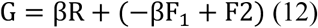

Such relations occurred when G reached its minimal values at the lower bound of the G∼R scatter plot. Similarly, we identified the minimal G in each bin of R values and conducted Linear fitting of these lowest data points to derive b and off_)_:

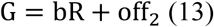

Comparing equations (12) and (13), we derived the relations

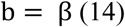

By solving (8) (9), (14) and (15), we obtained the values of α, β, F1 and F2.

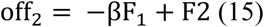

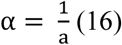

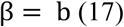

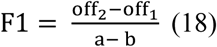

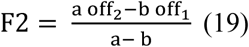

Finally, we solved equation (3) with the results of (16)-(19) to obtain G_0_ and R_0_, the real green and red signals. For the videos of each imaging session, we derived α, β, F1, and F2 using the average green images and average red images of the first 500 frames of the session. We then applied these parameters to each single frame of the videos from the entire session to de-mix red and green signals. Also see Figure S1A-D.

#### Bouton mask identification

We previously developed an automated image segmentation algorithm to identify boutons and extract masks for further signal processing^59^. First, an average image of absolute change in fluorescence (ΔF/F) was calculated for each trial type. This was done by averaging the evoked response maps for each condition across all N repetitions of that trial type. The formula used was |mean((F_i_-F_i0_)/F_i0_)|, i=1..N, where F_i_ represented the mean fluorescence during stimulus presentation and F_i0_ was the average baseline fluorescence during the 1 second prior to each stimulus onset. By using the absolute value of ΔF/F, we included boutons showing strong suppression by visual stimuli. Next, we applied a bouton identification procedure to each of these ΔF/F images independently. To start with, we performed local normalization by subtracting the local mean from each pixel and then dividing the difference by the local variance, with the local mean estimated through isotropic filtering of the image using a Gaussian kernel with a standard deviation of 3.4 µm. The local variance, on the other hand, was estimated using a larger Gaussian filter with a standard deviation of 60 µm. Morphological filters were then applied as follows to identify connected sets of pixels that together resembled the size and shape of a typical RGC bouton. First, small pixel gaps were filled by interpolation using a square-shaped structuring element of 1-by-1 µm. Next, we removed all small, unconnected structures via an ‘opening’ operation using the same structuring element. To obtain candidate masks, we first binarized the above images by setting pixels with values above 10% to 15% of the maximal pixel amplitude after filtering to ‘1’ while setting all other pixels to ‘0’. A Euclidian distance transform was then applied to these binary images (MATLAB built-in function ‘bwdist’). The watershed transform (MATLAB built-in function ‘watershed’) was used to finalize the segmentation. The results from the distance transform and watershed transform of individual ΔF/F images were combined by summing the distance transform across conditions. This value was then normalized by the bouton count obtained from the watershed transform. A final watershed transform was applied to this normalized distance image to increase the accuracy of the procedure and reduce the risk of false positives in the bouton identification procedure. Note that the bouton masks obtained from watershed transform did not overlap; moreover, we made sure any two bouton masks identified in the same imaging channel were separated by at least one line of pixels with the value ‘0’ by setting the adjacent pixels belonging to different bouton masks to 0. We extracted bouton masks from the green channel and red channel independently. We discarded any collicular bouton masks if their overlap with retinal bouton masks exceeded 30% of the collicular bouton’s total area or retinal bouton’s total area. In addition, to remove residual calcium signals not originating from the bouton itself, we estimated neuropil masks as circular annuli of 3 µm width, with the inner edge located 2 µm beyond the edge of a corresponding bouton mask. Pixels from adjacent bouton masks were excluded from these neuropil masks.

#### Cell mask identification

The automatic extraction of masks for dLGN or SC cell bodies was performed using a custom implementation of a deep-learning-based neuron segmentation procedure^104^. All cell masks were manually verified and adjusted as necessary to ensure accuracy and reliability. To eliminate background calcium signals not originating from the cell body, neuropil correction was performed. Neuropil masks took the form of circular annuli with a width of 9 µm, positioned with their inner edge 4 µm beyond the outermost edge of the corresponding cell body mask.

#### Time course extraction and correction

To obtain raw fluorescence traces for bouton masks, cell body masks, and neuropil masks, the fluorescence intensity value of a mask at each time point was defined as the average fluorescence across the pixels belonging to the mask.

To account for neuropil signals which might contaminate signals in the bouton or cell body trace, neuropil correction was applied by subtracting a scaled version of the corresponding neuropil trace (the scale factor was individually derived from linear regression between the minimal bound of the bouton or cell body trace and the corresponding neuropil trace) from each bouton or cell body trace^105^.

We also corrected baseline fluorescence F_0_ to remove the decay in fluorescence from activity evoked during the previous visual stimulus presentation^59^. Due to decay dynamics of GCaMP6f *in vivo* (as a result of buffering of calcium and the calcium indicator) after stimulus-evoked calcium activity, the GCaMP6f fluorescence sometimes did not fully return to baseline during the 1 second right after the offset of a previous stimulus presentation and persisted in the next 1 second used to calculate F_0_ for the following stimulus period. Therefore, a baseline correction was introduced that modeled this exponential decay of previously evoked GCaMP6f calcium activity during the 2-second inter-trial interval, F_prev-fit_(t) = a*exp(-t/𝜏)+c, where 𝜏 was the time constant and c was the corrected baseline F_0_. We used an experimentally determined fixed time constant of 1 second (in agreement with previously determined GCaMP6f dynamics *in vivo*^106^). This fitting procedure was independently carried out for each bouton or cell body in each single trial.

To assess the fractional change in fluorescence, ΔF/F_0_(t), following each visual stimulus presentation, the fitted, exponentially decaying contribution from the previous trial was first subtracted from F(t) during the 1-second interval prior to visual stimulus presentation and the 2-second interval during visual stimulus presentation. Then, the corrected baseline was used as the new baseline F_0_ to compute ΔF/F_0_(t). Single response values during a given trial were obtained by averaging the ΔF/F_0_(t) response during the 2-second stimulus window.

### Estimation of visual tuning preferences in boutons

#### Estimation of boutons with significant visual responses

We defined single-condition responses as the average from all the trials of a given stimulus condition. Visually-evoked responses were corrected by subtracting the average response across blank trials. There were approximately 20 repeats (or trials) of each noise bar position, 24 repeats of each combination of the 8 directions, 3 spatial frequencies (SF) and 3 temporal frequencies (TF) of drifting gratings, 50 repeats of each of the 8 directions at 0.08 cpd and 2 Hz, 10 or 20 repeats of luminance increments and decrements (‘ON’ and ‘OFF’), and 100 repeats of blank stimulation for each FOV in an imaging session. We determined a bouton to be significantly responsive under a stimulus condition for drifting grating stimuli or bandpass-filtered noise stimuli by requiring the amplitude of ΔF/F_0_(t) during the response window to exceed 2 standard deviations above or below the mean baseline activity (computed using the 1-second window before stimulus onset) for at least 8 out of 31 time points (15.5 Hz frame rate x 2 sec stimulus presentation). To determine whether a bouton had significant On or Off responses to flashing square stimuli, we utilized the average trace of the three square-stimulation locations (among all 64 white or black squares) evoking maximal or minimal bouton activity. If the ΔF/F_0_(t) of the maximal average trace during the response window exceeded 2 standard deviations above the mean baseline activity for at least 6 out of the 23 time points (15.5 Hz frame rate x 1.5 sec stimulus presentation), the bouton was considered to have significant positive On or Off response. If the ΔF/F_0_(t) of the minimal average trace during the response window exceeded 2 standard deviations below the mean baseline activity for at least 6 out of the 23 time points (15.5 Hz frame rate x 1.5 sec stimulus presentation), the bouton was considered to have significant negative On or Off response.

To assess if a given bouton exhibited a significant positive response at a particular spatial frequency and temporal frequency, we required that at least 3 out of the 8 directions at this spatial and temporal frequency evoked significant positive responses according to the criteria described above. A similar approach was used to determine if a bouton exhibited a significant negative response to a particular spatial and temporal frequency. Note that all boutons contributing to the main results (e.g., clustering of direction and axis selectivity) underwent additional quality controls (see below), ensuring that noisy boutons were excluded from our analyses.

#### Direction tuning curve fitting

For each collicular and retinal bouton showing a significant positive response at a given spatial and temporal frequency, a direction tuning curve was computed. The direction tuning curves were initially sampled in steps of 45°. To obtain a more precise estimate of the preferred axis and direction, a fitting approach was used to estimate the preferred direction. Tuning curves were fitted with a two-peaked Gaussian with offset^107^:

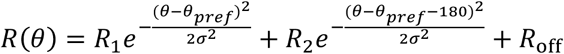 was the ΔF/F response for stimulus direction 𝜃 . This model assumed that the peaks of the two Gaussians were 180° apart. 𝜃*_pref_* was defined as the preferred direction evoking the strongest ΔF/F response, 𝑅_1_. 𝑅_2_ was the amplitude of the second peak located at 𝜃_pref_ + 180. It was also assumed that both Gaussians shared a common standard deviation, 𝜎. The fifth fitted parameter was a constant amplitude offset, 𝑅_off_.

Several steps were taken to improve the reliability of fitting direction tuning curves and to optimize the accuracy of estimating the preferred direction of motion^59^. To increase the number of input points for the fitting procedure from 8 to 25, a heuristic method of interpolation and extrapolation was implemented. First, a ninth point was added at 360°, which was identical to the one at 0°. Then, the number of input points was doubled from 9 to 17 by linear interpolation of the 9-point direction tuning curve. For the interpolated data point between the two most strongly driven initial directions (out of 9), we further adjusted the interpolated amplitude so that its value became a close approximation of that predicted point from a Gaussian curve fit through the rest of the points, thus reducing the error introduced by linear interpolation given the expected continuity of the curves. To this end, we applied the following empirical formula as described below. Note that our results were largely unchanged if this additional adjustment to the linear interpolation was omitted^59^. However, this additional peak adjustment resulted in significantly smaller residual values between the fitted curve and the initial 8- direction tuning curve.

The interpolated amplitude between the two most strongly driven initial directions was calculated as follows. 𝑅_S1_was defined as the strongest response out of all 8 directions and 𝑅_S2_as the stronger of the two responses for directions ± 45° adjacent to 𝑅_S1_. 𝑅_S3_ was defined as the weaker of the two responses adjacent to 𝑅_S1_. 𝑅_S4_ was defined as the response adjacent to 𝑅_S2_, at 90° from 𝑅_S1_. The interpolated ΔF/F response 𝑅_S12_ between 𝑅_S1_ and 𝑅_S2_ was defined as: 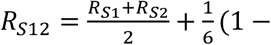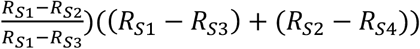. This method compared the slope between 𝑅_2"_ and 𝑅_2)_ with the slope between 𝑅_2"_ and 𝑅_23_. If the peak was flat, a maximum amount of 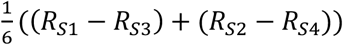 was added, corresponding roughly to the expected value of a Gaussian peak. If the absolute values of the slopes between 𝑅s_1_ and 𝑅_S2_ and between 𝑅_S1_ and 𝑅_S3_ were identical (and therefore 𝑅_S1_ was the real peak of the Gaussian), this corresponded to 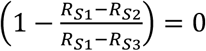 in the above equation, therefore resulting in no additional value added to the interpolation method. A similar method was used to interpolate negative peaks.

To further improve the stability of the fitting procedure and to better approximate the direction tuning curve, we added two shadow-copies of the two-peaked Gaussian function, circularly shifted by +360° and -360°:

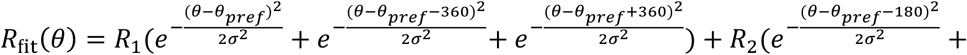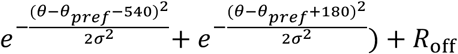 This addition of the shadow-copies increased the range to [-90°, 450°], and thus extended the fitted tuning curve by 4 additional directions (at 22.5° spacing) on either end. While the adjusted linear interpolation and the addition of the shadow copies improved the fitting procedure, similar results were obtained using the basic 17-point linearly interpolated tuning curve (data not shown).

A bootstrapping method involving random sampling of trials from each condition was then implemented to fit the tuning curves. Specifically, for each of the 100 iterations, the tuning curve was initially computed by randomly sampling (with replacement) and averaging responses from 24 trials sampled from each of the 8 directions. These 8-point tuning curves were then interpolated, extended, and finally fitted using the method described above. The final parameters used were the mean of the fitted parameters across the 100 sampling iterations.

To determine if the fitting procedure yielded high-quality fits, a combination of criteria was used. Each iteration of the fitting procedure yielded a coefficient of determination *r*^2^, defined as the explained variance using least-squares regression to fit the data. As a second control step, a combined coefficient of determination 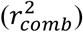 was computed by comparing the original direction tuning curve with the fitted curve derived from the average of each fitting parameter (across 100 iterations). To assess both the quality and the reliability of the fitting procedure, we introduced a heuristic goodness of fit, 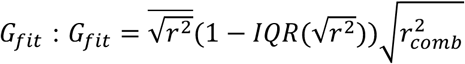, where IQR was defined as the interquartile range – the difference between the 75^th^-percentile and the 25^th^-percentile (of 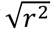 values across iterations). A bouton was considered to have a well-fit direction tuning curve at a given spatial frequency if the goodness of fit, 𝐺_fit_, was greater than 0.66. The threshold was chosen based on examination of a large proportion of example boutons, and values in the range of 0.5 to 0.9 yielded similar results. The complete direction curve fitting procedure was run separately for each spatial and temporal frequency combination (up to nine combinations).

#### Axis and direction selectivity

For each bouton or cell body, we calculated a ‘vector sum’ axis selectivity index (ASI; i.e., selectivity for a motion along a given axis) on each interpolated direction tuning curve^106^. This index was calculated by projecting the ΔF/F response for each of the 16 directions (after interpolation) in the range between 0° and 360° onto a circle with 2i progression and estimating the magnitude of the normalized vector sum, which ranged from 0 to 1 (maximum selectivity): 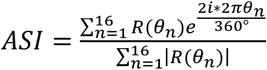. Opposite directions contributed in an additive fashion, while orthogonal directions canceled each other out. The ASI computation was iterated 100 times by bootstrapping and averaged for each spatial and temporal frequency. To obtain a final ASI estimate, the mean ASI was computed from the spatial and temporal frequency that evoked a peak response for the bouton or cell body if the response was significantly positive for the given spatial and temporal frequency.

In a similar manner, we computed a ‘vector sum’ direction selectivity index (DSI), by projecting the 16 directions onto a circle with 1i progression: 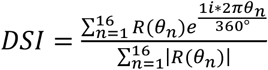. As with ASI, the DSI estimate was repeated with the bootstrapping method, and a final DSI estimate was computed as the mean DSI across 100 iterations of bootstrapping from the spatial and temporal frequency that yielded the peak and significant positive visual response.

#### Functional classification of axonal boutons and dLGN neurons

A bouton or neuron was classified as ‘direction-selective’ if, for the spatial and temporal frequency that yielded the strongest responses, (i) it exhibited significant positive responses at that particular spatial frequency and temporal frequency, (ii) the positive peak was at least 3 times greater than the negative peak across the 8 directions of stimuli, (iii) the tuning curve was successfully fit, with a goodness of fit greater than 0.67, (iv) the DSI exceeded 0.2 (a value equivalent to 0.33 if direction selectivity was calculated as 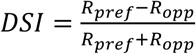, where 𝑅*_pref_* was the response in the preferred direction and 𝑅*^opp^* was the response in the opposite direction).

A bouton or neuron was defined as ‘axis-selective’ (i.e., most strongly responsive to motion along two opposite directions constituting a single axis of motion) if, for the spatial and temporal frequency that yielded the strongest responses, (i) it exhibited significant positive responses at that particular spatial frequency and temporal frequency, (ii) the positive peak was at least 3 times greater than the negative peak across the 8 directions of stimuli, (iii) the tuning curve was successfully fit (goodness of fit greater than 0.67), (iv) the DSI was below 0.2, and (v) the ASI exceeded 0.15 (a value equivalent to 0.33 if axis selectivity was calculated as 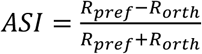, where 𝑅*_pref_* was the response in the preferred direction and 𝑅_orth_ was the mean response in the two directions orthogonal to the preferred one). We distinguish the term axis selectivity from orientation selectivity, as the latter is often used even for responses to stationary (i.e., non-drifting) oriented gratings – a stimulus not examined in this study. It is possible that certain axis-selective boutons may not be strongly driven by stationary gratings.

A bouton or neuron was defined as ‘broadly-tuned’ if, for the spatial and temporal frequency that yielded the strongest responses, (i) it exhibited a significant positive response, (ii) the ASI and DSI were below 0.15 and 0.2, respectively; (iii) the positive peak was at least 3 times greater than the negative peak across the 8 directions of stimuli.

A bouton or neuron was defined as ‘suppressed’ if it had significant negative responses for at least two spatial frequencies and no significant positive responses at any spatial frequency.

Boutons that showed significant visually-evoked responses but were not classified into any of the above conservatively defined categories were labeled as ‘unclassified’ and were not included in subsequent analyses.

Finally, a small proportion of candidate bouton masks were not significantly driven by any of the presented visual stimuli. These were classified as ‘unresponsive’ and not considered further.

#### Preferred direction of motion and preferred axis of motion

The preferred direction was defined for direction-selective boutons or neurons by taking the fitted 𝜃_./01_ values from the spatial and temporal frequency that evoked the peak response of the bouton. Estimates of the preferred direction ranged from 0° to 360°.

The preferred axis of motion was defined in a similar fashion, for both axis-selective and direction-selective boutons or neurons, by taking the preferred axis of motion from the spatial and temporal frequency that had the strongest response. Estimates of the preferred axis of motion ranged from 0° to 180°.

To compensate for the pitch angle of the mouse head during imaging acquisition (Pitch of 31.6° above the angle estimated for a typical ambulatory position^108^; Roll: 16.7°; measured using stereotaxic coordinates during headpost implant, N=3 mice), we rotated the preferred direction or axis of motion counterclockwise by 31.6°.

#### Preferred spatial and temporal frequency

For simplicity of presentation, we converted the spatial frequency of stimulation to integer numbers

(with two-octave spacing between integers) using the formula 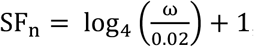, with 𝜔 being defined as the spatial frequency in cycles per degree. SF_"_, SF_)_, and SF_3_, respectively, corresponded to actual stimulus spatial frequencies of 0.02, 0.08, and 0.32 cycles/degree. Similarly, we converted the temporal frequency of stimulation by the formula TF_G_ = log_4_(2δ) + 1, with δ being defined as the temporal frequency in Hz. TF_"_, TF_)_, and TF_3_, respectively, corresponded to actual stimulus spatial frequencies of 0.5, 2, and 8 Hz.

For a given bouton, we selected the preferred direction as the direction evoking the strongest ΔF/F response among all eight directions. We first listed the ΔF/F response in the preferred direction with three spatial and three temporal frequency combinations in a 3x3 matrix. Next, the number of input points increased from 3x3 to 6x6 by linear interpolation (MATLAB function ‘imresize’) to improve the quality of the Gaussian fitting. Subsequently, we fitted a 2D Gaussian model on the 6x6 matrix (MATLAB function ‘lsqcurvefit’). If the 𝑟^)^ of fitting was larger than 0.3, we used the center of the fitted Gaussian to determine the spatial and temporal frequency preference. If the 𝑟^)^ was below 0.3, we estimated the preferred spatial frequency and temporal frequency as the combination evoking the strongest ΔF/F response within the 3x3 matrix by the original value.

#### On/Off preference index

Responses to flashing square stimulation were analyzed to determine the On/Off preference index. If the bouton had a significant positive On or Off response (see Subsection ‘Estimation of boutons with significant visual responses’), we used the average trace of the three square-stimulation locations (95 percentile of all 64 white or black squares) that evoked maximal activity to calculate the On/Off preference. If the bouton had a significant negative On or Off response, we utilized the average trace from three locations of white or black square stimuli evoking minimum activity (5 percentile of all 64 white or black squares), respectively. An On/Off preference index was calculated using the averaged response traces to luminance increments (On stimulus with white square stimuli) and luminance decrements (Off stimulus with black square stimuli). A positive response to only On, only Off, or an equally positive response to both On and Off corresponded to index values of 1, -1 or 0, respectively. Boutons lacking both a significant On response and a significant Off response were not considered. In addition, boutons that were significantly suppressed by an On stimulus were defined as Off-responsive, while boutons that were significantly suppressed by an Off stimulus were defined as On-responsive.

In order to take into account the dynamics of the evoked On and Off responses, a weighted On/Off preference index was introduced as follows: 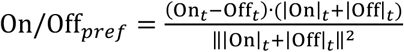 On_t_and Off_t_ were defined as the On and Off response time courses during the 1.5-second response window. In this equation, the term (On_t_ – Off_t_) determined the sign of the index at each timepoint. The dot product between this term and (|On|_@_ + |Off|_@_) was used to assign a relative weight to each timepoint according to its summed response magnitude. Then the numerator was normalized to obtain a single preference index between -1 and 1.

#### Coefficient of variation

To measure the variability of single-trial mean responses during the 2-second visual stimulation in retinal boutons, collicular boutons, and dLGN neurons, we calculated the coefficient of variation (CV) as follows:

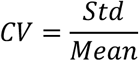

Here, 𝑆𝑡𝑑 represents the standard deviation of the single-trial mean responses across all fifty repeats under the preferred direction of drifting gratings (0.08 cpd, 2 Hz). 𝑀𝑒𝑎𝑛 represents the average response across these same fifty repeats.

#### Estimation of retinotopic preferences in axonal boutons

##### Retinotopic tuning curve fitting

Two retinotopic tuning curves were independently fit for each bouton: one for tuning along the azimuth axis and one for tuning along the elevation axis. Both curves consisted of eight evenly spaced values, each representing the average response across trials for a given location in the visual space of the oriented bar containing binarized spatiotemporal noise. Tuning curves were approximated using a Gaussian function: 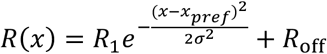 The ΔF/F response, 𝑅(𝑥), varied as a function of the retinotopic stimulus location 𝑥. The maximum response, 𝑅_1_ + 𝑅_off_, was evoked at 𝑥_pref_, the preferred retinotopic location. The standard deviation 𝜎 of the Gaussian was proportional to the receptive field size along this axis. To increase the number of points for fitting from 8 to 15, an interpolation method similar to the one used for fitting direction tuning curves was implemented. As responses were very reliable and well fit, no bootstrapping method was implemented. A retinotopic response was considered significant if 2 out of the 8 directions showed a significant response (see Subsection ‘Estimation of boutons with significant visual responses’) and the 𝑟^2^ of fitting exceeded 0.64. Retinotopic tuning curve fitting was also implemented for the neuropil rings surrounding each bouton, to estimate the local retinotopic preference in the field of view.

#### Retinotopic map estimation

Each pixel in the field of view was assigned a preferred retinotopic location by first designating the center of each neuropil ring with its preferred retinotopic location, then dilating by a disk of 13 µm radius from each neuropil center, respectively, and averaging the preferred retinotopy across overlapping disks. The final pixel-wise estimates of retinotopic preference were obtained by spatial smoothing using an isotropic two-dimensional Gaussian filter with a standard deviation of 4 µm.

The rate of change of retinotopy along the field of view (which was tangential to the surface of dLGN) was measured along the axis for which the retinotopic map changed the fastest. To compute this spatial axis, we first calculated the two-dimensional pixel-wise gradient: 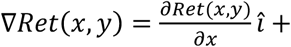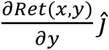. The spatial axis was defined as the normalized mean gradient vector across pixels, 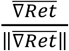. The smoothed retinotopic map was then projected onto the normalized mean gradient vector (i.e., onto the unit vector along the direction of maximal change in retinotopic preference). For each pixel, we derived its projected location along this new spatial axis as: 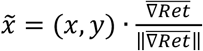. The relationship between the preferred retinotopic location and 𝑥a was modeled according to a linear function with offset: 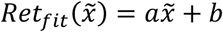. The fitted parameter 𝑎 (units: deg/µm) indicated the progression rate of the smoothed retinotopic map. We computed the normalized mean gradient axis and the scale factor, 𝑎, both for maps of azimuth and maps of elevation.

#### Fine-scale retinotopic scatter

Fine-scale retinotopic scatter (‘deg scatter’) was estimated as the absolute retinotopic deviation 𝑆_ret_(units: degrees of visual space): 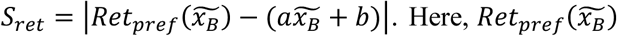 denoted the preferred retinotopic location of a given bouton, while 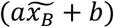 gave the predicted receptive field center based on the neuropil estimate, according to the projected location 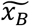 along the mean gradient axis. We also calculated the absolute deviation in spatial distance from the fitted spatial progression in the field *S^spa^* (‘distance scatter’, units: µm): 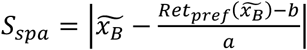. This value was equivalent to the distance that a bouton would need to be moved along azimuth or elevation in order to obtain a perfectly smooth map.

#### Receptive field fitting and area calculation

To determine the On or Off receptive field, we performed 2D Gaussian fitting of each bouton’s ΔF/F responses to 8x8 flashing black or white square stimuli, respectively. We excluded boutons with an 𝑟^2^ of fitting below 0.3 from further analysis, as they did not exhibit a significant or regular receptive field. For boutons with an 𝑟^2^ above 0.3, we used the center of the fitted Gaussian as the center of the receptive field. Moreover, the receptive field region was defined as the ellipse encompassing one standard deviation of the Gaussian function from the center. The area of the receptive field was then calculated using the equation: S = π𝑎b, where 𝑎 and 𝑏 represented the standard deviations of the 2D Gaussian, or the lengths of the semi-major and semi-minor axes of the ellipse, respectively.

#### Axon identification

Axon identification – the process of assigning boutons imaged in a field of view to the same axon – was carried out on data from blank trials (up to 200 trials per session) to avoid assessment of pairwise correlations during periods of visual stimulus presentation^59^. We assumed that boutons from the same axon would share spontaneous calcium events at a substantially higher frequency than pairs of boutons not belonging to a common axon. To identify spontaneous calcium events, the activity time courses of all blank trials were first concatenated to obtain a single ‘spontaneous’ activity trace for each bouton (15-30 minutes duration). We focused on periods containing significant spontaneous events for each bouton, as these periods were robust to sources of noise. To this end, we thresholded each spontaneous trace by 3 standard deviations above and below the mean activity. To identify entire events, including the baseline before event onset, time points in an interval of 700 milliseconds before and 700 milliseconds after each thresholded event were included. Finally, we concatenated these peri-event time courses of spontaneous activity for each bouton. All boutons exhibited some significant spontaneous activity and were thus included in the axon classification procedure.

The concatenated time courses of spontaneous events for a given bouton were then cross-correlated with the time courses of all other boutons during the same epochs. In this way, we created a matrix of Pearson correlation coefficients between all pairs of boutons in the field of view. To obtain a sparse matrix only populated by large values, the original matrix was further thresholded: the correlation coefficients for a given bouton were maintained if the coefficients were larger than 0.7 or if they exceeded 2 standard deviations above the mean value of all the coefficients between this bouton and all others. If neither of these conditions were met for a given bouton pair, the associated correlation coefficient was set to 0. The cosine similarity between every pair of boutons was then computed from the thresholded matrix of Pearson correlation coefficients. Each bouton in a pair had an associated vector of rectified spontaneous activity pairwise correlation coefficients with all other boutons in the field of view, and the cosine similarity between two boutons reflected the cosine of the angle between these two vectors of correlation coefficients (Figure S2B). This step was important: if a small minority of pairwise correlation coefficients were low due to noise, while most other coefficients were high for other pairs of boutons belonging to the same axon, this procedure would help ensure that all boutons were nevertheless properly assigned to the same axon. Next, we classified bouton clusters using agglomerative hierarchical clustering based on the pairwise distance, computed as ‘1 – cosine similarity’. We defined ‘correlation similarity’ as ‘1 - cluster distance’, where the cluster distance was the distance between two groups, each consisting of one or more bouton. This distance was calculated using the weighted-pair group method with arithmetic means (WPGMA) algorithm. We chose a cutoff threshold of correlation similarity to classify two groups of boutons as belonging to a common axon. Specifically, two groups of boutons with correlation similarity exceeding a threshold of 0.1 were assigned to the same axon. This threshold was highly conservative, as it erred on the side of combining groups together. Thus, this procedure minimized the likelihood that pairs of boutons that actually belonged to the same axon would be assigned to two different axons and included in subsequent pairwise analyses in Figure 3. On average, the mean Pearson correlation coefficient increased monotonically with correlation similarity as defined above, and the correlation similarity threshold of 0.1 corresponded to a mean Pearson pairwise correlation coefficient of 0.52.

#### Inter-bouton feature comparison

The absolute difference in the preferred direction of motion ranged between 0° and 180° and was only computed for DS/DS pairs.

The absolute difference in the logarithm (base 4) of the preferred spatial frequency ranged from 0 to 2 (i.e. from 0 to 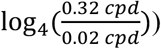, and was computed for all bouton pairs, within and across categories, if both boutons had a well-defined preferred spatial frequency.

The absolute difference in the logarithm (base 4) of the preferred temporal frequency ranged from 0 to 2 (i.e. from 0 to 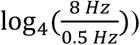, and was computed for all bouton pairs, within and across categories, if both boutons had a well-defined preferred temporal frequency.

The absolute difference in On/Off preference index ranged from 0 to 2 and was computed for all bouton pairs, within and across categories, if both boutons had significant On and/or Off responses.

#### Functional clustering

##### Functional clustering of bouton pairs

Functional clustering of bouton pairs was assessed by plotting the inter-bouton feature differences versus spatial distances between boutons belonging to different axons. A 4 µm sliding average was applied at 0.25 µm steps to these plots. The standard error of the mean was calculated by considering all pairs in a given 4 µm bin. We also computed the chance level, estimated after randomly permutating the feature differences across all bouton pairs spaced from 2 µm to 155 µm in the field of view while maintaining their spatial distances. This randomization was repeated 20 times. The standard error of the mean estimate for the permutated data was calculated for each randomization and then averaged across the 20 randomizations.

To normalize the above curves for comparison across different fields of view, we used a pairwise similarity index (𝑆𝐼) that compared the degree of actual similarity to the ‘null’ estimate of similarity using the above permutation procedure, as follows: 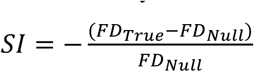. Here, the null functional distance 𝐹𝐷*_Null_* (the average absolute difference in feature preference across all permutated pairs within a defined inter-bouton distance) was subtracted from the true functional distance 𝐹𝐷_a/_0_ (the average absolute difference in feature preference across all pairs with inter-bouton distances in a defined range), followed by normalization. A similarity index value of 0 corresponded to similarity at chance levels, while an index value of 1 indicated identical feature preferences. Negative clustering indices were obtained if pairs were less similar than expected by chance. Clustering effects on a short spatial scale were assessed for pairs spaced 2 µm to 6 µm apart, while pairs spaced 2 µm to 155 µm apart served as a control for large-scale similarity or global bias.

##### Functional clustering of local groups of boutons

To quantify the functional clustering of nearby groups of boutons for each visual tuning feature, we computed a group similarity index for each visual feature and included all boutons within the vicinity of each given pixel in the image, assuming the local group of boutons met the following criteria: (i) the group of boutons within a 6 µm radius of the pixel must contained at least 3 AS or DS boutons which exhibited well-defined motion axis preferences, On/Off preferences, temporal frequency (TF), and spatial frequency (SF) preferences; (ii) the group of boutons must belong to at least two axons, with no more than 67% of the boutons belonging to any given axon; (iii) the group of boutons must contain both collicular boutons and retinal boutons. Pixels satisfying these topological criteria usually contained 3 to 12 acceptable boutons within the 6 µm radius. We then computed the group similarity index for the motion axis preference, On/Off preference, TF preference, and SF preference, respectively, among well-defined boutons near each pixel. To compute this group similarity index (SI), we first calculated (1 - standard deviation of On/Off preference indices), (1 - standard deviation of TE preference indices), (1 - standard deviation of SF preference indices), or the magnitude of the mean of polar unit vectors along preferred axes – a measure of groupwise circular variance in axis preference. We then normalized these values with the corresponding averages from 1000 shuffles according to (real SI - shuffled SI)/(1 - shuffled SI), where we shuffled the quadruplet of preference values (motion axis preference, On/Off preference, SF preference, and TF preference) among axons across all fields of view that contained the same number of boutons. A normalized similarity index was defined to be significant if it was larger than 95% of indices derived from shuffled data. For a group of boutons surrounding a given pixel to be considered as part of a combination- or relay-mode cluster, the similarity index for the corresponding visual feature (or features, for relay mode) had to be significant. For example, a group of boutons surrounding a given pixel was considered to be in an Axis-On/Off relay mode cluster if both its Axis similarity index and On/Off similarity index were significant. A group of boutons surrounding a given pixel was considered to be in an Axis combination mode cluster if the axis similarity index was significant but the On/Off similarity index were not. A group of boutons surrounding a given pixel was considered to be in a 3-feature relay cluster if Axis, On/Off and SF similarity indices; Axis, On/Off and TF similarity indices; Axis, SF and TF similarity indices or On/Off, SF and TF similarity indices were all significant. A group of boutons surrounding a given pixel was considered to be in a 4-feature relay cluster if Axis, On/Off, SF, and TF similarity indices were all significant.

To compute the frequency of each type of functional cluster, we calculated the number of unique pixels (i.e. the number of unique groups of boutons within a circle of 6 μm radius) belonging to each type of clusters and divided by the total number of unique pixels satisfying the aforementioned topological criteria. Quantification of these percentages for each type of cluster used the number of *unique* groups of boutons, as neighboring pixels could ‘double-count’ the same unique group of boutons.

#### Pupil size analysis

We used the pupil area to estimate the arousal level of the mouse. The pupil areas were automatically identified from the acquired videos using DeepLabCut^109^ with eight specifically-defined points surrounding the pupil. Subsequently, we used ‘fitEllipse’ function (Python 3) to fit the eight points and calculated the size of the fitted ellipse as the size of the pupil. We manually re-extracted the pupil if the likelihood of detection by DeepLabCut was less than 0.85, the distance of detection (defined as the average distance between the fitted ellipse and the eight detected points) was greater than 15 pixels, the difference of pupil areas in adjunct frames exceeded 150 pixels, or if the pupil area was greater than 8000 pixels or smaller than 50 pixels. We found that manual corrections were usually required for fewer than 1% of the frames. Subsequently, the pupil area time course was temporally smoothed by a median filter with a width of 3 adjacent frames.

For each trial, the average pupil area estimates from 1 s before to 2 s after the onset of visual stimulation were computed and used as the pupil area for that trial. To match the pupil size for the trials included for analysis after saline injection and after CNO injection, we established a threshold range of 20% - 80% of the pupil size across the entire imaging session. Any trials outside of this range were excluded from the comparison of visual responses after saline and CNO injections. Using 10% - 90% or 30% - 70% of the pupil size didn’t change the results in Figure S4.

#### Suppression index

In the one-day paradigm, we first performed pupil size matching before and after CNO injection. To assess the impact of silencing colliculogeniculate neurons on dLGN neurons, we defined a suppression index, SuI = [R_CNO_ – R_Saline_]/ [|R_CNO_| + |R_Saline_|], where R_CNO_, R_Saline_ represented the respective average response after CNO injection and after saline injection under the stimulation condition where the drifting gratings evoked maximum responses after saline injection. For neurons with negative averaged responses to drifting gratings, we reversed the sign of SuI. To determine the significance of the suppression effect (p < 0.05) for each neuron, we compared the responses following saline and CNO injections using the Mann-Whitney-Wilcoxon test. We only included those neurons with significant visually evoked responses across trials (See section: Estimation of boutons with significant visual responses) either after saline or after CNO injection in the later analysis.

In the two-day paradigm, we calculated the suppression indices on both days and compared whether the distributions of suppression indices were significantly different between the two days. We calculated the suppression index as SuI_Saline_ = [R_Saline_– R_Saline-Ctrl_]/ [|R_Saline_|+ |R_Saline-Ctrl_|] and SuI_CNO_ = [R_CNO_– R_CNO-Ctrl_]/ [|R_CNO_|+ |R_CNO-Ctrl_|], using the stimulation conditions evoking the peak activity for each neuron. R_Saline-Ctrl_ and R_CNO-Ctrl_ were the average responses across trials before saline or CNO injection, respectively; R_Saline_ and R_CNO_ were the single-trial responses after saline and CNO injection, respectively. For *Sup* neurons with negative averaged responses to drifting gratings, we reversed the sign of SuI. For each neuron, we determined whether the medians of the single-trial SuI_Saline_ and SuI_CNO_ were significantly different using the Mann-Whitney-Wilcoxon test (p < 0.05). We reported the average of SuI_Saline_ and SuI_CNO_ as the suppression index for each neuron in Figure S5D. For these analyses, we only included those neurons with significant responses (See section: Estimation of boutons with significant visual responses) in both imaging sessions.

#### Trial matching

To control for the potential influence of differences in pupil positions and pupil motion on the suppression index after silencing collicular input, we also calculated the suppression index using subsets of trials that were matched for both pupil position (both horizontal and vertical positions) and pupil motion (both horizontal and vertical motion, defined as the standard deviation of the time courses of horizontal and vertical displacement, respectively), during the 2-sec stimulus presentation for each trial. To achieve this, we randomly selected a trial after saline injection without replacement and paired it with a trial after CNO injection that had similar pupil positions and pupil motion, with the criteria that the difference in pupil positions had to be below 3 a.u. (∼1.5° of pupil rotation) and the difference in pupil motion had to be below 1 a.u (∼0.5°) between the two trials. If no after-CNO-injection trials fulfilled the criteria, the corresponding after-saline-injection trial was not considered. We continued the matching process until all the after-saline-injection trials were matched with after-CNO-injection trials. We then calculated the suppression index from all matched trials, excluding any unmatched trials. This matching procedure was repeated 50 times, with the order of selecting saline trials randomized each time. The final suppression index was calculated as the average of the results from these 50 repeats.

#### Classification of the suppression types

We classified the type of suppression as displacement, subtractive, divisive, or selective for each direction-selective or axis-selective neuron with response magnitudes significantly altered by silencing the colliculogeniculate input via CNO injection. First, we computed the direction tuning curve under the spatial and temporal frequency that yielded the maximum responses. A bootstrapping method involving random sampling of trials (using the MATLAB function ‘randsample’) with replacement from each of the eight directions was implemented to calculate the average tuning curve under each iteration. If the preferred directions were significantly different after saline and CNO injections, the neuron was considered to receive displacement suppression. If the preferred directions were not significantly different and the activity after CNO injection was higher than after saline injection in the preferred direction, the neuron was classified as ‘enhanced’. The remaining neurons were classified into subtractive, divisive, or selective suppression groups. The preferred directions of all tuning curves were aligned to 0°. Next, we performed bootstrapping to obtain the distribution of the suppression ratio (SR; defined as [R_CNO_ – R_Saline_]/ R_Saline_) in each direction. We then defined the 60^th^ percentile of the bootstrapped distribution to be the ‘high SR value’ and 40^th^ percentile of the distribution to be the ‘low SR value’. The null direction was defined as the direction with the minimum significant response (significantly larger than zero as determined by the student t-test), to avoid the directions that were no longer responsive under subtractive suppression. If the high SR value in the preferred direction was smaller than the low SR value in the null direction (and for AS neurons, we required the high SR value in the preferred direction to be smaller than both the null direction and the opposite null direction—e.g., if the null direction was -90°, the opposite null direction would be 90°), these neurons were classified into the subtractive suppression group. If the low SR value in the preferred direction was larger than the high SR value in the null direction and opposite null direction, these neurons were classified into the selective suppression group. The rest of the neurons were classified into the divisive suppression group.

#### Simulation of dLGN neurons from the tuning curves of collicular and retinal boutons

We simulated the tuning curves of individual dLGN neurons by linearly summing the weighted direction-selective (DS) collicular and DS retinal tuning curves. Given the rarity of displacement suppression after silencing collicular input and the tendency for nearby collicular and retinal boutons to share similar direction preferences, we assumed that the selected retinal and collicular tuning curves had the same preferred direction. Thus, all tuning curves were aligned with a preferred direction at 0°. Moreover, all tuning curves were normalized by their peak responses. The contribution ratio of the collicular tuning curve to the simulated dLGN neuron was determined by randomly sampling a number from a uniform distribution between 0 to 1. To account for slight variations in the contribution ratio due to random noise, we added noise from a normal distribution (mean = 0, sigma = 0.25) to the weights independently in all eight directions. Additionally, a noise factor from a normal distribution (mean = 0, sigma = 0.25) was added to the simulated dLGN tuning curve to mimic the viability of dLGN neuronal activity. Next, we simulated the collicular silencing by removing the collicular tuning curve from the linear model and adding newly sampled random noise. The dLGN neuron activity before and after collicular silencing was normalized by the maximum response amplitude. Subsequently, we performed bootstrapping and classified the simulated dLGN neurons according to the procedure described in the previous section.

We tested additional conditions to further verify the importance of the collicular DSI distribution obtained from our in vivo recordings. In the first condition, we simulated dLGN neurons using two retinal tuning curves to eliminate differences in the DSI distribution. Similarly, we simulated dLGN neurons using two collicular tuning curves, yielding results similar to the first condition (data not shown).

#### Statistics

Statistical tests were conducted using MATLAB. Non-parametric tests and parametric tests were used for comparing two independent groups (Mann-Whitney-Wilcoxon test or student t-test), two related groups (Wilcoxon signed-rank test), or multiple groups (Kruskal-Wallis test with *post hoc* Bonferroni correction or Dunn’s multiple comparisons test). To determine whether two populations have the same distribution of categorical variables, the Chi-squared test with *post hoc* Pearson residuals test was used^110^. P < 0.05 was considered significant. Additional details on sample sizes, statistical tests, and significant levels for each experiment can be found in figure legends, Results and Methods. All acquired data were included for analyses.

##### Linear mixed-effects model

To address the interdependence among dLGN neurons from the same animal, we examined the significance between various variables using linear mixed-effects models (’fitlme’ function in MATLAB)^111^. In each testing instance, we considered the animal (MouseNum) as a random effect and the type of injection (GroupNum; saline injection as 0 and CNO injection as 1) as a fixed effect. Meanwhile, we used (GroupNum|MouseNum) to nest the type of injection under the mouse ID. Finally, we wrote the function as ‘Data∼GroupNum+(1|MouseNum)+(GroupNum|MouseNum)’, where Data represents the neuronal activity. This model has the following mathematical form:

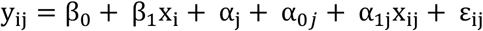

In this function, y_ij_ is the i^th^ observation for the j^th^ mouse; β_0_ is the fixed intercept; β_1_ is the fixed effect coefficient for x_i_; x_i_is the experimental group, either saline injection or CNO injection for the i^th^ observation; α_j_ is the random effect for the j^th^mouse, with the distribution α_j_ ∼ N(0, σ_1_^2)^); α_oj_are random effects, with the distribution α_ij_ ∼ N(0, σ_3_^2^);. α_ij_ are random effects, with the distribution α_ij_ ∼ N(0, σ_3_^2^); x_ij_ is the experimental group, either saline injection or CNO injection for the i^th^observation of the j^th^mouse; and ε_ij_represents the random error, with the distribution ε_ij_ ∼ N(0, σ^2^).

## DATA AND SOFTWARE AVAILABILITY

Requests for analyses and raw data on calcium imaging results may be made to the Lead Contact, Liang Liang

---

**Figure S1.**
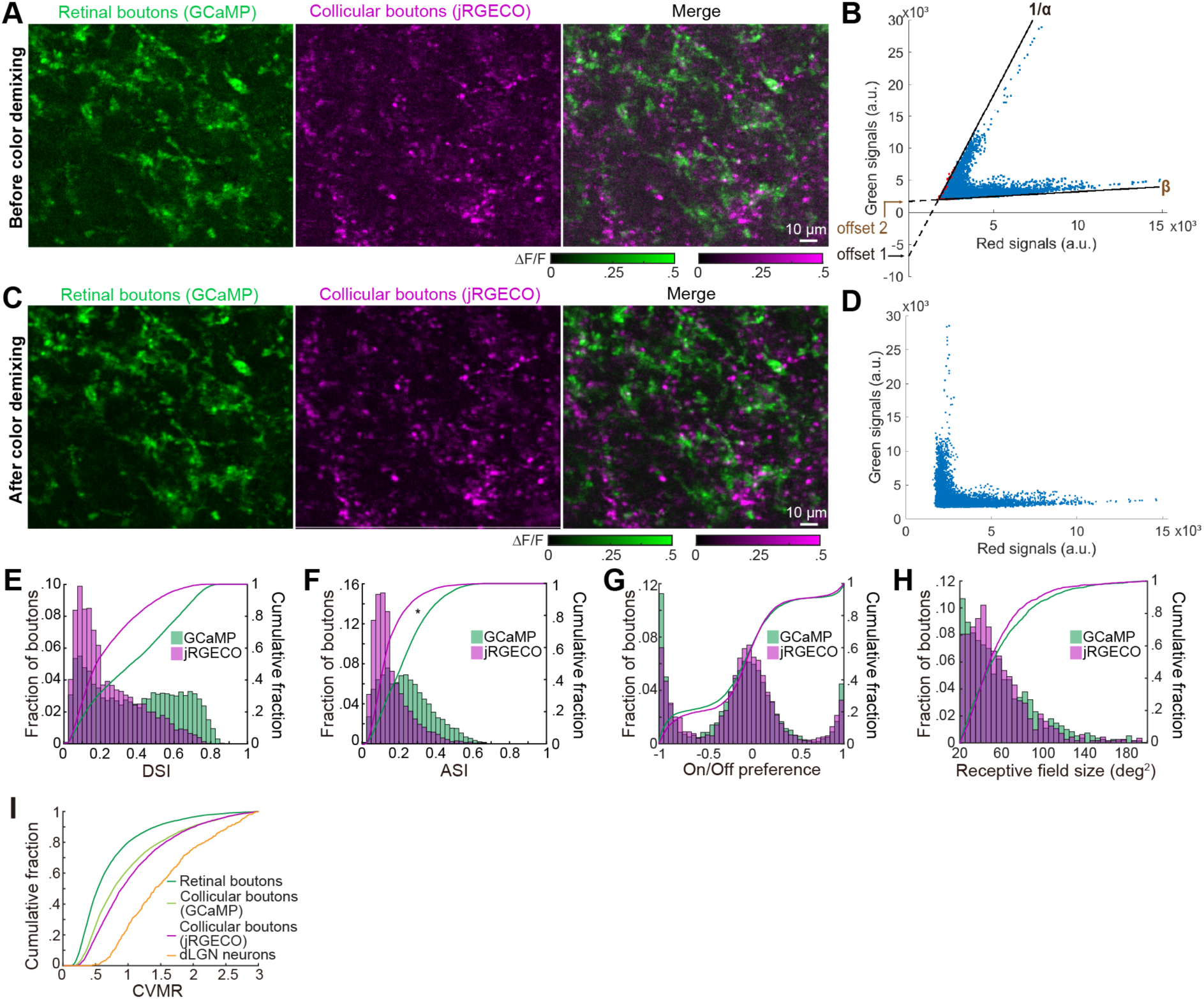
Color-de-mixing Method and Comparison of Visual Responses Between GCaMP-labeled and jRGECO-labeled Colliculogeniculate Boutons. (A) Visually evoked signals in retinal and collicular boutons before de-mixing the bleed-through signals between the red and green channels (due to the emission spectra overlap between GCaMP6f and jRGECO1a) in an example field of view (FOV) of the dLGN. ΔF/F: fractional change in fluorescence. (B) Scatter plot of the average raw fluorescence signals in the green channel against the red channel for each pixel in the FOV. α and β are the green-into-red and red-into-green leakage ratios, respectively. The slopes and offsets (1/α, offset 1 for green leakage, and β, offset 2 for red leakage) were fitted to the lowest envelope data points (labeled in red; bin size was 30 pixels), which primarily resulted from leakage from the other channel (see STAR Methods). (C) Visually evoked signals in retinal and collicular boutons in the same FOV of A after de-mixing GCaMP6f and jRGECO1a signals. (D) Scatter plot of the average raw fluorescence signals in the green channel against the red channel after color de-mixing. (E-H) Distributions of the DSI, ASI, On/Off preference, and receptive field size of GCaMP6f-labeled (green) and jRGECO1a-labeled (magenta) collicular boutons (DSI: p = 0.09; ASI, p = 0.02; On/Off preference, p = 0.36; Receptive field size, p = 0.76; linear mixed-effects model). GCaMP6f: N = 9292 boutons, 10 FOVs from 5 mice; jRGECO1a: N = 6455 boutons,18 FOVs from 8 mice. (I) Responses in dLGN neurons were more variable across trials than collicular and retinal boutons, and collicular boutons were more variable than retinal boutons (p = 0.005 between retinal boutons and GCaMP6f-labeled collicular boutons; p < 10^-7^ between retinal boutons and jRGECO1a-labeled collicular boutons; p < 10^-37^ between retinal boutons and dLGN neurons; p < 10^-5^ between jRGECO-labeled collicular boutons and dLGN neurons; p = 0.78 between GCaMP- and jRGECO-labeled collicular boutons). All the data are presented as mean ± SEM. *, p < 0.05; **, p < 0.01; ***, p < 0.001. E-I: linear mixed-effects model.

**Figure S2.**
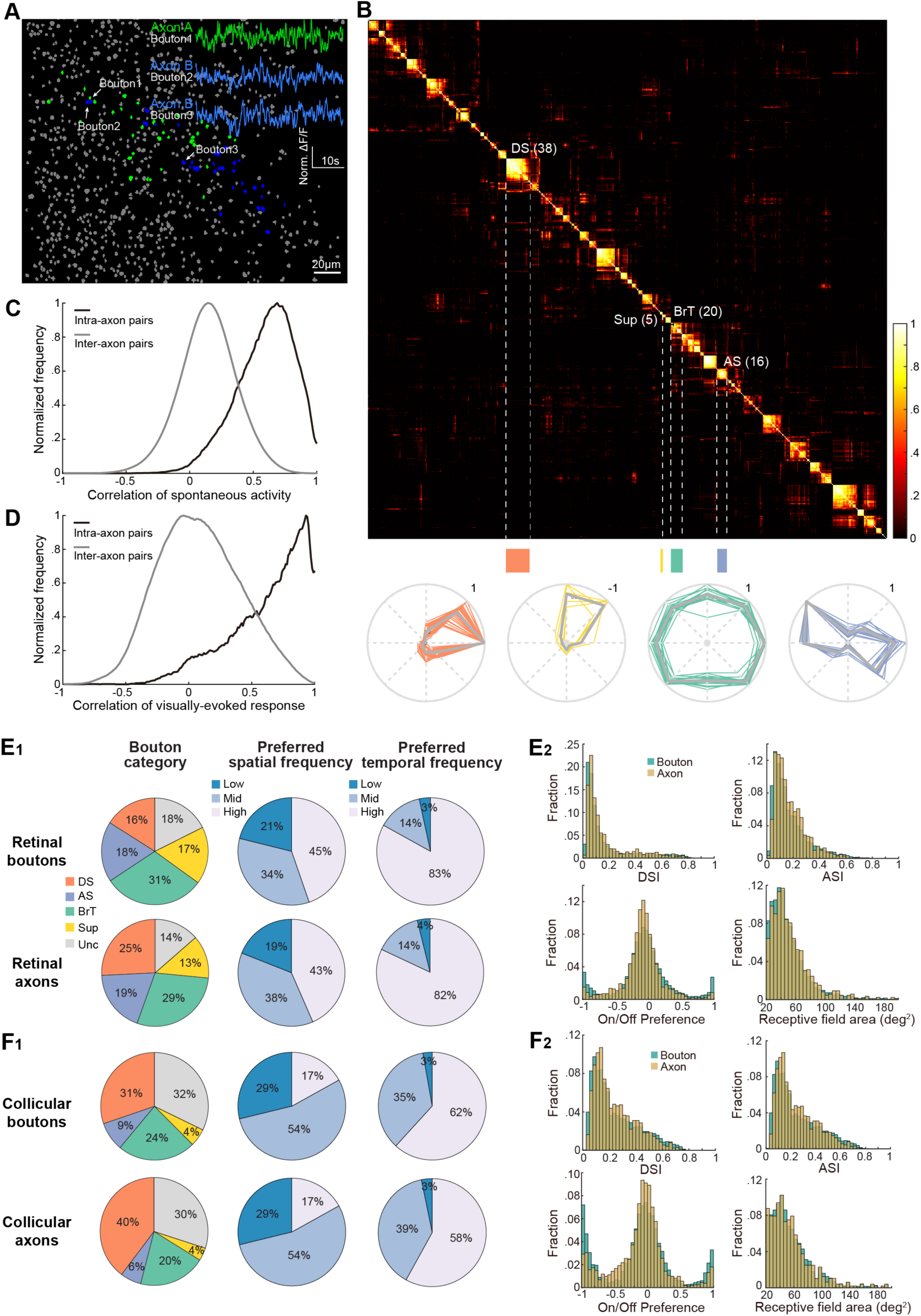
Assignment of Collicular Boutons to the Same Axon. (A) Boutons with highly correlated spontaneous activity were assigned to the same axon (e.g., Axon A in green; and Axon B in blue). Inset: An example pair of boutons assigned to the same axon (Bouton 2 versus 3) had high correlation in their spontaneous activity (Pearson correlation coefficient: 0.79) despite being approximately 80 μm apart, while nearby or distant boutons assigned to different axons showed lower pairwise correlation (Bouton 1 versus 2: 0.13; 1 versus 3: 0.11). (B) Top: Matrix of pairwise cosine similarity in spontaneous activity (a measure of the similarity between a pair of boutons for their correlation coefficients of spontaneous activity with all the other boutons in the FOV; see STAR Methods), sorted using hierarchical clustering. Four blocks of boutons assigned to four different axons are highlighted. The functional category and the number of boutons for each of the four blocks were labeled next to the block. Bottom: Peak-normalized direction tuning curves for all boutons from these four axons. (C) Distributions of pairwise correlation coefficients of spontaneous activity for inter- and intra-axonal bouton pairs. Note that a small subset of bouton pairs was still classified as belonging to the same axon despite the lack of strongly correlated activity, as these bouton pairs exhibited high similarity in their overall correlation patterns with all boutons in the field of view. Pairs of boutons with similar spontaneous activity were assigned to the same axon if the metric of pairwise similarity, ‘correlation similarity’ (derived from cosine similarity, see STAR Methods), exceeded a threshold of 0.1. This threshold was highly conservative, as it erred on the side of combining groups of boutons together. Thus, this procedure minimized the likelihood that pairs of boutons that actually belonged to the same axon would be assigned to two different axons and would thereby be included in subsequent pairwise analyses in Figures 3 and 4. (D) Similar to C, but based on visually-evoked activity. (E) E_1_: Distributions of retinal boutons and retinal axons (after classifying boutons into axons) across functional categories (p < 0.0001), spatial frequency preferences (Low, Mid, High: SF = 0.02, 0.08, 0.32 cpd, respectively; p = 0.0010), temporal frequency preferences (Low, Mid, High, TF = 0.5, 2, 8 Hz, respectively; p = 0.04). E_2_: Distributions of retinal boutons and retinal axons across DSI (p = 0.77), ASI (p = 0.72), On/Off preferences (p = 0.38), and receptive field areas (p = 0.88). N = 11319 retinal boutons, 2189 retinal axons, from 18 FOVs, 8 mice. (F) similar to E but based on collicular boutons and collicular axons across functional categories (p < 0.0001), spatial frequency preferences (p = 0.99), temporal frequency preferences (p = 0.0008), DSI (p = 0.22), ASI (p = 0.56, On/Off preferences (p = 0.65), and receptive field areas (p = 0.47). N = 6455 collicular boutons, 2938 collicular axons from 18 FOVs, 8 mice. *, p < 0.05; **, p < 0.01; ***, p < 0.001. E_1_, F_1_: Chi-squared test. E_2_, F_2_: linear mixed-effects model.

**Figure S3.**
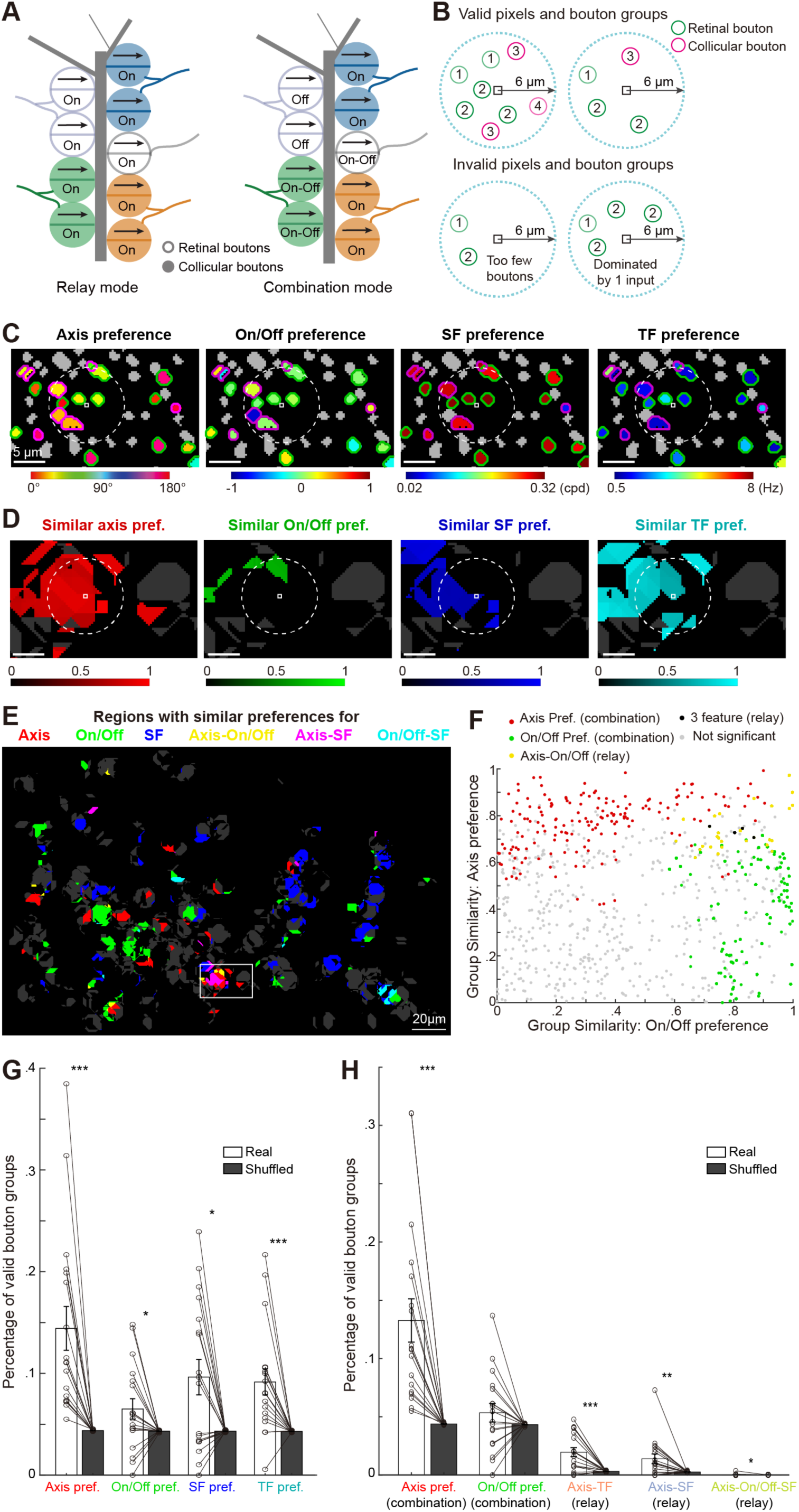
Relay-mode and Combination-mode Convergence of Nearby Retinal and Collicular Boutons. (A) Schematic illustrating ‘relay mode’ (nearby boutons exhibit similar preferences for multiple features, left) or ‘combination mode’ (nearby boutons share preferences for a single feature, right) between retinal and collicular boutons. (B) Schematic illustrating analyses of “groupwise” functional similarity. Analyses were restricted to pixels (black squares) whose 6-um radius circle (blue dotted line) contained three or more boutons and included at least one collicular bouton and one retinal bouton. We required that these boutons had well-defined preference estimates for motion axis (including both direction-selective and axis-selective boutons), On/Off, spatial frequency (SF), and temporal frequency (TF), and <67% of boutons in the group belonged to a single axon. Bouton numbering denotes their axon identity. (C) Feature preferences of boutons in an example subregion, colored by preferences for the motion axis, On/Off, SF, and TF, respectively. Gray boutons lack well-defined preference estimates. Retinal boutons are outlined in green and collicular boutons are outlined in magenta. (D) Pixel maps of group-wise similarity indices for motion axis preference, On/Off preference, SF preference, and TF preference for the same subregion in C. Only pixels with significant similarity index values (exceeding 95% of shuffled estimates) are colored, while other valid pixels are gray, and invalid pixels are black. Dashed white circles in C and D illustrate an example of a pixel surrounded by a local group of boutons with similar preferences for motion axis, SF, and TF, but not for On/Off. (E) Pseudocolored image illustrating various modes of convergence for groups of boutons in an example FOV (gray rectangle, subregion shown in C-D). Red, green, and blue: combination-mode groups with similar preferences for only one feature (motion axis, On/Off preference, or SF, respectively). Yellow, purple, cyan: relay-mode bouton groups with similar preferences for two or more features. (F) Groupwise similarity indices of the motion axis preference and On/Off preference for valid pixels in E. Bouton groups could exhibit similar preferences for motion axis only (“axis preference combination mode”), On/Off only (“On/Off preference combination mode”), both features (“Axis- On/Off relay mode”), or all 3 features (“3 feature relay”). Gray circles indicate groups that did not exhibit significant groupwise similarity (i.e., >95% of shuffled estimates) for either feature preference. (G) Percentages of valid groups of nearby boutons with significant similarity in specific feature preferences for each of the 18 FOVs. White bars, real data; black bars, randomly shuffled data. The statistic results are as follows: Axis preference similar group (real: 0.144 ± 0.021, shuffled: 0.043 ± 0.0001; p < 0.0001), On/Off preference similar group (real: 0.065 ± 0.010, shuffled: 0.043 ± 0.0002; p = 0.04), SF preference similar group (real: 0.097 ± 0.017, shuffled: 0.043 ± 0.0002; p = 0.03), TF similar group (real: 0.092 ± 0.013 shuffled: 0.043 ± 0.0001; p = 0.0004). (H) Percentages of valid groups of nearby boutons exhibiting specific combination-mode or relay-mode convergence for each of the 18 FOVs. White bars, real data; black bars, randomly shuffled data. The statistic results are as follows: Axis-only combination mode (real: 0.133 ± 0.019, shuffled: 0.044 ± 0.0002; p < 0.0001), On/Off-only combination mode (real: 0.054 ± 0.008 shuffled: 0.043 ± 0.0002; p = 0.21), Axis-TF relay mode (real: 0.020 ± 0.003 shuffled: 0.003 ± 0.0001; p = 0.0003), Axis-SF relay mode (real: 0.014 ± 0.004, shuffle: 0.0003 ± 0.0001; p = 0.004), Axis-On/Off-SF relay mode (real: 0.0003 ± 0.0001 shuffle: 0.0002 ± 0.00001; p = 0.03). Here, we only plotted the types of relay-mode convergence with the percentages of unique bouton groups significantly higher than the chance level. All the data are plotted as mean ± SEM. Wilcoxon signed-rank test was used for all tests in G and H. *, p < 0.05; **, p < 0.01; ***, p < 0.001.

**Figure S4.**
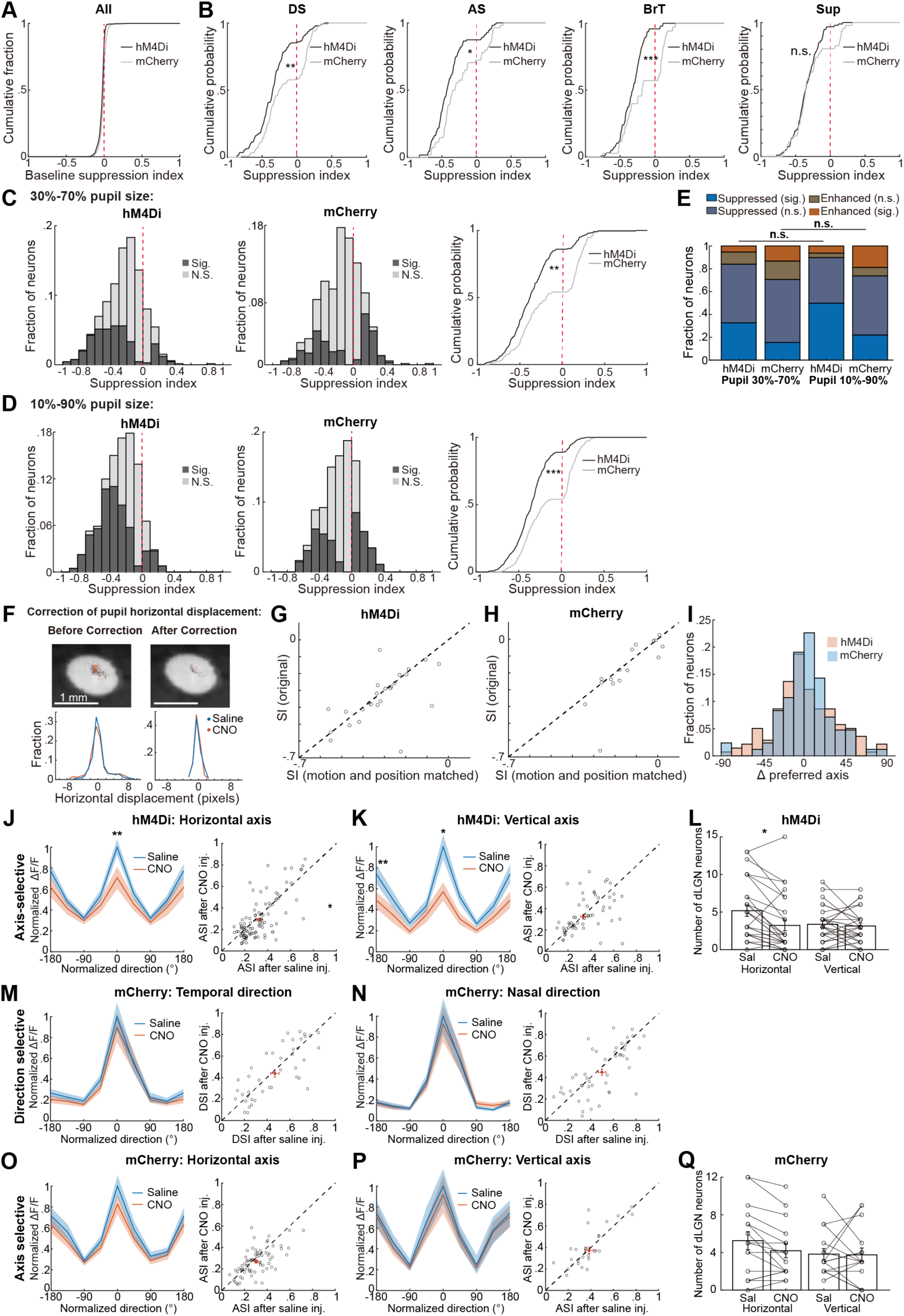
Effects of CNO Injection, Pupil Size and Displacement on dLGN Neurons. (A) Cumulative distribution of suppression indices of dLGN neuron baseline activity in hM4Di- and mCherry-injected Rorβ-Cre mice (hM4Di: 1070 neurons; mCherry: 944 neurons; p = 0.07, linear mixed-effects model). All panels in this figure: N = 27 FOVs from 11 hM4Di-injected Rorβ-Cre mice and 16 FOVs from 6 mCherry-injected Rorβ-Cre mice. (B) Cumulative distributions of visual response suppression indices of dLGN neurons in hM4Di- vs. mCherry-injected mice for different functional categories (DS, p = 0.007; AS, p = 0.013; BrT, p < 10^-3^; Sup, p = 0.27). (C) Distribution of visual response suppression indices of dLGN neurons after silencing the colliculogeniculate projection in hM4Di (left) vs. mCherry-injected mice (middle), when only trials with the pupil size within 30% - 70% of the whole pupil size pool were included. Right: Cumulative distribution of suppression indices of significantly impacted dLGN neurons in hM4Di-vs. mCherry-injected mice (p = 0.0011). (D) Distribution of visual response suppression indices of dLGN neurons after silencing the colliculogeniculate projection in hM4Di (left) vs. mCherry-injected mice (middle), when trials with the pupil size within 10% - 90% of the whole pupil size pool were included. Right: Cumulative distribution of suppression indices of significantly impacted dLGN neurons in hM4Di vs. mCherry-injected mice (p = 1.0*10^-5^). (E) The stacked bar plot summarizing the distribution results in in C and D. Altering the pupil size threshold didn’t change the result that dLGN neurons in hM4Di-injected mice were suppressed more strongly than in mCherry-injected mice (hM4Di Pupil 10-90% vs hM4Di Pupil 30-70%: p = 0.89; mCherry Pupil 10-90% vs mCherry Pupil 30-70%: p = 0.75). (F) Top: pupil centroid positions for all trials following saline injection (blue) and CNO injection (red) from one example imaging session before (left) or after (right) correction for the pupil position and motion, overlaid on the background pupil image (See STAR Methods). Bottom: distributions of the horizontal pupil displacement in trials following saline and CNO injection before (left) or after (right) correction, respectively. The correction resulted in more similar and restricted horizontal pupil displacement after saline and CNO injection. (G) The mean suppression index of each FOV was not significantly altered after pupil position and pupil motion correction in hM4Di-injected mice (p = 0.57). (H) Similar to G but in mCherry-injected mice (p = 0.44). Only dLGN neurons with responses significantly altered after CNO administration both before and after the correction were included here. (I) Distributions of the change in the preferred motion axis (Preferred axis_CNO_ – Preferred axis_Saline_) were similar between hM4Di- and mCherry-injected mice (p = 0.43). (J) Left: Average normalized tuning curves from all horizontal-axis-preferring dLGN neurons following saline and CNO injections (p = 0.14 at -180°, p = 0.41 at -90°, p = 0.003 at 0°, p = 0.18 at 90°; n = 106 neurons from all FOVs). Right: ASI values across individual neurons after CNO injection were significantly smaller than after saline injection (ASI: after saline inj., 0.330 ± 0.014, after CNO inj., 0.291 ± 0.018; p = 0.048). (K) Similar to J but for all vertical-axis-preferring neurons (p = 0.002 at -180°, p = 0.14 at -90°, p = 0.02 at 0°, p = 0.12 at 90°; n = 61 neurons from all FOVs). Right: ASI after saline and after CNO injection was not significantly different (ASI after saline inj., 0.353 ± 0.020; ASI after CNO inj., 0.332 ± 0.023; p = 0.215). (L) Only the number of horizontal-axis-preferring dLGN neurons was significantly reduced after CNO administration in hM4Di-labeled mice (Horizontal Saline: 5.19 ± 0.75; Horizontal CNO: 3.22 ± 0.71; Vertical Saline: 3.37±0.46; Vertical CNO: 3.15±0.41; horizontal p = 0.048, vertical p = 0.64). (M) Results in M-Q were from mCherry-injected mice. Left: Average normalized tuning curves across temporal-direction-preferring dLGN neurons after CNO (orange) and after saline (blue) injection were not significantly different in each of the directions (p = 0.88 at -180°, p = 0.58 at 0°; n = 44 neurons from all FOVs). Right: DSI values after saline and after CNO injection were not significantly different (after saline.: 0.467 ± 0.034, after CNO: 0.438 ± 0.038; p = 0.43). The red cross represents mean ± SEM in M-P. (N) Similar to M but for all nasal-direction-preferring dLGN neurons. Left: n = 49 neurons from all FOVs; p = 0.86 at -180°, p = 0.77 at 0°. Right: Scatterplot of the DSI after saline and CNO injections (after saline: 0.499 ± 0.030, after CNO: 0.450 ± 0.034; p = 0.25). (O) Similar to M but for all horizontal-axis-preferring dLGN neurons (n = 63 neurons; p = 0.47 at - 180°, p = 0.92 at -90°, p = 0.23 at 0°, p = 0.77 at 90°). Right: ASI values after saline and after CNO injection were not significantly different (after saline: 0.300 ± 0.016, after CNO: 0.270 ± 0.015; p = 0.11). (P) Similar to M but for all vertical-axis-preferring dLGN neurons (n = 29 neurons; p = 0.85 at -180°, p = 0.62 at -90°, p = 0.74 at 0°, p = 0.72 at 90°). Right: ASI values after saline and after CNO injection were not significantly different (after saline: 0.385 ± 0.031, after CNO: 0.373 ± 0.038; p = 0.83). (Q) Similar to L but in mCherry-labeled control mice (Horizontal Saline: 5.25 ± 0.95; Horizontal CNO: 4.19 ± 0.75; Vertical Saline: 3.81 ± 0.59; Vertical CNO: 3.75 ± 0.72; horizontal p = 0.37, vertical p = 0.95). All data are presented as mean ± SEM. *, p < 0.05; **, p < 0.01; ***, p < 0.001. A-E, G-Q: linear mixed-effects model.

**Figure S5.**
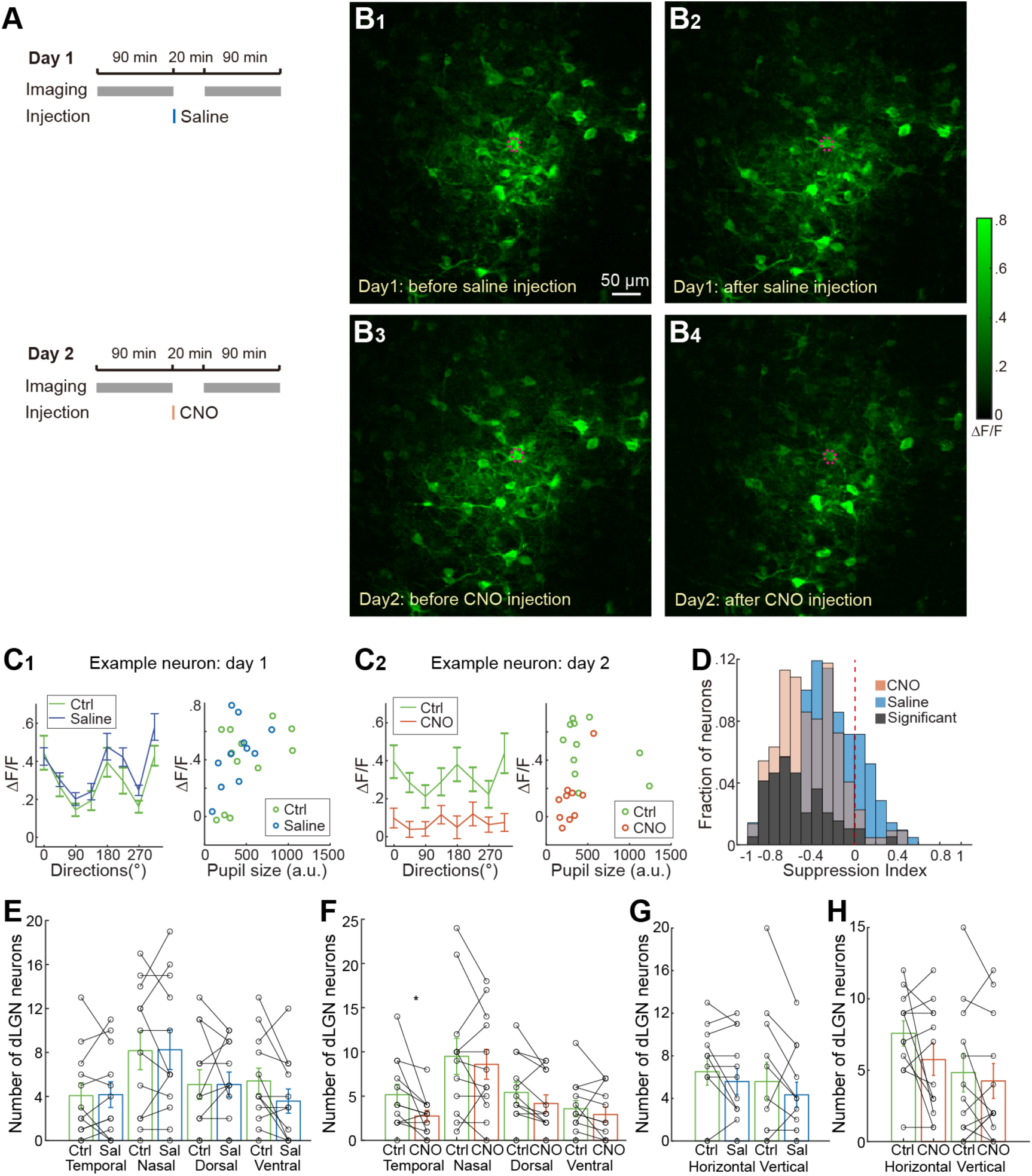
The Majority of dLGN Neurons were Suppressed after Silencing Collicular Input in a Two-day Experimental Paradigm. (A) Experimental timeline. Responses to the same set of 90-minute visual stimuli from the same field of view were recorded during two imaging sessions on consecutive days. On day 1, responses were recorded for 90 minutes before and after saline injection. On day 2, responses were recorded for 90 minutes before and after CNO injection. In this experiment, AAV-FLEX-hM4Di-mCherry was injected into the SC and AAV-GCaMP6f was injected into the dLGN of Rorβ-Cre mice. (B) Visually evoked fluorescence changes (maximum projection across stimulation conditions) in the same example FOV on day 1 before (B_1_) and after (B_2_) saline injection and on day 2 before (B_3_) and after (B_4_) CNO injection. Responses from the circled example dLGN neuron are shown in (C). (C) C_1_: Left, tuning curves of the example neuron before (green) and after (blue) saline injection on day 1; right, similar pupil size distributions before and after saline injection. C_2_: Left, tuning curves of the example neuron before (green) and after (orange) CNO injection on day 2; right, similar pupil size distributions before and after CNO injection. (D) Distributions of suppression indices of dLGN neurons after saline injection on day 1 (blue) and after CNO injection on day 2 (orange). Black, the fraction of neurons with significantly different suppression indices between the two days (61 out of 168 neurons; see STAR Methods). (E) The number of direction-selective neurons was not significantly affected after saline injection in hM4Di-labeled mice (temporal: p = 0.97, nasal: p = 0.97, dorsal: p = 0.93, ventral: p = 0.15, linear mixed-effects model). (F) The number of temporal-direction-preferring direction-selective dLGN neurons was significantly reduced after CNO injection (temporal: p = 0.03, nasal: p = 0.60, dorsal: p = 0.23, ventral: p = 0.39). (G) The number of axis-selective neurons was not significantly affected after saline injection in hM4Di- labeled mice (horizontal: p = 0.53, vertical: p = 0.36). (H) The number of axis-selective neurons before and after CNO injection (horizontal: p = 0.09, vertical: p = 0.49) N = 12 FOVs from 9 mice for (E-H). All the data are presented as mean ± SEM. *, p < 0.05; **, p < 0.01; ***, p < 0.001. E-H: linear mixed-effects model.

**Figure S6.**
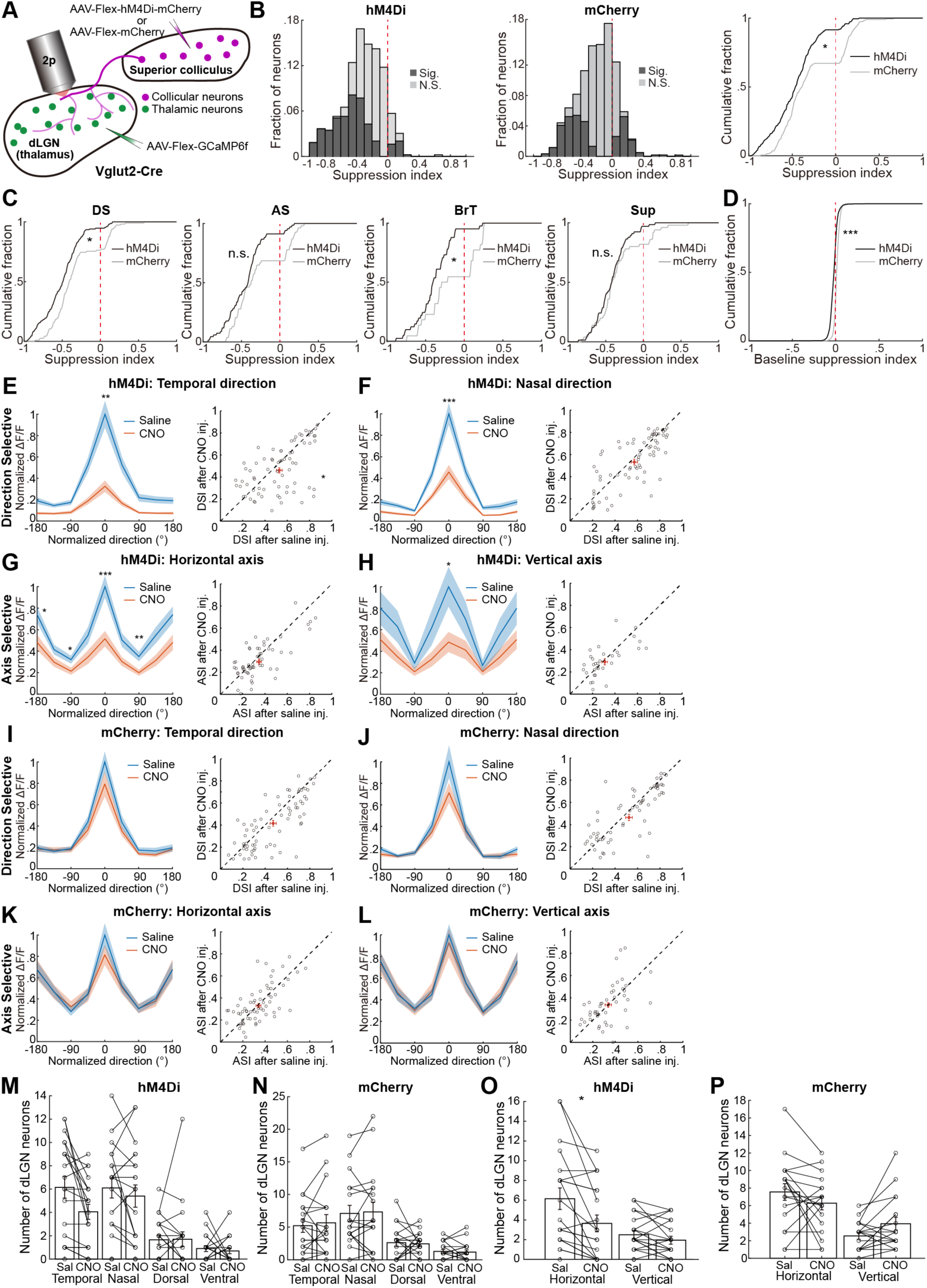
Silencing of Excitatory Collicular Input Reduced Visual Responses in Excitatory dLGN Neurons and the Number of Excitatory dLGN Neurons Preferring the Temporal Direction and Horizontal Axis of Motion. (A) Schematic of measuring the influence of excitatory colliculogeniculate input on excitatory dLGN neurons (thalamocortical cells) in Vglut2-Cre mice. (B) Distribution of visual response suppression indices of excitatory dLGN neurons after administering CNO in the mice injected with Cre-dependent hM4Di (left) or mCherry (middle). Black, significantly affected; gray, not significantly affected. 512 out of 1290 neurons or 429 out of 1686 neurons were significantly affected by CNO injection in hM4Di (n = 20 FOVs from 7 mice) or mCherry-injected mice (n = 18 FOVs from 6 mice), respectively. Right: Cumulative distributions of response suppression indices of excitatory dLGN neurons were significantly different between hM4Di- and mCherry-injected mice (p = 0.041, linear mixed-effects model). (C) Cumulative distributions of response suppression indices for distinct functional categories of excitatory dLGN neurons in hM4Di- or mCherry-injected mice (DS, p = 0.013; AS, p = 0.058; BrT, p = 0.04, Sup, p = 0.28). (D) Cumulative distributions of baseline suppression indices for excitatory dLGN neurons were significantly different between hM4Di- and mCherry-injected mice (p < 10^-3^). (E) Left: normalized tuning curves of all significantly impacted temporal-direction-preferring excitatory dLGN neurons in hM4Di-injected mice after saline (blue) and after CNO (orange) injection (p = 0.09 at -180°, p = 0.004 at 0°; n = 77 neurons). Right: Scatter plot of the DSI after saline and after CNO injection. The mean DSI after saline and after CNO injection was significantly different (DSI after saline injection: 0.529 ± 0.024, after CNO injection: 0.461 ± 0.025; p = 0.04). Red crosses in E-L represent the mean ± SEM. (F) Similar to E but for the significantly impacted nasal-direction-preferring excitatory neurons (n = 70 neurons; left: p = 0.08 at -180°, p < 10^-3^ at 0°; right: DSI after saline injection: 0.571 ± 0.026, after CNO injection: 0.532 ± 0.027, p = 0.25). (G) Similar to E but for the significantly impacted horizontal-axis-preferring excitatory neurons (n = 66 neurons; left: p = 0.011 at -180°, p = 0.02 at -90°, p < 10^-3^ at 0°, p = 0.002 at 90°; right: ASI after saline injection: 0.347 ± 0.022, after CNO injection: 0.295 ± 0.020, p = 0.08). (H) Similar to E but for the significantly impacted vertical-axis-preferring excitatory neurons (n = 34 neurons; left: p = 0.12 at -180°, p = 0.31 at -90°, p = 0.02 at 0°, p = 0.41 at 90°; right: ASI after saline inj.: 0.309 ± 0.027, after CNO inj.: 0.291 ± 0.025, p = 0.56). (I) I-L are in mCherry-injected mice. I: similar to E but for the significantly impacted temporal- direction-preferring excitatory neurons in mCherry-injected mice (n = 69 neurons; including both significantly enhanced and significantly suppressed neurons after CNO injection; left: p = 0.97 at -180°, p = 0.19 at 0°; right: DSI after saline inj.: 0.471 ± 0.027, after CNO inj.: 0.414 ± 0.028, p = 0.11). (J) Similar to F but in mCherry-injected mice (n = 68 neurons; left: p = 0.07 at -180°, p = 0.14 at 0°; right: DSI after saline inj.: 0.523 ± 0.029, after CNO inj.: 0.471 ± 0.030, p = 0.15). (K) Similar to G but in mCherry-injected mice (n = 64 neurons; left: p = 0.68 at -180°, p = 0.50 at -90°, p = 0.27 at 0°, p = 0.92 at 90°; right: ASI after saline inj.: 0.341 ± 0.019, after CNO inj.: 0.330 ± 0.022, p = 0.70). (L) Similar to H but in mCherry-injected mice (n = 46 neurons; left: p = 0.77 at -180°, p = 0.75 at -90°, p = 0.76 at 0°, p = 0.87 at 90°; right: ASI after saline inj.: 0.337 ± 0.022, after CNO inj.: 0.337 ± 0.028, p = 0.89). (M) Only the number of temporal-direction-preferring excitatory dLGN neurons was reduced after CNO injection in hM4Di-injected Vglut2-Cre mice (temporal saline: 6.15 ± 0.92; temporal CNO: 4.05 ± 0.65; nasal saline: 6.10 ± 0.86; nasal CNO: 5.40 ± 0.95; dorsal saline: 1.65 ± 0.39; dorsal CNO: 1.70 ± 0.63; ventral saline: 0.90 ± 0.25; ventral CNO: 0.70 ± 0.24; temporal direction, p = 0.054; nasal direction, p = 0.42; dorsal (upward) direction, p = 0.95; ventral (downward) direction, p = 0.59). (N) The number of direction-selective excitatory dLGN neurons was not significantly altered after CNO injection in mCherry-injected mice (temporal saline: 5.22 ± 1.03; temporal CNO: 5.67 ± 1.26; nasal saline: 7.11 ± 1.23; nasal CNO: 7.33 ±1.48; dorsal saline: 2.61 ± 0.61; dorsal CNO: 2.44 ± 0.40; ventral saline: 1.28 ± 0.30; ventral CNO: 1.17 ± 0.35; temporal direction, p = 0.95; nasal direction, p = 0.91; dorsal direction, p = 0.81; ventral direction, p = 0.62). (O) Only the number of horizontal-axis-preferring excitatory dLGN neurons was significantly reduced after CNO injection in hM4Di-injected mice (horizontal saline: 6.15 ± 1.08; horizontal CNO: 3.65 ± 0.85; vertical saline: 2.50 ± 0.42; vertical CNO: 1.95 ± 0.40; horizontal axis, p = 0.04; vertical axis, p = 0.27). (P) The number of axis-selective excitatory dLGN neurons was not significantly altered after CNO injection in mCherry-injected mice (horizontal saline: 7.56 ± 0.94; horizontal CNO: 6.28 ± 0.76; vertical saline: 2.56 ± 0.39; vertical CNO: 3.94 ± 0.72; horizontal axis, p = 0.23; vertical axis, p = 0.07). All the data are presented as mean ± SEM. *, p < 0.05; **, p < 0.01; ***, p < 0.001. B-P: linear mixed-effects model.

**Figure S7.**
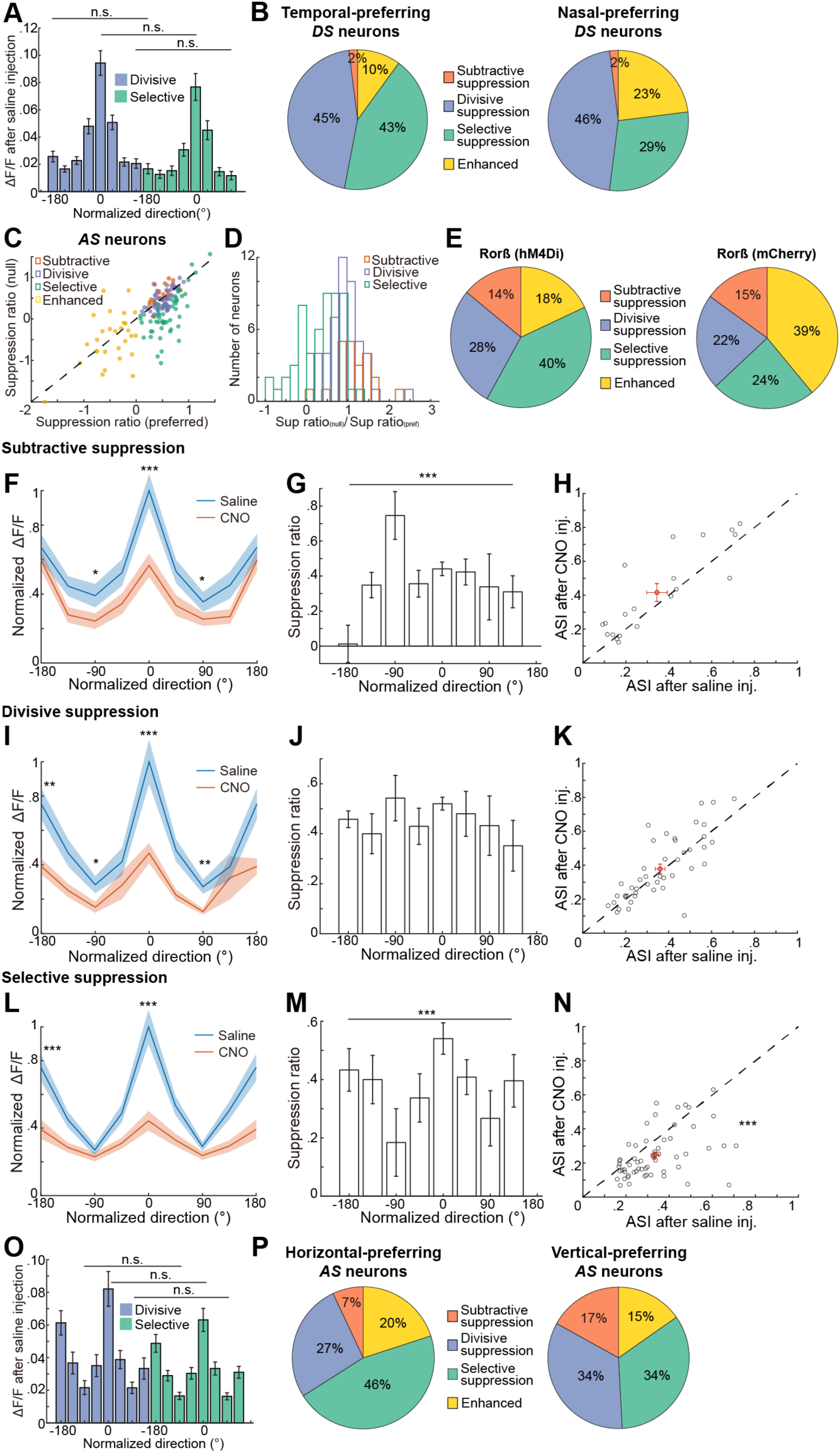
Silencing Colliculogeniculate Input Divisively or Selectively Suppressed Direction-selective and Axis-selective dLGN Neurons. (A) Visually evoked fluorescence changes (ΔF/F) in divisively-suppressed (blue) and selectively-suppressed (green) DS dLGN neurons were not significantly different before CNO injection (between divisively and selectively suppressed neurons: p = 0.10 at -180°, p = 0.22 at 0°, p = 0.06 at 135°, linear mixed-effects model). This result suggests that the relatively small suppression in the null direction and large suppression in the preferred direction in ‘selectively-suppressed’ DS dLGN neurons were not caused by the ‘floor effect’ or saturation of calcium signals. (B) DS dLGN neurons preferring the temporal direction (left) had a higher proportion of selectively-suppressed neurons compared to those preferring the nasal direction (right) (p = 0.049, Chi-squared test; p = 1.00, 0.89, 0.04, and 0.013 for subtractive, divisive, selective, and enhanced categories, respectively, Chi-squared *post hoc* Pearson residuals test). (C) Scatterplot of the suppression ratio for the preferred motion axis vs. the null (orthogonal) axis under subtractive (orange), divisive (blue), or selective (green) suppression, or enhancement of responses following the silencing of collicular input. Results in C, D, F-O are from 27 FOVs of 11 hM4Di-injected Rorβ-Cre mice. (D) Distributions of the ratio of the suppression ratio for the null axis over the suppression ratio for the preferred axis for dLGN neurons under different types of suppression. (E) Larger fractions of AS dLGN neurons underwent divisive and selective suppression in hM4Di-injected Rorβ-Cre mice (n = 23, 62, 52, and 27 neurons for the subtractive, divisive, selective suppression, and enhanced categories, respectively) compared to mCherry-injected Rorβ-Cre mice (n = 13, 24, 18, and 36 neurons for the subtractive, divisive, selective suppression, and enhanced categories, respectively) (p=0.005, Chi-squared test; p = 0.95, 0.06, 0.04, and 4.25*10^-5^ for subtractive, divisive, selective, and enhanced categories, respectively, Chi-squared *post hoc* Pearson residuals test). Note that the proportion of selectively-suppressed AS neurons relative to divisively-suppressed AS neurons was higher in hM4Di-injected mice compared to mCherry-injected mice. (F) Averaged, peak-aligned, normalized tuning curves for AS dLGN neurons exhibiting subtractive suppression, shown after saline and after CNO injection (p = 0.33 at -180°, p = 0.02 at -90°, p = 1.0*10^-5^ at 0°, p = 0.04 at 90°; n = 23 neurons). (G) The suppression ratios were significantly different across normalized directions for neurons in F (p < 0.001, Kruskal–Wallis test; p < 0.0001 between -180° and -90°, p = 0.006 between -180° and 0°, p = 0.01 between -180° and 45°, Dunn’s multiple comparisons test). (H) In subtractively-suppressed AS dLGN neurons, the ASI after CNO injection was larger than after saline injection (ASI after saline inj.: 0.345 ± 0.052; after CNO inj.: 0.416 ± 0.048; p = 0.33). (I) Similar to F but for AS dLGN neurons exhibiting divisive suppression after CNO injection (p = 0.0012 at -180°, p = 0.048 at -90°, p < 10^-3^ at 0°, p = 0.0010 at 90°; n = 52 neurons). (J) The suppression ratios were not significantly different across normalized directions for neurons in I (p = 0.56, Kruskal–Wallis test). (K) In divisively-suppressed AS dLGN neurons, the ASI after CNO injection was not significantly different from that after saline injection (ASI after saline inj.: 0.360 ± 0.022; after CNO inj.: 0.379 ± 0.027; p = 0.54). (L) Similar to F but for AS dLGN neurons exhibiting selective suppression (p < 10^-3^ at -180°, p = 0.20 at -90°, p < 10^-5^ at 0°, p = 0.18 at 90°; n = 62 neurons). (M) A significant difference was observed in the suppression ratio across normalized directions for neurons in L (p < 0.001, Kruskal–Wallis test; p < 0.0001 between -180° and -90°, p < 0.0001 between -180° and 90°, p < 0.0001 between -90° and 0°, p = 0.0259 between -90° and 135°, p < 0.0001 between 0° and 90°, Dunn’s multiple comparisons test). (N) In selectively-suppressed AS dLGN neurons, the ASI after CNO injection was significantly smaller than after saline injection (ASI after saline inj.: 0.334 ± 0.018, after CNO inj.: 0.247 ± 0.017, p <10^-3^). (O) Visually evoked fluorescence changes (ΔF/F) for the preferred and null axes were not significantly different between divisively-suppressed (blue) and selectively-suppressed (green) AS dLGN neurons (p = 0.19 at -180°, p = 0.42 at -90°, p = 0.25 at 0°, p = 0.35 at 90°). This result suggests that the relatively smaller suppression in the null axis and relatively larger suppression in the preferred axis in ‘selectively-suppressed’ AS dLGN neurons were not due to the ‘floor effect’ or saturation of calcium signals. (P) Similar to B but in AS dLGN neurons preferring the horizontal motion axis (left) and vertical motion axis (right) (p = 0.04, Chi-squared test; p = 0.03, 0.26, 0.06, and 0.37 for subtractive, divisive, selective, and enhanced categories, respectively, Chi-squared *post hoc* Pearson residuals test) All the data are presented as mean ± SEM. *, p < 0.05; **, p < 0.01; ***, p < 0.001. A, F, H, I, K, L, N, and O: linear mixed-effects model.

**Figure S8.**
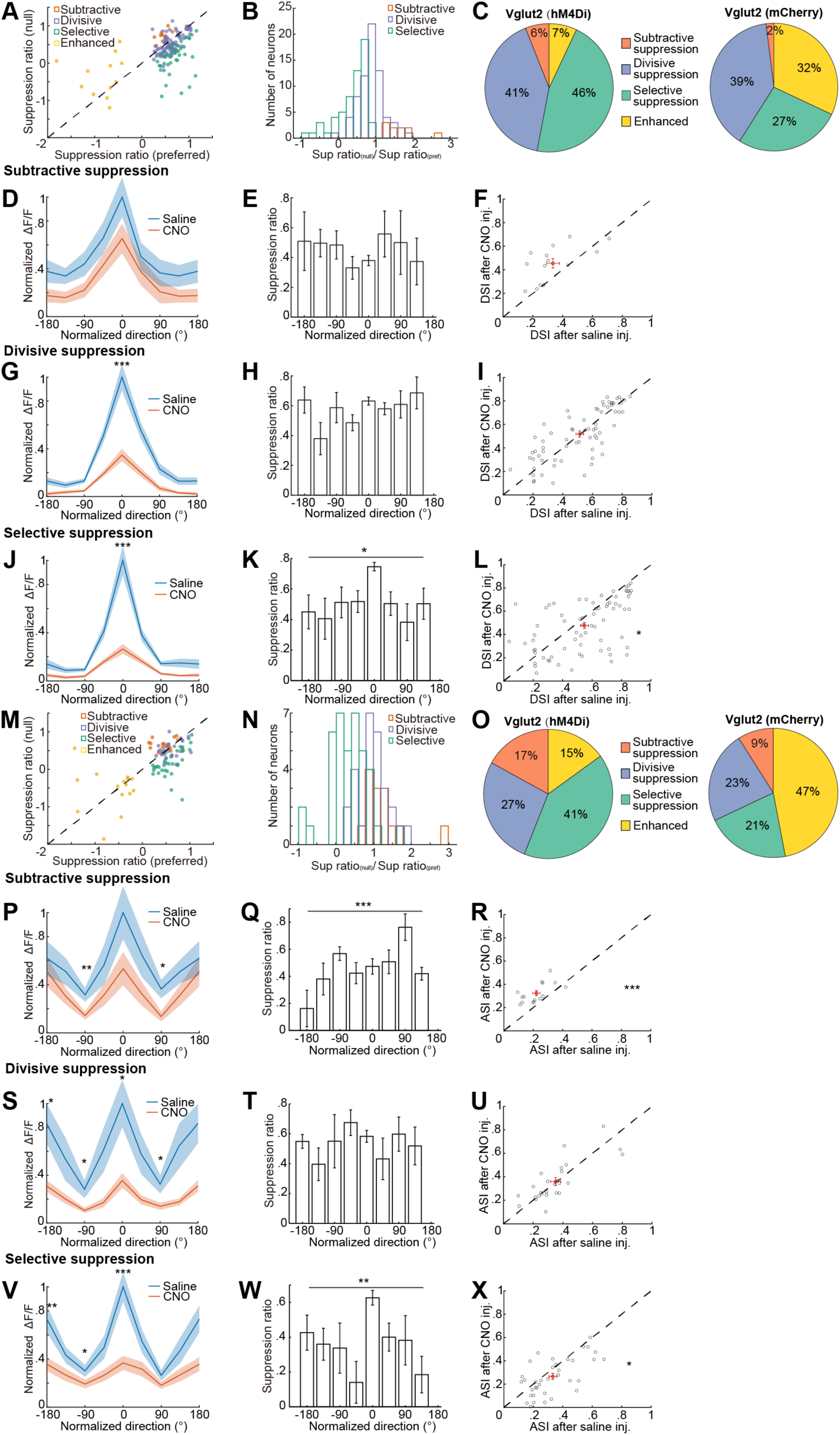
Distinct Suppression Effects on Excitatory dLGN Neurons. (A) Scatterplot of the suppression ratio in the null direction vs. in the preferred direction for DS excitatory dLGN neurons under subtractive (orange), divisive (blue), and selective (green) suppression, and response enhancement (yellow) following the silencing of colliculogeniculate input. Results in A-X are from 20 FOVs of 7 Cre-dependent-hM4Di-injected Vglut2-Cre mice. (B) Distinct distributions of the suppression ratio in the null direction/suppression ratio in the preferred direction for DS excitatory dLGN neurons under subtractive suppression (orange), divisive suppression (blue), and selective suppression (green). (C) Larger fractions of DS excitatory dLGN neurons underwent divisive and selective suppression in hM4Di-injected Vglut2 mice (n = 10, 78, 70, and 11 neurons for subtractive, divisive, selective suppression, and enhanced categories, respectively) compared to mCherry-injected Vglut2 mice (n = 4, 62, 44, and 51 neurons for the subtractive, divisive, selective suppression, and enhanced categories, respectively). p < 0.001, Chi-squared test; p = 0.12, 0.62, 0.0004, and 1.4*10^-7^ for subtractive, divisive, selective suppression, and enhanced categories, respectively, *post hoc* Pearson residuals test. (D) Averaged, peak-aligned, normalized tuning curves after saline injection and after CNO injection for DS excitatory dLGN neurons that exhibited subtractive suppression following CNO injection (p = 0.19 at -180°, p = 0.22 at 0°; n = 10 neurons). (E) The suppression ratio for neurons in D was not significantly different across normalized directions (p = 0.47; Kruskal–Wallis test). (F) In subtractively-suppressed DS excitatory dLGN neurons, the DSI after CNO injection was larger than that after saline injection (DSI after saline injection: 0.395 ± 0.061, after CNO injection: 0.478 ± 0.072; p = 0.16). (G) Similar to D but for DS excitatory dLGN neurons exhibiting divisive suppression after CNO injection (p = 0.06 at -180°, p = 4.5*10^-5^ at 0°; n = 70 neurons). (H) The suppression ratio for neurons in G was not significantly different across normalized directions (p = 0.08; Kruskal–Wallis test). (I) The DSIs after CNO injection and after saline injection were not significantly different in divisively-suppressed DS excitatory dLGN neurons (DSI after saline inj.: 0.514 ± 0.025, after CNO inj.: 0.518 ± 0.026; p = 0.89). (J) Similar as D but for DS excitatory dLGN neurons exhibiting selective suppression in the preferred direction after CNO injection (p = 0.07 at -180°, p = 1.9*10^-4^ at 0°; n = 78 neurons). (K) The suppression ratios were significantly different across normalized directions (p = 0.016, Kruskal–Wallis test). (L) In selectively-suppressed DS excitatory dLGN neurons, the DSI after CNO injection was significantly smaller than that after saline injection (DSI after saline inj.: 0.548 ± 0.027, after CNO inj.: 0.475 ± 0.025; p = 0.04). (M) Scatterplot of the suppression ratio for the null motion axis vs. the suppression ratio for the preferred axis for AS excitatory dLGN neurons exhibiting subtractive (orange), divisive (blue), and selective (green) suppression, and enhanced responses (yellow) following the silencing of excitatory collicular neurons. (N) Distinct distributions of the suppression ratio for the null axis/suppression ratio for the preferred axis for AS excitatory dLGN neurons under different types of suppression. (O) Larger fractions of AS excitatory dLGN neurons underwent divisive and selective suppression in hM4Di-injected Vglut2 mice (n = 17, 27, 41, and 15 neurons for the subtractively, divisively, selectively suppressed, and enhanced categories, respectively) compared to mCherry-injected Vglut2 mice (n = 10, 24, 23, and 51 neurons for the subtractively, divisively, selectively suppressed, and enhanced categories, respectively). p < 0.001, Chi-squared test; p = 0.08, 0.64, 0.002, and 2.5*10^-5^ for subtractively, divisively, selectively suppressed, and enhanced categories, respectively, Chi-squared *post hoc* Pearson residuals test). The proportion of selectively-suppressed AS neurons relative to divisively-suppressed AS neurons was higher in hM4Di-injected mice compared to mCherry-injected mice. (P) Averaged, peak-aligned, normalized tuning curves after saline injection and after CNO injection for AS excitatory dLGN neurons that exhibited subtractive suppression following CNO injection (p = 0.55 at -180°, p = 0.008 at -90°, p = 0.09 at 0°, p = 0.02 at 90°; n = 17 neurons). (Q) The suppression ratio for neurons in P was significantly different across normalized directions (p = 0.0007, Kruskal–Wallis test; p = 0.0002 between -180° and 90°; Dunn’s multiple comparisons test). (R) In subtractively-suppressed AS excitatory dLGN neurons, the ASI after CNO injection was significantly larger than after saline injection (ASI after saline inj.: 0.221 ± 0.022; after CNO inj.: 0.324 ± 0.024; p < 10^-3^). (S) Similar to P but for AS excitatory dLGN neurons exhibiting divisive suppression after CNO injection (p = 0.02 at -180°, p = 0.04 at -90°, p = 0.010 at 0°, p = 0.03 at 90°; n = 27 neurons). (T) The suppression ratio for neurons in S was not significantly different across normalized directions (p = 0.49, Kruskal–Wallis test). (U) The ASIs after CNO injection and after saline injection were not significantly different in divisively-suppressed AS excitatory dLGN neurons (ASI after saline inj.: 0.352 ± 0.033, after CNO inj.: 0.360 ± 0.033; p = 0.87). (V) Similar to P but for the AS excitatory dLGN neurons exhibiting selective suppression in the preferred direction after CNO injection (p = 0.002 at -180°, p = 0.03 at -90°, p < 10^-4^ at 0°, p = 0.09 at 90°; n = 41 neurons). (W) The suppression ratio for neurons in V was significantly different across normalized directions (p = 0.005, Kruskal–Wallis test; p = 0.007 between 0° and -45°, p = 0.007 between 0° and 135°, Dunn’s multiple comparisons test). (X) In selectively-suppressed AS excitatory dLGN neurons, the ASI after CNO injection was significantly smaller than after saline injection (ASI after saline inj.: 0.333 ± 0.025, after CNO inj.: 0.266 ± 0.025; p = 0.03). All the data are plotted as mean ± SEM. *, p < 0.05; **, p < 0.01; ***, p < 0.001. D, F, G, I, J, L, P, R, S, U, V and X: linear mixed-effects model.

**Figure S9.**
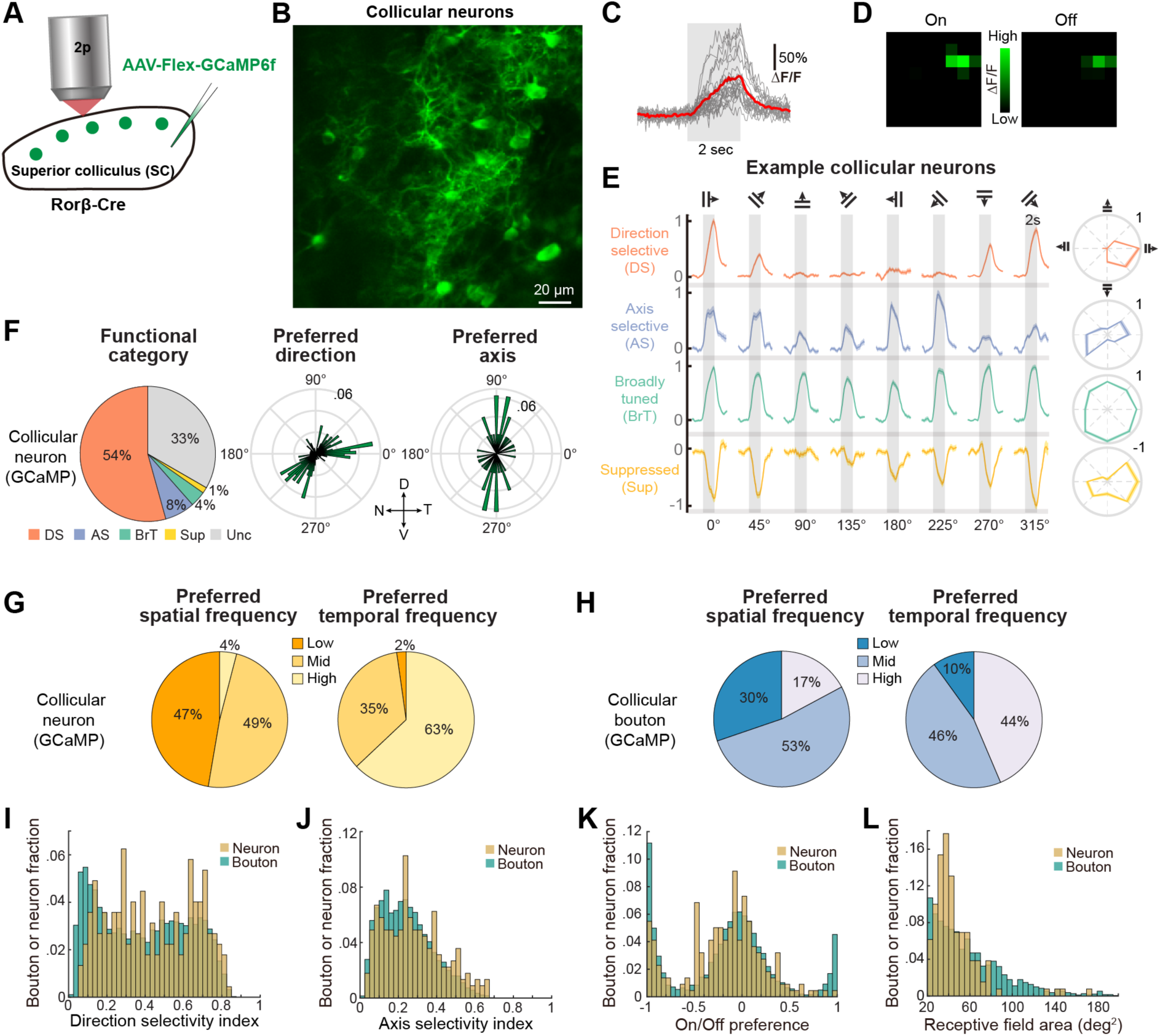
Rorβ-positive Collicular Neurons Showed Similar Visual Response Properties to Collicular Boutons in the dLGN. (A) Schematic of two-photon calcium imaging of Rorβ-positive collicular neurons in the SC of awake, head-restrained Rorβ-Cre mice. (B) Averaged raw images of collicular neurons expressing GCaMP6f in an example field of view. (C) Single-trial responses of a collicular neuron to the repeated presentation of the same visual stimuli (gray bar). (D) The On and Off receptive fields for a collicular neuron mapped by sparse flashing square stimuli. (E) Collicular neurons were classified into functional categories based on their responses to 2-s presentations of drifting gratings (gray bars). Left: Example normalized response time courses of one collicular neuron in each category. Right: Normalized mean response tuning curves. (F) Left: Distribution of neurons across functional categories. Middle: Polar distribution of the preferred motion direction for direction-selective neurons. Right: Polar distribution of the preferred motion axis for axis-selective neurons (n = 188 DS neurons, 24 AS neurons, 12 BrT neurons, 4 Sup neurons; comparisons between the distributions of GCaMP-labeled collicular neurons and collicular boutons in Figure 1F: p < 0.0001, Chi-squared test; DS: p = 0.36, AS: p = 0.01, BrT p < 10^-6^, Sup p = 0.01, Unclassified (Unc) p < 10^-13^, Chi-squared *post hoc* Pearson residuals test). (G) Distributions of collicular neurons across spatial frequency (SF) preferences (Low, Mid, High: SF = 0.02, 0.08, 0.32 cpd, respectively), and across temporal frequency (TF) preferences (Low, Mid, High, TF = 0.5, 2, 8 Hz, respectively). (H) Similar to G but in Rorb-Cre+ colliculogeniculate boutons labeled with GCaMP (between G and H, SF: p < 0.0001, Chi-squared test; Low p < 10^-66^, Middle p = 0.43, High p < 10^-30^, *post hoc* Pearson residuals test; TF: p < 0.0001, Chi-squared test; Low p < 10^-17^, Middle p < 10^-5^, High p < 10^-66^, *post hoc* Pearson residuals test). (I) Distributions of the DSI across GCaMP6f-labeled collicular neurons (yellow) and collicular boutons (blue) (mean of DSI: collicular neurons 0.438, collicular boutons 0.386, p = 0.04). G-K: n = 224 neurons, 8 FOVs, 2 mice; n = 9292 boutons, 10 FOVs, 5 mice. (J) Distributions of the ASI across GCaMP6f-labeled collicular neurons (yellow) and collicular boutons (blue) (mean of ASI: collicular neurons 0.281, collicular boutons 0.243, p = 0.03). (K) Distributions of the On/Off preference across collicular neurons and across collicular boutons (the means of the two distributions were not significantly different, p = 0.49). (L) Distributions of the receptive field areas across collicular neurons and across collicular boutons (the means of the two distributions were not significantly different, p = 0.59). All the data was plotted as mean ± SEM. *, p < 0.05; **, p < 0.01; ***, p < 0.001. I-M: linear mixed-effects model.

## REFERENCES

1. Guillery, R.W., and Sherman, S.M. (2002). Thalamic relay functions and their role in corticocortical communication: Generalizations from the visual system. Neuron 33, 163–175. 10.1016/S0896-6273(01)00582-7.

2. Guido, W. (2018). Development, form, and function of the mouse visual thalamus. Journal of Neurophysiology 120, 211–225. 10.1152/jn.00651.2017.

3. Güillery, R.W. (1969). A quantitative study of synaptic interconnections in the dorsal lateral geniculate nucleus of the cat. Zeitschrift für Zellforschung und Mikroskopische Anatomie 96, 39–48. 10.1007/BF00321475.

4. Boyapati, J., and Henry, G. (1984). Corticofugal axons in the lateral geniculate nucleus of the cat. Experimental brain research 53, 335–340. 10.1007/BF00238163.

5. Smith, Y., Paré, D., Deschênes, M., Parent, A., and Steriade, M. (1988). Cholinergic and non-cholinergic projections from the upper brainstem core to the visual thalamus in the cat. Experimental Brain Research 70, 166–180. 10.1007/BF00271858.

6. Harting, J.K., Huerta, M.F., Hashikawa, T., and van Lieshout, D.P. (1991). Projection of the mammalian superior colliculus upon the dorsal lateral geniculate nucleus: Organization of tectogeniculate pathways in nineteen species. Journal of Comparative Neurology 304, 275–306. 10.1002/cne.903040210.

7. Wilson, J.R. (1993). Circuitry of the dorsal lateral geniculate nucleus in the cat and monkey. Acta Anatomica 147, 1–13. 10.1159/000147475.

8. Feig, S., and Harting, J.K. (1994). Ultrastuctural studies of the primate lateral geniculate nucleus: Morphology and spatial relationships of axon terminals arising from the retina, visual cortex (area 17), superior colliculus, parabigminal nucleus, and pretectum of Galago crassicaudatus. Journal of Comparative Neurology 343, 17–34. 10.1002/cne.903430103.

9. McCormick, D.A., and Bal, T. (1997). Sleep and arousal: Thalamocortical mechanisms. Annual Review of Neuroscience 20, 185–215. 10.1146/annurev.neuro.20.1.185.

10. Erişir, A., Van Horn, S.C., and Sherman, S.M. (1997). Relative numbers of cortical and brainstem inputs to the lateral geniculate nucleus. Proceedings of the National Academy of Sciences of the United States of America 94, 1517–1520. 10.1073/pnas.94.4.1517.

11. Gale, S.D., and Murphy, G.J. (2014). Distinct representation and distribution of visual information by specific cell types in mouse superficial superior colliculus. Journal of Neuroscience 34, 13458–13471. 10.1523/JNEUROSCI.2768-14.2014.

12. Basso, M.A., Bickford, M.E., and Cang, J. (2021). Unraveling circuits of visual perception and cognition through the superior colliculus. Neuron 109, 918–937. 10.1016/j.neuron.2021.01.013.

13. Reggiani, J.D.S., Jiang, Q., Barbini, M., Lutas, A., Liang, L., Fernando, J., Deng, F., Wan, J., Li, Y., Chen, C., and Andermann, M.L. (2023). Brainstem serotonin neurons selectively gate retinal information flow to thalamus. Neuron 111, 711–726 e711. 10.1016/j.neuron.2022.12.006.

14. Van Horn, S.C., Erişir, A., and Sherman, S.M. (2000). Relative distribution of synapses in the A-laminae of the lateral geniculate nucleus of the cat. Journal of Comparative Neurology 416, 509–520. 10.1002/(SICI)1096-9861(20000124)416:4<509::AID-CNE7>3.0.CO;2-H.

15. Mooney, R.D., Nikoletseas, M.M., Ruiz, S.A., and Rhoades, R.W. (1988). Receptive-field properties and morphological characteristics of the superior collicular neurons that project to the lateral posterior and dorsal lateral geniculate nuclei in the hamster. Journal of Neurophysiology 59, 1333–1351. 10.1152/jn.1988.59.5.1333.

16. Torrealba, F., Partlow, G.D., and Guillery, R.W. (1981). Organization of the projection from the superior colliculus to the dorsal lateral geniculate nucleus of the cat. Neuroscience 6, 1341–1360. 10.1016/0306-4522(81)90192-5.

17. Baldwin, M.K.L., and Bourne, J.A. (2017). 3.10 - The Evolution of Subcortical Pathways to the Extrastriate Cortex. Evolution of Nervous Systems (Second Edition), 165-185. 10.1016/B978-0-12-804042-3.00081-6.

18. Bickford, M.E., Zhou, N., Krahe, T.E., Govindaiah, G., and Guido, W. (2015). Retinal and tectal “Driver-Like” inputs converge in the shell of the mouse dorsal lateral geniculate nucleus. Journal of Neuroscience 35, 10523–10534. 10.1523/JNEUROSCI.3375-14.2015.

19. Sahibzada, N., Dean, P., and Redgrave, P. (1986). Movements resembling orientation or avoidance elicited by electrical stimulation of the superior colliculus in rats. Journal of Neuroscience 6, 723–733. 10.1523/jneurosci.06-03-00723.1986.

20. Yilmaz, M., and Meister, M. (2013). Rapid innate defensive responses of mice to looming visual stimuli. Current Biology 23, 2011–2015. 10.1016/j.cub.2013.08.015.

21. Zhao, X., Liu, M., and Cang, J. (2014). Visual cortex modulates the magnitude but not the selectivity of looming-evoked responses in the superior colliculus of awake mice. Neuron 84, 202–213. 10.1016/j.neuron.2014.08.037.

22. Wei, P., Liu, N., Zhang, Z., Liu, X., Tang, Y., He, X., Wu, B., Zhou, Z., Liu, Y., Li, J., et al. (2015). Processing of visually evoked innate fear by a non-canonical thalamic pathway. Nature Communications 6. 10.1038/ncomms7756.

23. Shang, C., Liu, Z., Chen, Z., Shi, Y., Wang, Q., Liu, S., Li, D., and Cao, P. (2015). A parvalbumin-positive excitatory visual pathway to trigger fear responses in mice. Science 348, 1472–1477. 10.1126/science.aaa8694.

24. Ellis, E.M., Gauvain, G., Sivyer, B., and Murphy, G.J. (2016). Shared and distinct retinal input to the mouse superior colliculus and dorsal lateral geniculate nucleus. Journal of Neurophysiology 116, 602–610. 10.1152/jn.00227.2016.

25. Seabrook, T.A., Burbridge, T.J., Crair, M.C., and Huberman, A.D. (2017). Architecture, Function, and Assembly of the Mouse Visual System. Annual Review of Neuroscience 40, 499–538. 10.1146/annurev-neuro-071714-033842.

26. Ito, S., and Feldheim, D.A. (2018). The mouse superior colliculus: An emerging model for studying circuit formation and function. Frontiers in Neural Circuits 12, 1–11. 10.3389/fncir.2018.00010.

27. Cang, J., Savier, E., Barchini, J., and Liu, X. (2018). Visual Function, Organization, and Development of the Mouse Superior Colliculus. Annu Rev Vis Sci 4, 239–262. 10.1146/annurev-vision-091517-034142.

28. Evans, D.A., Stempel, A.V., Vale, R., Ruehle, S., Lefler, Y., and Branco, T. (2018). A synaptic threshold mechanism for computing escape decisions. Nature 558, 590–594. 10.1038/s41586-018-0244-6.

29. Hoy, J.L., Bishop, H.I., and Niell, C.M. (2019). Defined Cell Types in Superior Colliculus Make Distinct Contributions to Prey Capture Behavior in the Mouse. Current Biology 29, 4130–4138.e4135. 10.1016/j.cub.2019.10.017.

30. Fratzl, A., Koltchev, A.M., Vissers, N., Tan, Y.L., Marques-Smith, A., Stempel, A.V., Branco, T., and Hofer, S.B. (2021). Flexible inhibitory control of visually evoked defensive behavior by the ventral lateral geniculate nucleus. Neuron 109, 3810–3822.e3819. 10.1016/j.neuron.2021.09.003.

31. Li, C., Kühn, N.K., Alkislar, I., Sans-Dublanc, A., Zemmouri, F., Paesmans, S., Calzoni, A., Ooms, F., Reinhard, K., and Farrow, K. (2023). Pathway-specific inputs to the superior colliculus support flexible responses to visual threat. Science Advances 9. 10.1126/sciadv.ade3874.

32. Hafed, Z.M., Hoffmann, K.P., Chen, C.Y., and Bogadhi, A.R. (2023). Visual Functions of the Primate Superior Colliculus. Annual Review of Vision Science 9, 361–383. 10.1146/annurev-vision-111022-123817.

33. Zahler, S.H., Taylor, D.E., Wright, B.S., Wong, J.Y., Shvareva, V.A., Park, Y.A., and Feinberg, E.H. (2023). Hindbrain modules differentially transform activity of single collicular neurons to coordinate movements. Cell 186, 3062–3078.e3020. 10.1016/j.cell.2023.05.031.

34. Kerschensteiner, D., and Feller, M.B. (2024). Mapping the Retina onto the Brain. Cold Spring Harbor Perspectives in Biology 16. 10.1101/cshperspect.a041512.

35. Lyon, D.C., Nassi, J.J., and Callaway, E.M. (2010). A Disynaptic Relay from Superior Colliculus to Dorsal Stream Visual Cortex in Macaque Monkey. Neuron 65, 270–279. 10.1016/j.neuron.2010.01.003.

36. Leopold, D.A. (2012). Primary visual cortex: Awareness and blindsight. Annual Review of Neuroscience 35, 91–109. 10.1146/annurev-neuro-062111-150356.

37. Beltramo, R., and Scanziani, M. (2019). A collicular visual cortex: Neocortical space for an ancient midbrain visual structure. Science 363, 64–69. 10.1126/science.aau7052.

38. Bennett, C., Gale, S.D., Garrett, M.E., Newton, M.L., Callaway, E.M., Murphy, G.J., and Olsen, S.R. (2019). Higher-Order Thalamic Circuits Channel Parallel Streams of Visual Information in Mice. Neuron 102, 477–492.e475. 10.1016/j.neuron.2019.02.010.

39. Brenner, J.M., Beltramo, R., Gerfen, C.R., Ruediger, S., and Scanziani, M. (2023). A genetically defined tecto-thalamic pathway drives a system of superior-colliculus-dependent visual cortices. Neuron 111, 2247–2257.e2247. 10.1016/j.neuron.2023.04.022.

40. Marshel, J.H., Kaye, A.P., Nauhaus, I., and Callaway, E.M. (2012). Anterior-Posterior Direction Opponency in the Superficial Mouse Lateral Geniculate Nucleus. Neuron 76, 713–720. 10.1016/j.neuron.2012.09.021.

41. Zhao, X., Chen, H., Liu, X., and Cang, J. (2013). Orientation-selective responses in the mouse lateral geniculate nucleus. Journal of Neuroscience 33, 12751–12763. 10.1523/JNEUROSCI.0095-13.2013.

42. Piscopo, D.M., El-Danaf, R.N., Huberman, A.D., and Niell, C.M. (2013). Diverse visual features encoded in mouse lateral geniculate nucleus. Journal of Neuroscience 33, 4642–4656. 10.1523/JNEUROSCI.5187-12.2013.

43. Scholl, B., Tan, A.Y., Corey, J., and Priebe, N.J. (2013). Emergence of orientation selectivity in the Mammalian visual pathway. J Neurosci 33, 10616–10624. 10.1523/JNEUROSCI.0404-13.2013.

44. Born, G., Schneider-Soupiadis, F.A., Erisken, S., Vaiceliunaite, A., Lao, C.L., Mobarhan, M.H., Spacek, M.A., Einevoll, G.T., and Busse, L. (2021). Corticothalamic feedback sculpts visual spatial integration in mouse thalamus. Nat Neurosci 24, 1711–1720. 10.1038/s41593-021-00943-0.

45. Kaplan, E., and Shapley, R. (1984). The origin of the S (slow) potential in the mammalian Lateral Geniculate Nucleus. Experimental Brain Research 55, 111–116. 10.1007/BF00240504.

46. Cleland, B.G., Dubin, M.W., and Levick, W.R. (1971). Simultaneous recording of input and output of lateral geniculate neurones. Nature New Biology 231, 191–192. 10.1038/newbio231191a0.

47. Cruz-Martin, A., El-Danaf, R.N., Osakada, F., Sriram, B., Dhande, O.S., Nguyen, P.L., Callaway, E.M., Ghosh, A., and Huberman, A.D. (2014). A dedicated circuit links direction-selective retinal ganglion cells to the primary visual cortex. Nature 507, 358–361. 10.1038/nature12989.

48. Wang, L., Sarnaik, R., Rangarajan, K., Liu, X., and Cang, J. (2010). Visual receptive field properties of neurons in the superficial superior colliculus of the mouse. Journal of Neuroscience 30, 16573–16584. 10.1523/JNEUROSCI.3305-10.2010.

49. Inayat, S., Barchini, J., Chen, H., Feng, L., Liu, X., and Cang, J. (2015). Neurons in the most superficial lamina of the mouse superior colliculus are highly selective for stimulus direction. Journal of Neuroscience 35, 7992–8003. 10.1523/JNEUROSCI.0173-15.2015.

50. De Franceschi, G., and Solomon, S.G. (2018). Visual response properties of neurons in the superficial layers of the superior colliculus of awake mouse. Journal of Physiology 596, 6307–6332. 10.1113/JP276964.

51. de Malmazet, D., Kuhn, N.K., and Farrow, K. (2018). Retinotopic Separation of Nasal and Temporal Motion Selectivity in the Mouse Superior Colliculus. Curr Biol 28, 2961–2969 e2964. 10.1016/j.cub.2018.07.001.

52. Chen, H., Savier, E.L., DePiero, V.J., and Cang, J. (2021). Lack of Evidence for Stereotypical Direction Columns in the Mouse Superior Colliculus. J Neurosci 41, 461–473. 10.1523/JNEUROSCI.1155-20.2020.

53. Kasai, M., and Isa, T. (2022). Effects of Light Isoflurane Anesthesia on Organization of Direction and Orientation Selectivity in the Superficial Layer of the Mouse Superior Colliculus. J Neurosci 42, 619–630. 10.1523/JNEUROSCI.1196-21.2021.

54. Gale, S.D., and Murphy, G.J. (2018). Distinct cell types in the superficial superior colliculus project to the dorsal lateral geniculate and lateral posterior thalamic nuclei. Journal of Neurophysiology 120, 1286–1292. 10.1152/jn.00248.2018.

55. Byun, H., Lee, H.L., Liu, H., Forrest, D., Rudenko, A., and Kim, I.J. (2019). Rorβ regulates selective axon-target innervation in the mammalian midbrain. Development (Cambridge) 146. 10.1242/dev.171926.

56. Balcioglu, A., Gillani, R., Doron, M., Burnell, K., Ku, T., Erisir, A., Chung, K., Segev, I., and Nedivi, E. (2023). Mapping thalamic innervation to individual L2/3 pyramidal neurons and modeling their ‘readout’ of visual input. Nature Neuroscience 26, 470–480. 10.1038/s41593-022-01253-9.

57. Hammer, S., Monavarfeshani, A., Lemon, T., Su, J., and Fox, M.A. (2015). Multiple Retinal Axons Converge onto Relay Cells in the Adult Mouse Thalamus. Cell Reports 12, 1575–1583. 10.1016/j.celrep.2015.08.003.

58. Morgan, J.L., Berger, D.R., Wetzel, A.W., and Lichtman, J.W. (2016). The Fuzzy Logic of Network Connectivity in Mouse Visual Thalamus. Cell 165, 192–206. 10.1016/j.cell.2016.02.033.

59. Liang, L., Fratzl, A., Goldey, G., Ramesh, R.N., Sugden, A.U., Morgan, J.L., Chen, C., and Andermann, M.L. (2018). A Fine-Scale Functional Logic to Convergence from Retina to Thalamus. Cell 173, 1343–1355.e1324. 10.1016/j.cell.2018.04.041.

60. Ahmadlou, M., Zweifel, L.S., and Heimel, J.A. (2018). Functional modulation of primary visual cortex by the superior colliculus in the mouse. Nature Communications 9, 1–13. 10.1038/s41467-018-06389-6.

61. Chen, T.W., Wardill, T.J., Sun, Y., Pulver, S.R., Renninger, S.L., Baohan, A., Schreiter, E.R., Kerr, R.A., Orger, M.B., Jayaraman, V., et al. (2013). Ultrasensitive fluorescent proteins for imaging neuronal activity. Nature 499, 295–300. 10.1038/nature12354.

62. Dana, H., Mohar, B., Sun, Y., Narayan, S., Gordus, A., Hasseman, J.P., Tsegaye, G., Holt, G.T., Hu, A., Walpita, D., et al. (2016). Sensitive red protein calcium indicators for imaging neural activity. Elife 5. 10.7554/eLife.12727.

63. Kara, P., Reinagel, P., and Reid, R.C. (2000). Low response variability in simultaneously recorded retinal, thalamic, and cortical neurons. Neuron 27, 635–646. 10.1016/S0896-6273(00)00072-6.

64. Savier, E.L., Chen, H., and Cang, J. (2019). Effects of locomotion on visual responses in the mouse superior colliculus. Journal of Neuroscience 39, 9360–9368. 10.1523/JNEUROSCI.1854-19.2019.

65. Molotchnikoff, S., Delaunais, D., Casanova, C., and Lachapelle, P. (1986). Influence of a local inactivation in the superior colliculus on lateral geniculate responses in rabbits. Brain Research 375, 66–72. 10.1016/0006-8993(86)90959-5.

66. Molotchnikoff, S., Casanova, C., and Cérat, A. (1988). The consequences of the superior colliculus output on lateral geniculate and pulvinar responses. Progress in Brain Research 75, 67–74. 10.1016/S0079-6123(08)60466-5.

67. Xue, J.T., Kim, C.B.Y., Moore, R.J., and Spear, P.D. (1994). Influence of the superior colliculus on responses of lateral geniculate neurons in the cat. Visual Neuroscience 11, 1059–1076. 10.1017/S095252380000688X.

68. Roth, B.L. (2016). DREADDs for Neuroscientists. Neuron 89, 683–694. 10.1016/j.neuron.2016.01.040.

69. Arandia-Romero, I., Tanabe, S., Drugowitsch, J., Kohn, A., and Moreno-Bote, R. (2016). Multiplicative and Additive Modulation of Neuronal Tuning with Population Activity Affects Encoded Information. Neuron 89, 1305–1316. 10.1016/j.neuron.2016.01.044.

70. Lee, S.H., Kwan, A.C., Zhang, S., Phoumthipphavong, V., Flannery, J.G., Masmanidis, S.C., Taniguchi, H., Huang, Z.J., Zhang, F., Boyden, E.S., et al. (2012). Activation of specific interneurons improves V1 feature selectivity and visual perception. Nature 488, 379–383. 10.1038/nature11312.

71. El-Boustani, S., Wilson, N.R., Runyan, C.A., and Sur, M. (2014). El-boustani et al. reply. Nature 508, 2012–2015. 10.1038/nature13130.

72. Ito, S., Feldheim, D.A., and Litke, A.M. (2017). Segregation of Visual Response Properties in the Mouse Superior Colliculus and Their Modulation during Locomotion. J Neurosci 37, 8428–8443. 10.1523/JNEUROSCI.3689-16.2017.

73. Liu, Y., Savier, E.L., DePiero, V.J., Chen, C., Schwalbe, D.C., Abraham-Fan, R.J., Chen, H., Campbell, J.N., and Cang, J. (2023). Mapping visual functions onto molecular cell types in the mouse superior colliculus. Neuron 111, 1876–1886.e1875. 10.1016/j.neuron.2023.03.036.

74. Li, Y.T., and Meister, M. (2023). Functional cell types in the mouse superior colliculus. eLife 12, 1–23. 10.7554/eLife.82367.

75. Schwartz, O.S., and Yonehara, K. (2023). The superior colliculus response space has globally high– and locally low-dimensionality. bioRxiv 4, 2023.2011.2006.565916. doi.org/10.1101/2023.11.06.565916.

76. Whyland, K.L., Slusarczyk, A.S., and Bickford, M.E. (2020). GABAergic cell types in the superficial layers of the mouse superior colliculus. Journal of Comparative Neurology 528, 308–320. 10.1002/cne.24754.

77. de Malmazet, D., Kühn, N.K., Li, C., and Farrow, K. (2024). Retinal origin of orientation but not direction selective maps in the superior colliculus. Current Biology. 10.1016/j.cub.2024.02.001.

78. Shi, X., Barchini, J., Ledesma, H.A., Koren, D., Jin, Y., Liu, X., Wei, W., and Cang, J. (2017). Retinal origin of direction selectivity in the superior colliculus. Nature Neuroscience 20, 550–558. 10.1038/nn.4498.

79. Baden, T., Berens, P., Franke, K., Roman Roson, M., Bethge, M., and Euler, T. (2016). The functional diversity of retinal ganglion cells in the mouse. Nature 529, 345–350. 10.1038/nature16468.

80. Román Rosón, M., Bauer, Y., Kotkat, A.H., Berens, P., Euler, T., and Busse, L. (2019). Mouse dLGN Receives Functional Input from a Diverse Population of Retinal Ganglion Cells with Limited Convergence. Neuron 102, 462–476.e468. 10.1016/j.neuron.2019.01.040.

81. Spacek, M.A., Crombie, D., Bauer, Y., Born, G., Liu, X., Katzner, S., and Busse, L. (2022). Robust effects of corticothalamic feedback and behavioral state on movie responses in mouse dLGN. eLife 11, 1–32. 10.7554/eLife.70469.

82. Parker, P.R.L., Abe, E.T.T., Leonard, E.S.P., Martins, D.M., and Niell, C.M. (2022). Joint coding of visual input and eye/head position in V1 of freely moving mice. Neuron 110, 3897–3906.e3895. 10.1016/j.neuron.2022.08.029.

83. DePiero, V.J., Deng, Z., Chen, C., Savier, E.L., Chen, H., Wei, W., and Cang, J. (2024). Transformation of Motion Pattern Selectivity from Retina to Superior Colliculus. Journal of Neuroscience 44, 1–14. 10.1523/JNEUROSCI.1704-23.2024.

84. Liang, L., and Chen, C. (2020). Organization, Function, and Development of the Mouse Retinogeniculate Synapse. Annual Review of Vision Science 6, 261–285. 10.1146/annurev-vision-121219-081753.

85. Sherman, S.M., and Guillery, R.W. (1996). Functional organization of thalamocortical relays. Journal of Neurophysiology 76, 1367–1395. 10.1152/jn.1996.76.3.1367.

86. Sherman, S.M., and Guillery, R.W. (1998). On the actions that one nerve cell can have on another: Distinguishing "drivers" from "modulators". Proceedings of the National Academy of Sciences of the United States of America 95, 7121–7126. 10.1073/pnas.95.12.7121.

87. Briggs, F., and Usrey, W.M. (2008). Emerging views of corticothalamic function. Current Opinion in Neurobiology 18, 403–407. 10.1016/j.conb.2008.09.002.

88. Sibille, J., Gehr, C., Benichov, J.I., Balasubramanian, H., Teh, K.L., Lupashina, T., Vallentin, D., and Kremkow, J. (2022). High-density electrode recordings reveal strong and specific connections between retinal ganglion cells and midbrain neurons. Nat Commun 13, 5218. 10.1038/s41467-022-32775-2.

89. Sokhadze, G., Whyland, K.L., Bickford, M.E., and Guido, W. (2022). The organization of cholinergic projections in the visual thalamus of the mouse. J Comp Neurol 530, 1081–1098. 10.1002/cne.25235.

90. Lee, S.-H., Kwan, A.C., and Dan, Y. (2014). Interneuron subtypes and orientation tuning. Nature 508, E1–E2. 10.1038/nature13128.

91. Murphy, B.K., and Miller, K.D. (2003). Multiplicative Gain Changes Are Induced by Excitation or Inhibition Alone. The Journal of Neuroscience 23, 1–12.

92. Huberman, A.D., Feller, M.B., and Chapman, B. (2008). Mechanisms underlying development of visual maps and receptive fields. Annual Review of Neuroscience 31, 479–509. 10.1146/annurev.neuro.31.060407.125533.

93. Dhande, O.S., Hua, E.W., Guh, E., Yeh, J., Bhatt, S., Zhang, Y., Ruthazer, E.S., Feller, M.B., and Crair, M.C. (2011). Development of single retinofugal axon arbors in normal and β2 knock- out mice. Journal of Neuroscience 31, 3384–3399. 10.1523/JNEUROSCI.4899-10.2011.

94. Thompson, A., Gribizis, A., Chen, C., and Crair, M.C. (2017). Activity-dependent development of visual receptive fields. Current Opinion in Neurobiology 42, 136–143. 10.1016/j.conb.2016.12.007.

95. Gribizis, A., Ge, X., Daigle, T.L., Ackman, J.B., Zeng, H., Lee, D., and Crair, M.C. (2019). Visual Cortex Gains Independence from Peripheral Drive before Eye Opening. Neuron 104, 711–723.e713. 10.1016/j.neuron.2019.08.015.

96. Matsumoto, N., Barson, D., Liang, L., and Crair, M.C. (2024). Hebbian instruction of axonal connectivity by endogenous correlated spontaneous activity. Science (New York, N.Y.) 385, eadh7814. 10.1126/science.adh7814.

97. Hong, Y.K., Park, S.H., Litvina, E.Y., Morales, J., Sanes, J.R., and Chen, C. (2014). Refinement of the Retinogeniculate Synapse by Bouton Clustering. Neuron 84, 332–339. 10.1016/j.neuron.2014.08.059.

98. Brainard, D.H. (1997). The Psychophysics Toolbox. Spatial vision 10, 433–436.

99. Niell, C.M., and Stryker, M.P. (2008). Highly selective receptive fields in mouse visual cortex. Journal of Neuroscience 28, 7520–7536. 10.1523/JNEUROSCI.0623-08.2008.

100. Barchini, J., Shi, X., Chen, H., and Cang, J. (2018). Bidirectional encoding of motion contrast in the mouse superior colliculus. Elife 7. 10.7554/eLife.35261.

101. Horii-Hayashi, N., and Nishi, M. (2021). Protocol for behavioral tests using chemogenetically manipulated mice. STAR Protocols 2, 100418. 10.1016/j.xpro.2021.100418.

102. Bonin, V., Histed, M.H., Yurgenson, S., and Clay Reid, R. (2011). Local diversity and fine-scale organization of receptive fields in mouse visual cortex. Journal of Neuroscience 31, 18506–18521. 10.1523/JNEUROSCI.2974-11.2011.

103. Burgess, C.R., Ramesh, R.N., Sugden, A.U., Levandowski, K.M., Minnig, M.A., Fenselau, H., Lowell, B.B., and Andermann, M.L. (2016). Hunger-Dependent Enhancement of Food Cue Responses in Mouse Postrhinal Cortex and Lateral Amygdala. Neuron 91, 1154–1169. 10.1016/j.neuron.2016.07.032.

104. Bao, Y., Soltanian-Zadeh, S., Farsiu, S., and Gong, Y. (2021). Segmentation of Neurons from Fluorescence Calcium Recordings Beyond Real-time. Nat Mach Intell 3, 590–600. 10.1038/s42256-021-00342-x.

105. Schröder, S., Steinmetz, N.A., Krumin, M., Pachitariu, M., Rizzi, M., Lagnado, L., Harris, K.D., and Carandini, M. (2020). Arousal Modulates Retinal Output. Neuron 107, 487–495.e489. 10.1016/j.neuron.2020.04.026.

106. Kerlin, A.M., Andermann, M.L., Berezovskii, V.K., and Reid, R.C. (2010). Broadly Tuned Response Properties of Diverse Inhibitory Neuron Subtypes in Mouse Visual Cortex. Neuron 67, 858–871. 10.1016/j.neuron.2010.08.002.

107. Sun, W., Tan, Z., Mensh, B.D., and Ji, N. (2016). Thalamus provides layer 4 of primary visual cortex with orientation- and direction-tuned inputs. Nature Neuroscience 19, 308–315. 10.1038/nn.4196.

108. Oommen, B.S., and Stahl, J.S. (2008). Eye orientation during static tilts and its relationship to spontaneous head pitch in the laboratory mouse. Brain Research 1193, 57–66. 10.1016/j.brainres.2007.11.053.

109. Mathis, A., Mamidanna, P., Cury, K.M., Abe, T., Murthy, V.N., Mathis, M.W., and Bethge, M. (2018). DeepLabCut: markerless pose estimation of user-defined body parts with deep learning. Nature Neuroscience 21, 1281–1289. 10.1038/s41593-018-0209-y.

110. Bewick, V., Cheek, L., and Ball, J. (2004). Statistics review 8: Qualitative data - Tests of association. Critical Care 8, 46–53. 10.1186/cc2428.

111. Ferguson, K.A., Salameh, J., Alba, C., Selwyn, H., Barnes, C., Lohani, S., and Cardin, J.A. (2023). VIP interneurons regulate cortical size tuning and visual perception. Cell Reports 42, 113088. 10.1016/j.celrep.2023.113088.

